# GPCR-targeted imaging and manipulation of homeostatic microglia in living systems

**DOI:** 10.1101/2025.04.01.646501

**Authors:** Swathi Vanaja Chandrasekharan, Payal Chaubey, Sunny Khandelwal, Raghavender Medishetti, Sushreeta Chakraborty, Shruthi Rajaram, Kavya Jayakumar, Pikaso Latua, Rythem Goyal, R Gopalakrishna, Sougat Das, Kiranam Chatti, Kalyaneswar Mandal, Aneesh Tazhe Veetil

## Abstract

Microglia, the resident innate immune cells of the brain, are known to perform key roles such as synaptic pruning, apoptotic debris removal, and pathogen defense in the central nervous system. Microglial mutations are directly linked to many neurodevelopmental (e.g., schizophrenia) and neurodegenerative (e.g., Alzheimer’s disease) disorders, indicating the diagnostic and therapeutic potential of microglia for treating these conditions. Currently, we lack robust molecular tools to specifically image and manipulate microglia *in vivo*, which presents a major hurdle in our understanding of the brain-wide functions of these cells during the early onset of brain diseases. Here, we describe a molecular technology for imaging and manipulation of homeostatic microglia in live organisms (e.g., in zebrafish and mice) by covalently targeting the purinergic receptor, P2RY12. Using this technology, we imaged microglia-pathogen interactions in the larval zebrafish brain and revealed various morphological states of microglia in the adult mouse brain. We further expanded the microglia labelling approach to single-microglia tracking and microglial surfaceome mapping using photoactivatable fluorophores and photoproximity labelling, respectively. We anticipate the use of this universal tool for studying microglial biology across species to reveal the dynamics and polarization of resting microglia into a reactive state found in many neurodegenerative diseases.

## Introduction

Microglia are the resident macrophages of the brain that play a central role in brain development, neuronal circuit remodelling and in neurodegeneration^1–3^. They perform housekeeping functions such as apoptotic debris removal and synapse elimination during early stages of development^4–6^. To achieve this, microglia sense their microenvironment using a collection of receptor proteins known as the ‘sensome’ of microglia, expressed on their cell surface^7^. Our current understanding of microglial biology is primarily gained through the use of genetic tools that enable their visualization and manipulation in *in vivo* settings. For example, CX3CR1-GFP mice, which express green fluorescent protein (GFP) in microglia, have been instrumental in studying microglial biology during brain development and in neurodegenerative diseases^8^. More recently, two additional microglia-specific transgenic lines, Hexb^tdTomato^ and *P2ry12-CreER*, have been described^9,10^. These transgenic tools targeting microglia have been employed to reveal the developmental dynamics of microglia and their phenotypic polarization in various mouse models of neurodegenerative diseases. Although, the transgenic lines are powerful tools for manipulating microglia, they are mostly limited to mice, and recent reports caution against using such lines (e.g., CX3cr1^CreER^) for studying developing microglia, as microglial development has been perturbed due to DNA-damage induced neuroinflammation in those transgenic mice^11^.

New approaches to target microglia based on lentivirus and engineered adeno-associated virus (AAV) platforms were also described^12–18^. Such genetic tools are utilized to target microglia in specific brain regions (e.g., striatum or cortex). AAVs used in these studies are not microglia-specific, and they non-specifically target astrocytes and neurons. This problem was partly circumvented by co-expressing microRNA constructs (e.g., *miR9, miR124*), which suppress non-specific expression of the target genes in neurons^16,18^. Moreover, viral-based genetic cargo delivery tools are known to activate microglia and trigger neuroinflammation, as microglia possess a complete cellular machinery to sense and respond to viruses^19,20^. Microglia-specific, non-activating AAVs have yet to be discovered. Another issue with AAV plasmids is their limited capacity for genetic cargo incorporation; they work well when the cargo load is less than 4.5 kb in size.

Non-viral methods to target microglia would be ideal for studying the functions of homeostatic microglia in *in vivo* settings. Additionally, if such a method can be used to target microglia across species, it would be instrumental for gaining new insights into microglial biology in an evolutionary context. There are a number of studies that describe the activated microglial phenotypes present in diseases such as Alzheimer’s, Multiple Sclerosis (MS), Huntington’s disease, stroke models, etc.^21–25^. Studies that delve into the investigation of homeostatic microglia are limited largely due to the paucity of tools that can investigate the resting phenotype without activating immune signaling pathways. A general non-viral method that can label microglia fluorescently in a silent manner, without activating the cells, would be highly desirable for imaging microglia in various animal models, and in live primary brain tissue samples, where genetic manipulations are ineffective.

Here, we aim to develop a new tool for the live imaging and modulation of microglia without priming them into an activated state. To this end, we first scanned the previously reported microglial sensome proteins and their mRNA expression levels^7^. We searched for marker proteins that are exclusive to microglia and not expressed by astrocytes, neurons, or peripheral macrophages. We found that P2RY12 is one of the highly expressed proteins by microglia with undetectable levels of expression in astrocytes and neurons^7,26,27^. P2RY12 is a metabotropic G-protein-coupled receptor (GPCR) that senses nucleotides such as ADP and ATP present in the brain parenchyma^28,29^. Previous work has shown the importance of ADP and ATP sensing by microglia in responding to brain injury and apoptosis^29,30^. Microglia sense ATP gradients and undergo chemotaxis toward the sites of injury to engulf cellular debris through phagocytosis to mitigate the spread of neuroinflammation^29^. Recent work has demonstrated the critical signaling roles of microglial P2RY12 receptors in orchestrating microglia-neuron communications and proper neuronal connectivity in mouse models^31–33^.

Given the high levels of expression of P2RY12 receptors in microglia and their diverse and emerging biology in neuron-microglia crosstalk in health and disease, we planned to develop a chemical method for imaging homeostatic microglia by targeting the P2RY12 receptor. To this end, we explored the potential of P2RY12 antagonists as microglia imaging agents. Several synthetic P2RY12 antagonists have been previously reported, and some of them are FDA-approved as anti-platelet medications (e.g., clopidogrel) to target the P2RY12 receptor present on platelets and to prevent their aggregation and blood clotting^34^. Next, we selected molecules from the piperazinyl-glutamate-pyrimidine-based P2RY12 antagonist series^35,36^ through molecular docking studies as initial candidates for the development of fluorescent microglial imaging probes. Using a piperazinyl-glutamate-pyrimidine P2RY12 antagonist and adopting ligand-directed protein labelling strategies reported for visualizing neurotransmitter receptors^37,38^ and GPCRs^39^, we have designed and synthesized a series of covalent fluorescent probes for imaging homeostatic microglia. We refer to these probes as **MITIGATE** (<u>M</u>icroglial Imaging <u>T</u>hrough Intrinsic <u>G</u>PCR-<u>A</u>ssisted <u>T</u>ethering of <u>E</u>xogenous molecules), which enable non-perturbative, covalent attachment of fluorophores and other molecular handles to microglial-P2RY12 receptors, thus allowing their long-term visualization *in vitro*, in larval zebrafish (*Danio rerio*), and in adult mouse brains. Using **MITIGATE** probes, we demonstrated two proof-of-concept applications: (i) single-microglia tracking in a population of cell types and (ii) photoproximity labelling of microglial surface proteins.

## Results

### *MITIGATE^C^* covalently labels homeostatic microglia marker protein, P2RY12 receptor

To image homeostatic microglia in real time, we have employed the ligand-directed protein labelling approach to label the P2RY12 receptor with a far-red fluorophore (CY5 or A647) ^40^. To this end, we have selected a piperazinyl-glutamate-pyrimidine antagonist with a reported affinity of 2.3 nM to P2RY12. Our initial molecular docking studies indicated that ligands in the piperazinyl-glutamate-pyrimidine series are amenable to chemical modifications without affecting their binding properties to P2RY12. Next, we have synthesized a P2RY12 ligand with a triethylene glycol linker attached with an azide handle for further modification with fluorophores such as DBCO-cyanine-5 and DBCO-Alexa647 using strain-promoted click chemistry (Figure.1A). We have initially synthesized two molecules in this series: (i) the probe with the covalent reactive centre (**MITIGATE^C^**) and the nonreactive probe (**MITIGATE^NC^**) (Figure 1A). The chemical synthesis and characterization of the probes are described in the SI methods (Schemes 1, 2). The reactive **MITIGATE^C^** probe essentially contains three modules, the first module is the P2RY12 ligand (green, Figure 1A), an acylsulfonamide reactive centre (blue, Figure 1A) for the nucleophilic attack of P2RY12 side-chain residues, and a click chemistry handle (magenta, Figure 1A) for facile functionalization with DBCO-fluorophore conjugates. The non-covalent probe (**MITIGATE^NC^**) lacks the acylsulfonamide reactive centre, and it cannot form a covalent bond with the P2RY12 protein.

**Fig. 1.**
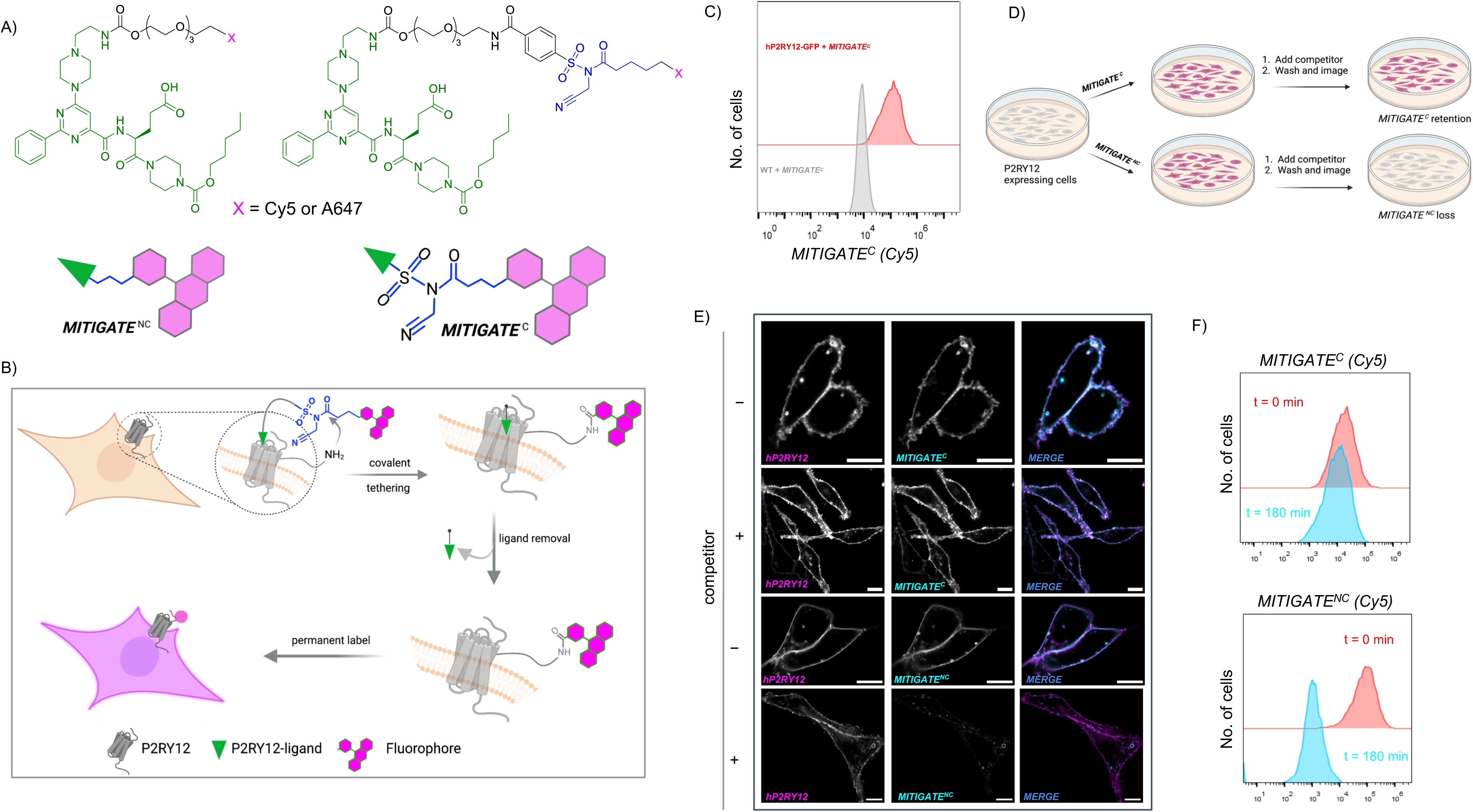
Design, development, and validation of the MITIGATE probes. A) Chemical structures of **MITIGATE**^NC^ and **MITIGATE**^C^ probes B) Schematic showing **MITIGATE^C^**-driven covalent labelling of the P2RY12 receptor C) Histograms of the flow cytometry readouts of **MITIGATE**^C(Cy5)^ signal from wild type CHO and CHO cells expressing hP2RY12-GFP D) Schematic depicting the competition experiments using 250 nM **MITIGATE**^NC^ and **MITIGATE**^C^-labelled hP2RY12-GFP expressing CHO cells in the presence of an excess of P2RY12 antagonist E) Confocal images showing the retention of MITIGATE^C^ signal (cyan) upon competition with P2RY12 antagonist. MITIGATE^NC^ signal was lost upon competition (lower panel, cyan) F) Flow cytometry readouts showing retention of MITIGATE^C^ signal on cells at 0 h (red) and 3 hours (cyan) after the addition of competitor ligand. **MITIGATE**^NC^-treated cells showed dramatic loss of signal after 3 h of incubation with the competitor (cyan). Scale bar: 10 µm. All the results are reproduced in three independent replicates.

The mechanism behind the ligand-directed chemistry is depicted in Figure 1B. In a first step, the **MITIGATE^C^** probe binds to the P2RY12 protein through the interaction with the ligand (green arrowhead). Upon **MITIGATE^C^** binding to the P2RY12, side-chain amino acids (e.g., lysine) can nucleophilically attack the acylsulfonamide reactive centre to result in an amide bond formation by permanently attaching the fluorophore to the P2RY12. Subsequently, the ligand detaches from the protein, leaving behind the fluorophore attached to P2RY12 (Figure 1B). To test the performance of the **MITIGATE** probes, we have added 250 nM of the probe to CHO cells stably expressing human P2RY12-GFP constructs. After incubating the **MITIGATE^C^** probe for one hour and washing, the cells were subjected to flow cytometry based on the signal in the **MITIGATE^C^** channel (Cy5). We have noticed a clear shift (10-fold) in signal in the Cy5 channel due to the covalent attachment of **MITIGATE^C^** probes to the cells (Figure 1C).

Next, we have imaged the **MITIGATE**^C^ and **MITIGATE**^NC^ treated cells to check the probe localization on cells in presence and absence of excess P2RY12 antagonist (for chemical structure, see Figure S1) as a competitor (Figure 1E). In the absence of a P2RY12 competitor, we observed both **MITIGATE**^C^ and **MITIGATE**^NC^ equally labelling the P2RY12-GFP receptor on the plasma membrane of the cells using confocal microscopy (Figure 1E). To check the covalent nature of the bonding of **MITIGATE**^C^ with the P2RY12 receptor, we have introduced excess of the unlabelled P2RY12 antagonist (4x) to the cell culture medium to allow competition (Figure 1D). In the presence of the competitor, **MITIGATE**^NC^ signal was lost from the cell surface, whereas **MITIGATE**^C^ signal was unchanged (Figure 1E, middle panel). The competition was further demonstrated using flow cytometry experiments (Figure 1F) using **MITIGATE**^C^ and **MITIGATE**^NC^ probes, where, 180 minutes of P2RY12 competitor addition resulted in the complete loss of signal from the non-covalent **MITIGATE^NC^** probe treated cells (Figure 1F, bottom panel, cyan). The kinetics of competition was assessed by incubating **MITIGATE^C^** and **MITIGATE**^NC^ probe at different time intervals and the quantification of the data revealed a 90% loss of signal from non-covalent **MITIGATE**^NC^ treated cells within two hours of the experiment. But, the **MITIGATE^C^** signal remained unchanged throughout the experimental time window of 3 hours(Figure S3), indicating the versatility of covalent-tethering approach for long-term imaging studies. We detected **MITIGATE^C^** signal from the cells using as low as 1.5 nM of the probe (Figure S4).

### *MITIGATE^C^* labelling is P2RY12-specific and is non-perturbative

Next, to establish the specific targeting of P2RY12 receptor with the **MITIGATE**^C-A647^ probe, we have treated CHO cells expressing hP2RY12-GFP with 10 μM of **MITIGATE**^C-A647^ (Figure 2A) probe for one hour. We then performed an immuno-precipitation (IP) experiment using *anti*-P2RY12 antibody (Figure 2B & 2C), and the enriched protein fraction was visualized using an *anti*-A647 antibody in a western blot. The solubilized protein fraction showed a prominent A647 emission peak over the untreated control (Figure S5). In the western blot, a band corresponding to the hP2RY12-GFP protein (66.5 kDa, Figure 2C, red arrowhead) was specifically enriched in the **MITIGATE**^C-A647^ treated cells (lane 3), whereas it was absent in the WT CHO cells (lane 1) and in untreated hP2RY12 CHO cells (lane 2), demonstrating the specificity of **MITIGATE^C^**probe to the P2RY12 receptor. Next, we checked whether the covalent attachment of the probe has any effect on ATP/ADP induced P2RY12 receptor signalling. To this end, we performed calcium imaging experiments in hP2RY12 CHO cells that were pre-labelled with **MITIGATE**^C(Cy5)^ probe (Figures 2D-2F). ADP, the natural ligand of P2RY12 receptor, is known to induce cytosolic calcium elevations to initiate chemotactic signalling responses within microglia. **MITIGATE**^C(Cy5)^ labelled CHO-P2RY12 cells (Figure 2E) were loaded with Rhod-4-AM ester dye to monitor ATP/ADP-induced calcium elevations. **MITIGATE**^C(Cy5)^ labelled cells showed robust calcium spikes upon ATP addition (Figure 2F, bottom panel, SI Videos 1 & 2). The spike amplitudes and the time of spiking were comparable in both treated and untreated cells (Figures 2G & 2H), indicating that the ADP binding pocket in P2RY12 is unobstructed by **MITIGATE**^C(Cy5)^ probe labelling. This is particularly important when studying homeostatic microglial signalling in an *in vivo* context.

**Fig. 2.**
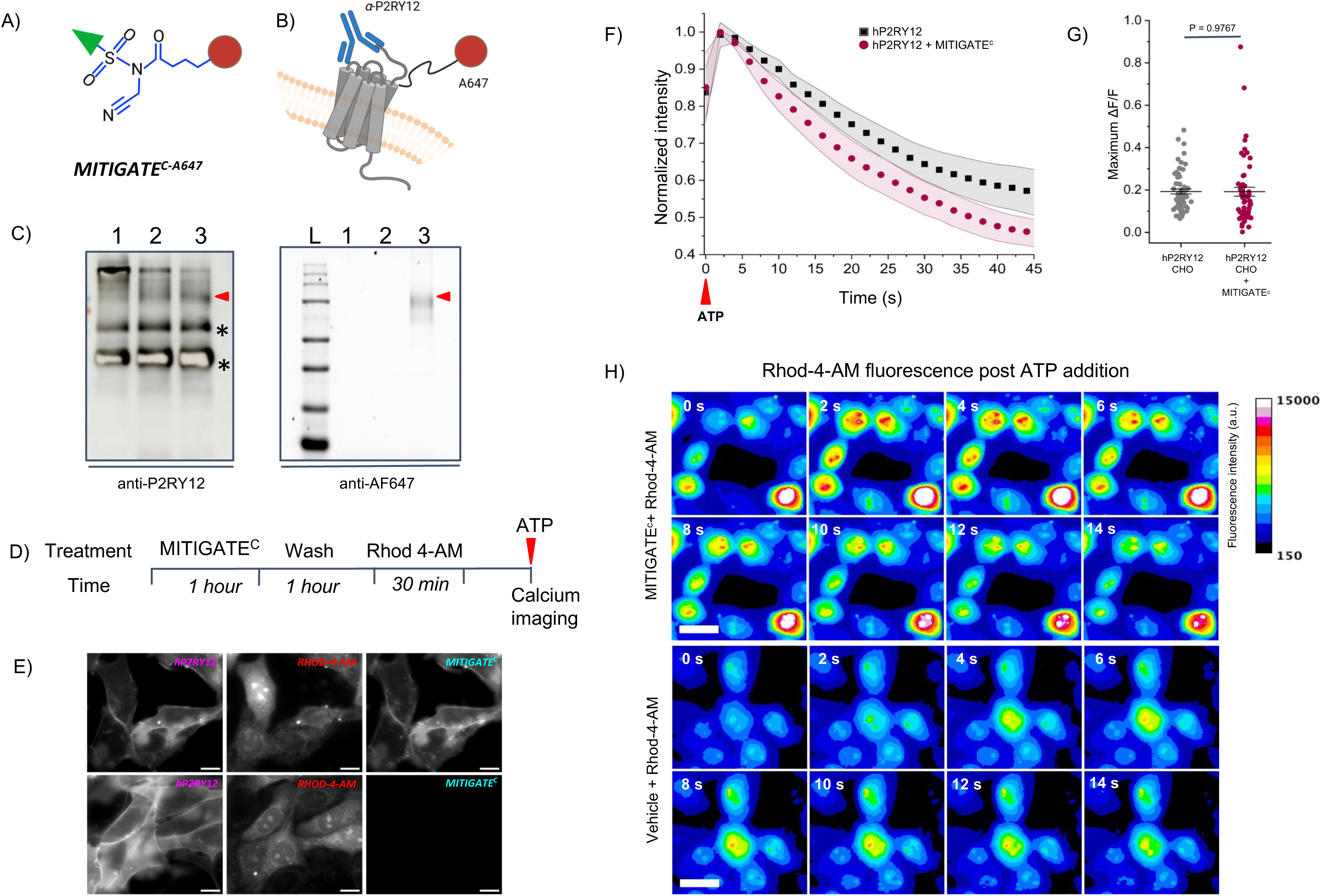
MITIGATE^C^ specifically targets the P2RY12-receptor in a non-perturbative manner. A) A cartoon showing the general structure of **MITIGATE**^c^-A647. Green arrowhead represents the P2RY12 ligand, red circle and blue represents A647 fluorophore and the reactive centre B) Schematic showing the immunoprecipitation (IP) experiment of A647-labelled hP2RY12⎼ GFP receptors using *anti-*P2RY12 antibody. (C) Western blots showing the expected hP2RY12-GFP (∼ 66.5 kDa) band in the A647 channel. The asterisk denotes non-specific bands observed when using the *anti-*P2RY12 antibody. D) Experimental design of P2RY12-activated calcium sensing using **MITIGATE**^C(CY5)^ -labelled cells E) widefield images showing the loading of Rhod-4-AM (red, 𝛌_ex_= 561 nm) dye in hP2RY12-GFP (magenta, 𝛌_ex_= 488 nm) expressing, **MITIGATE**^C(CY5)^-prelabelled (top) and unlabelled cells (bottom, cyan, 𝛌_ex_= 647 nm) F) Time traces of intracellular Ca^2+^ spikes observed upon addition of 300 µM of ATP is plotted as the fluorescence ratio (F/F_0_) of Rhod-4-AM signal (red, 𝛌_ex_= 540 nm). G) Mean amplitude of the ATP-induced Ca^2+^ signal is plotted for **MITIGATE**^C(CY5)^ -labelled and unlabelled cells (n = 60 cells in each group) H) Montage showing the ATP-induced calcium heatmaps of **MITIGATE**^C(CY5)^ -labelled and unlabelled cells. Scale bar: 20 µm. All the results are reproduced in three independent replicates.

### *MITIGATE^C^* revealed primary microglial morphology in culture

We next explored the potential of **MITIGATE**^C(Cy5)^ probe in labelling the P2RY12 receptor in BV2-microglia and in primary mouse microglia. Since the P2RY12 ligand used in the construction of **MITIGATE**^C(Cy5)^ had not been tested against mouse P2RY12 previously, we first checked the ligand interaction sites within the mouse P2RY12 (mP2RY12) receptor using structural alignment. We found that most of the amino acid residues involved in ligand binding are conserved between hP2RY12 and mP2RY12 (Figure 3A). Next, to validate the labelling of mP2RY12 with the **MITIGATE**^C(Cy5)^ probe, we cloned mP2RY12-GFP and expressed it in HEK293T cells (Figure 3B). We found localization of the mP2RY12-GFP (magenta) to the plasma membrane, and it was fluorescently labelled specifically by treating the cells with 250 nM of **MITIGATE**^C(Cy5)^ probe (Figure 3B, cyan). We observed a consistent 10-fold increase in **MITIGATE**^C(Cy5)^ fluorescence signal from mP2RY12 expressing cells compared to the WT-HEK293T cells when treated with **MITIGATE**^C(Cy5)^ and analysed using flow cytometry (Figure 3D). We next attempted labelling of mP2RY12 in mouse BV2 microglial cells and BV2 cells overexpressing mP2RY12-GFP. We observed efficient labelling of the mP2RY12 receptor in BV2 cells using **MITIGATE**^C(Cy5)^ (cyan) probe, and the signal colocalized with the mP2RY12-GFP (magenta) signal on the plasma membrane (Figure 3C). **MITIGATE**^C(Cy5)^ revealed the preferential enrichment of the P2RY12 receptors at the filopodia of the BV2 cells (orange arrowhead, Figure 3C). The localization of the P2RY12 receptor at the filopodia of motile microglia has been previously observed in primary microglial cells^41^. The expression of P2RY12 is known to downregulate during the immortalization of the mouse microglial cell line (BV2 cells). Wild-type BV2 cells are known to express only basal levels of endogenous P2RY12 receptors on their plasma membrane. Using the **MITIGATE**^C(Cy5)^ probe, we revealed the endogenous expression of P2RY12 in WT-BV2 cells, demonstrating the potential use of the probe to sort low P2RY12 expressing cells (Figure 3E).

**Fig. 3.**
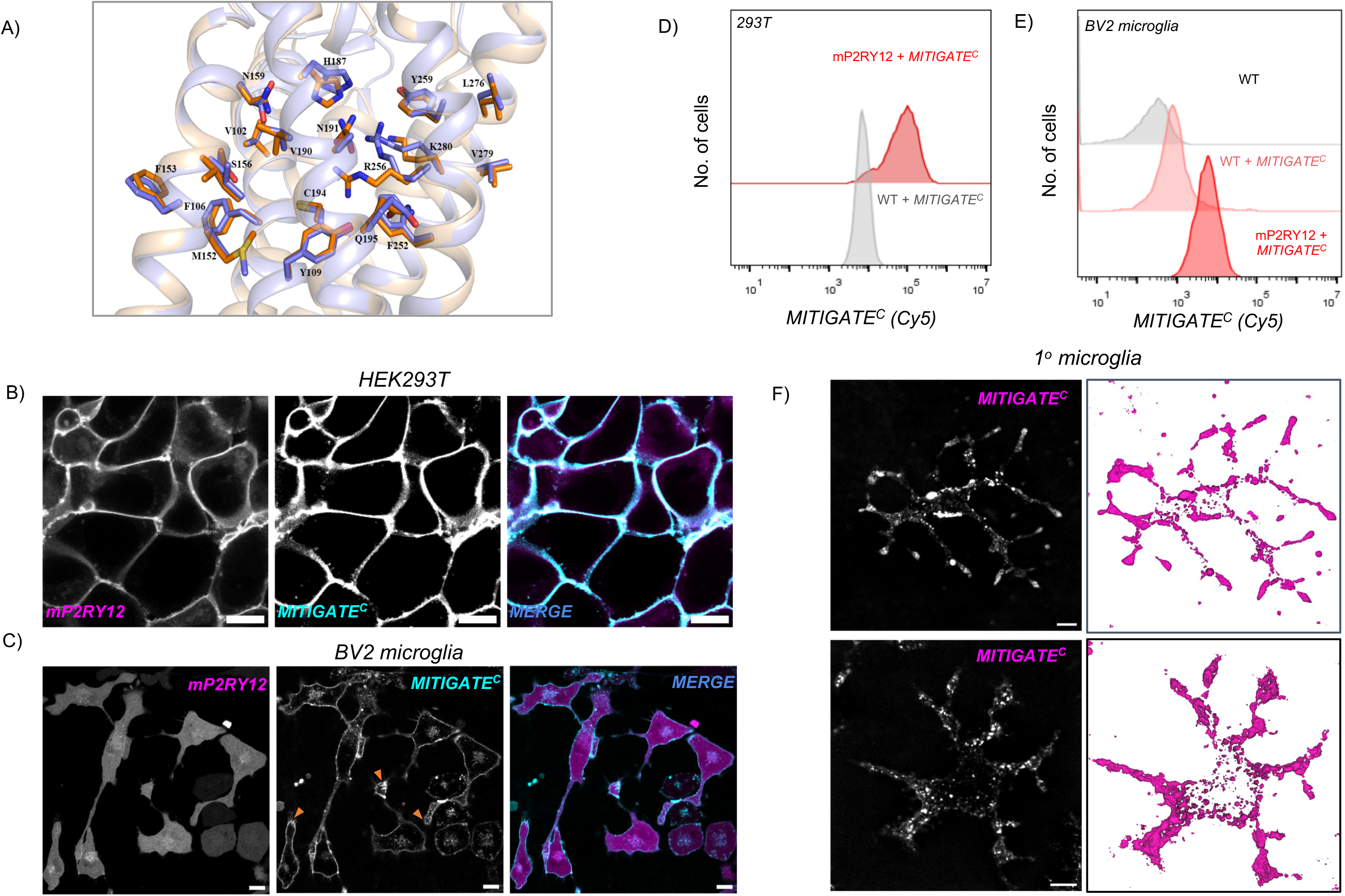
MITIGATE^C^ aids visualization of primary microglia in culture. A) Structural alignment of hP2RY12 receptor; (PDB ID: 4PXZ (wheat) with the AlphaFold-predicted mP2RY12 receptor (light blue, cartoon representation). Conserved residues interacting with P2RY12-antagonist used in our study are highlighted in stick representation. B) Confocal images showing colocalization of the mP2RY12-GFP (magenta, 𝛌_ex_= 488 nm) with **MITIGATE**^C(Cy5)^ (cyan, 𝛌_ex_= 647 nm) when expressed in HEK293T cells. C) Confocal images showing the colocalization of mP2RY12-GFP (magenta, 𝛌_ex_= 488 nm) with **MITIGATE**^C(Cy5)^ (cyan, 𝛌_ex_= 647 nm) in BV2 cells. Localization of the P2RY12 protein was evident at the tip of the filopodia (orange arrowhead) (D) Flow cytometry readouts for **MITIGATE**^C(Cy5)^ signal obtained from HEK293T-WT and mP2RY12-GFP expressing HEK293T cells. E) Flow cytometry readouts for **MITIGATE**^C(Cy5)^ signal obtained from BV2-WT and BV2 cells expressing mP2RY12-GFP. F) Two ROI’s showing the confocal images of P2RY12 labelling using **MITIGATE**^C(Cy5)^ (magenta, 𝛌_ex_= 647 nm) in cultured primary mouse microglia. ROIs showing the **MITIGATE**^C(Cy5)^ labelling in 3D-rendered primary microglia. Scale bar: 10 µm. All the results are reproduced in three independent replicates.

We next performed experiments on cultured primary microglial cells isolated from mouse pups on postnatal days 3-4 (Figure S6). The isolation of microglia from the brain is known to rapidly downregulate P2RY12 expression in culture^28^. We assessed the endogenous expression of P2RY12 in primary microglia using **MITIGATE**^C(Cy5)^ probe after harvesting and four days in culture. **MITIGATE**^C(Cy5)^ revealed ramified microglial morphology with extended processes (Figure 3F), and preferential localization of the P2RY12 receptor at the tips of the microglial processes was evident (magenta, Fig. 3F, SI Video 3) using confocal imaging. P2RY12 expression at the filopodial terminals is proposed to drive local purinergic signalling during microglial migration in response to ATP, and it acts as a specific interaction zone for contacting other cell types in the brain (e.g. neuronal soma and synapses)^32^.

### *MITIGATE^C^* enables the visualization of microglia in healthy and E. coli-infected larval zebrafish brains

We next explored the potential of **MITIGATE**^C(Cy5)^ probe for *in vivo* microglial imaging using larval zebrafish as a model system. We have first performed protein sequence and structural alignment studies using AlphaFold2-predicted structures of the zebrafish P2RY12 receptor (zP2RY12) (Figure S8A), and verified the identity of amino acids required for the zP2RY12 interaction with the synthetic P2RY12 antagonist used in our design of the **MITIGATE^C^** probe. We then cloned the zP2RY12-Staygold fusion construct and expressed it in HEK293T and CHO cells. **MITIGATE**^C(Cy5)^ probe addition to HEK293T-zP2RY12 cells showed strong plasma membrane localization of the probe (Figure 4A, cyan) and it colocalized with the zP2RY12-Staygold signal (Figure 4A, magenta). We observed a tenfold enhancement of **MITIGATE**^C(Cy5)^ signal in flow cytometry experiments in probe treated cells (Figure 4A, bottom right panel) compared to the WT control. The same trend was also seen in zP2RY12 CHO cells when treated with **MITIGATE**^C(Cy5)^ probe (Figure S8B-C). This promising *in vitro* data motivated us to validate microglial targeting of **MITIGATE**^C(Cy5)^ probe in the developing zebrafish central nervous system (CNS).

**Fig. 4.**
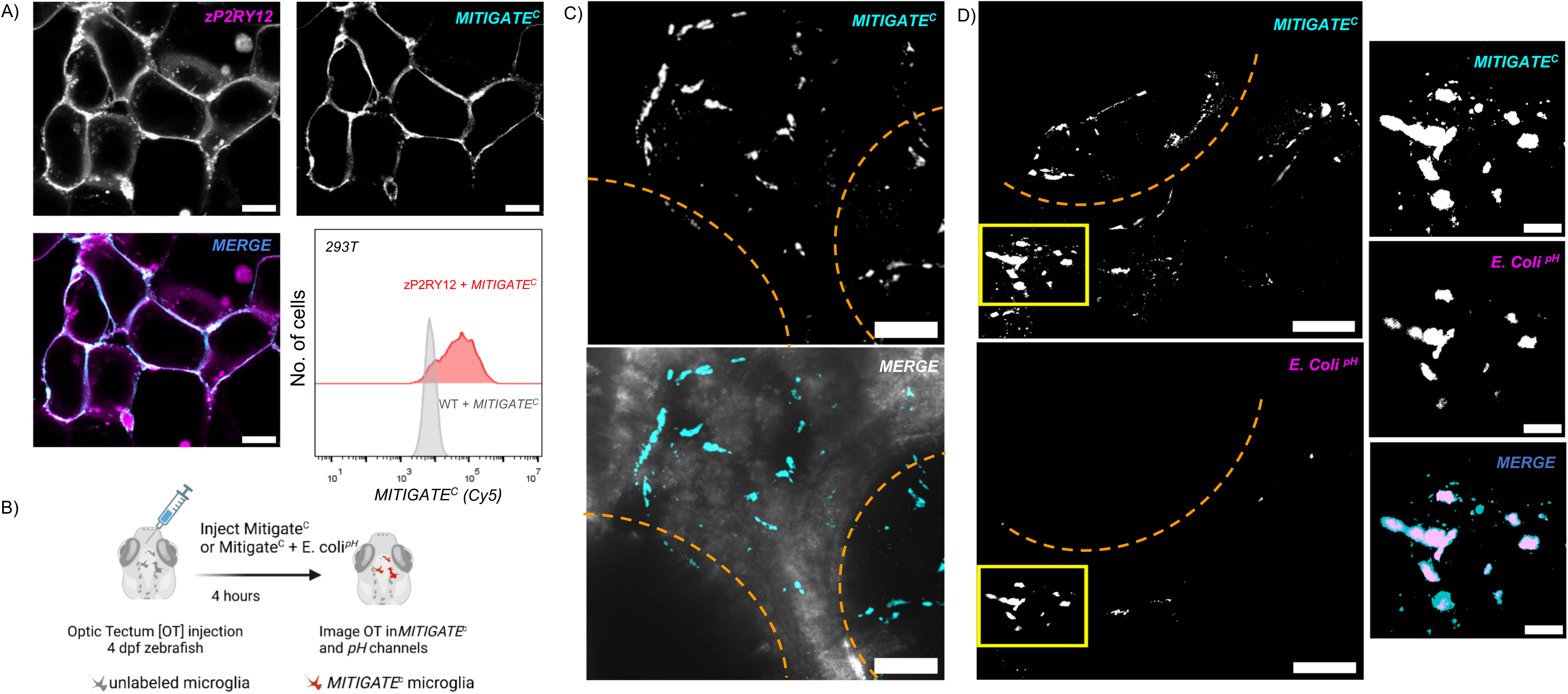
MITIGATE^C(Cy5)^ enables the visualization of microglia in larval zebrafish brains: A) Confocal images showing **MITIGATE**^C(Cy5)^ labelling of zP2RY12-Staygold (magenta, 𝛌_ex_= 488 nm) receptors expressed in HEK293T cells (cyan, 𝛌_ex_= 640 nm). Flow cytometry readouts of **MITIGATE**^C(Cy5)^ labelled zP2RY12-Staygold in HEK293T cells. B) Experimental design showing the microinjection of **MITIGATE^C(Cy5)^**probe into the optic tectum of 4dpf old larval zebrafish C) Confocal images showing the labelling of microglia in **MITIGATE**^C(Cy5)^ injected live zebrafish brain (cyan, 𝛌_ex_= 640 nm) D) Confocal images showing the phagocytosis of injected pHrodo-*E.Coli* particles (magenta, 𝛌_ex_= 514 nm) by **MITIGATE**^C(Cy5)^-labelled microglial cells (cyan) in live zebrafish brain. Inset shows the zoomed images from the boxed ROI. Scale bar: 60 µm. Inset scale bar: 20 µm. All the results are reproduced in three independent replicates.

Larval zebrafish have been an instrumental model in advancing our understanding of microglial biology in brain development. Pioneering studies by Nüsslein-Volhard have revealed the dynamic interplay of microglial cells with neurons and their housekeeping functions such as apoptotic debris removal and pathogen elimination through phagocytosis^4^. Here, we employed 4 dpf (days post fertilization) old larval zebrafish to test the ability of the **MITIGATE**^C(Cy5)^ probe to target microglia in the whole brain. Larval zebrafish at 4 dpf approximately have about 30 developing microglia in their brains, and most of the microglial studies have been conducted in the optic tectum region using this model system (Figure 4B). We injected 3 nL of the **MITIGATE**^C(Cy5)^ probe (10 µM stock) into the optic tectum region of the anesthetized wild-type larval zebrafish and allowed them to recover in the E3 buffer for 4 hours (Figure 4B). Subsequently, we anesthetized the fish in the tricaine-E3 buffer and immobilized them dorsally in an agar bed to perform confocal imaging. The whole-brain imaging of **MITIGATE**^C(Cy5)^ injected fish revealed cells with early developmental microglia morphology, and they were distributed throughout the optic tectum and within the eyes (Figure 4C, SI Video 4). Since there are no zP2RY12 antibodies available to probe microglia in fish, we have conducted an E. coli infection experiment in the brain to monitor E. coli uptake by microglia following the protocol described in the original report by N**ü**sslein-Volhard^4^. Injection of pH-indicator-labelled E. coli (pHrodo E. coli) into the larval zebrafish brain is known to initiate microglial engulfment of the injected particles^4^. This process is solely performed by microglia in the developing zebrafish. We first labelled microglia using **MITIGATE**^C(Cy5)^ and then injected heat-killed pHrodo E. coli particles into the optic tectum of the larval fish, and probed the localization of E. coli to the microglial cells in the brain. Live confocal imaging experiments revealed specific targeting of pHrodo E. coli particles into the **MITIGATE**^C(Cy5)^ labelled cells, thus, proving the microglial identity of those cells (Figure 4D). Three-dimensional (3D) rendering of the cells clearly demonstrated E. coli particles present inside **MITIGATE**^C(Cy5)^ - labelled cells in the brain (SI video 5). Thus, the **MITIGATE**^C(Cy5)^ probe revealed snapshots of microglia-pathogen interactions within the developing larval zebrafish, demonstrating its *in vivo* potential in long-term imaging and studying microglia-pathogen interactions in wild-type zebrafish.

### *MITIGATE^C^* targets homeostatic microglia in adult mouse brain

Imaging of homeostatic microglia and manipulating them *in vivo* is challenging because viral-based transduction of genetic fluorescent reporters (e.g. GFP) is known to induce proinflammatory immune signalling. This is a major bottleneck in understanding the physiological functions of resting microglial phenotypes across species. To expand the potential of the **MITIGATE**^C^ probes for *in vivo* imaging of microglia in mice, we injected the probe into the lateral ventricle (LV) of 4-month-old mouse (Figure 5A). **MITIGATE**^C(A647)^ probe (3 µL, 50 µM) and vehicle (1% DMSO in PBS) were injected into the LV of anesthetized mice, and the animals were left to recover from anesthesia for 18 hours. The brains of **MITIGATE**^C(A647)^ and vehicle injected mice were isolated after transcardial perfusion with PBS and the tissue was fixed using paraformaldehyde. Confocal imaging of sagittal sections showed **MITIGATE**^C(A647)^ probe labelling of microglia-like cells across the brain (Figure 5B, Figure S9). To validate the microglial identity of these cells, we have performed immunofluorescence experiments using microglia marker IBA-1 and astrocyte marker GFAP. We observed a good colocalization between **MITIGATE**^C(A647)^ and IBA-1 antibody (Figure 5C, SI Video 6), whereas the astrocyte marker GFAP showed anti-colocalization (Figure 5C, SI Video7), which established microglia-specific labelling of **MITIGATE**^C(A647)^. Specific targeting of the **MITIGATE**^C(A647)^ probe to microglia further validates the findings of previous transcriptomic studies that showed exclusive expression of the P2RY12 receptor in homeostatic microglia and not by any other cell types in the brain. Using **MITIGATE**^C(A647)^ probe, we observed ramified microglial processes throughout the brain regions, such as the striatum, cerebellum, hippocampus and cortex (Figure S9, SI Video 8). We used a microinjection protocol for the delivery of **MITIGATE**^C(A647)^ probe into the lateral ventricle (see Methods). Microglia are known to respond to local needle injury by transforming their phenotype from a resting state into an activated state. We examined the morphology of microglia present at the distal (> 250 microns away) and at the proximal site of injection (< 250 microns), and observed amoeboid-shaped cells proximal to the injury site (Figure 5D). Microglia present at > 250 microns away from the site of injury showed a clear ramified morphology (Figure 5D). Next, we performed morphometric analysis (Sholl analysis) to quantify the morphological complexity of microglia visualized in **MITIGATE**^C(A647)^ channel (Figure 5E). We observed a 50% reduction in the number of branch intersections on microglia present near the site of injury when compared to the distal microglia. Microglial mean branch lengths were also reduced at the proximal site of injury (Figure 5F). This observation demonstrates the versatility of the **MITIGATE**^C(A647)^ probe in imaging and quantification of both homeostatic and primed microglial phenotypes induced by needle injury in wild-type animals, without the aid of antibody markers.

**Fig. 5.**
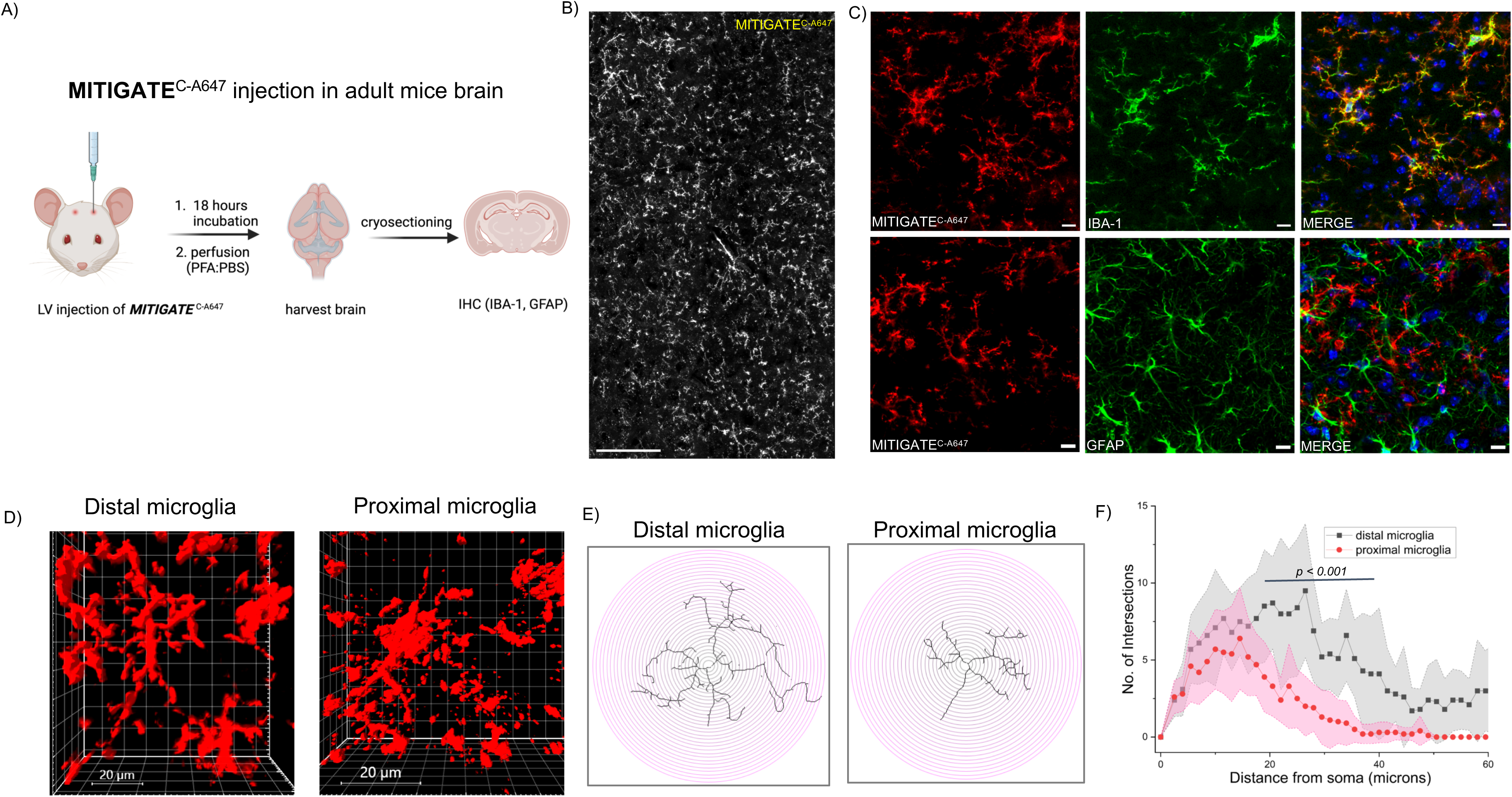
MITIGATE^C-A647^ targets homeostatic microglia in adult mouse brain: A) Schematic showing the experimental design of **MITIGATE**^C-A647^ microinjection into the adult mouse brain B) A large ROI from the midbrain showing uniform labelling of microglia with **MITIGATE**^C-A647^ probe (grey, 𝛌_ex_= 640 nm). Scale bar: 100 µm. C) Images showing colocalization of **MITIGATE**^C-A647^ (red, 𝛌_ex_= 640 nm) with the anti-IBA1 antibody (green, 𝛌_ex_= 546 nm, top panel) and anti-colocalization with the anti-GFAP antibody (green, 𝛌_ex_= 488 nm, bottom panel). Scale bar: 10 µm. D) Imaris 3D-rendering showing morphology of microglial cells away (> 250 microns) and near (< 250 microns) the probe injection site, visualized in the **MITIGATE**^C-A647^ channel E) Morphometric analysis showing skeletonized microglia near and away from the injection site. Sholl analysis was performed with a recurring distance of 1.5 µm from the centre. F) Sholl analysis showing the number of intersections vs distance plot. One-tailed student statistical test is performed. Grey plot represents distal microglia present away from the injection site; Pink plot represents microglia present near the needle injury site. Scale bar: 10 µm. All the results are reproduced in three independent replicates.

### *MITIGATE* offers a platform for single-microglia tracking and proximity labelling of microglial surfaceome

Next, we explored the potential applications of the **MITIGATE**^C^ probes. Recent transcriptomics studies revealed the existence of different microglial cell types displaying a spectrum of phenotypes across the brain^42^. A non-perturbative method to image and modulate microglia at single-cell resolution would be desirable to understand their biology in a brain microenvironment-specific manner. To monitor single microglial dynamics within a population, we have designed a **MITIGATE**^C^ probe with a photoactivatable deep-red fluorophore based on the PA-Janelia Fluor-646 system^43^, which we refer to as **MITIGATE**^PA^ (Figures 6A, 6B & SI Scheme 3). Localized photolysis of **MITIGATE**^PA^ with a 405 nm laser would result in bright fluorescence at 660 nm, enabling single microglia visualization and tracking (SI Video 9). To demonstrate the concept, we cocultured BV2 microglia expressing mP2RY12-GFP receptors with DIV10 hippocampal neurons for a day (Figure S6 and SI Video 10), and the **MITIGATE**^PA^ probe (250 nM) was added to the culture to label all the microglia. We then photoactivated single microglia within a region of interest using a 405 nm laser to mark the cell with Janelia Fluor-646, resulting from the photoactivation of **MITIGATE**^PA^ probe, thus allowing tracking of single microglia at 660 nm (Figure 6B, SI Video 11). This proof-of-concept experiment demonstrated the potential of **MITIGATE**^PA^ probe in revealing single microglia-neuron interactions in co-culture systems in a nonperturbative manner. We anticipate several potential applications of **MITIGATE**^PA^ probe in mapping neuro-immune interactions, particularly in advancing our understanding of human microglial biology using recently developed human iPS-derived microglial (iMG)-neuron co-culture systems^44,45^.

**Fig. 6.**
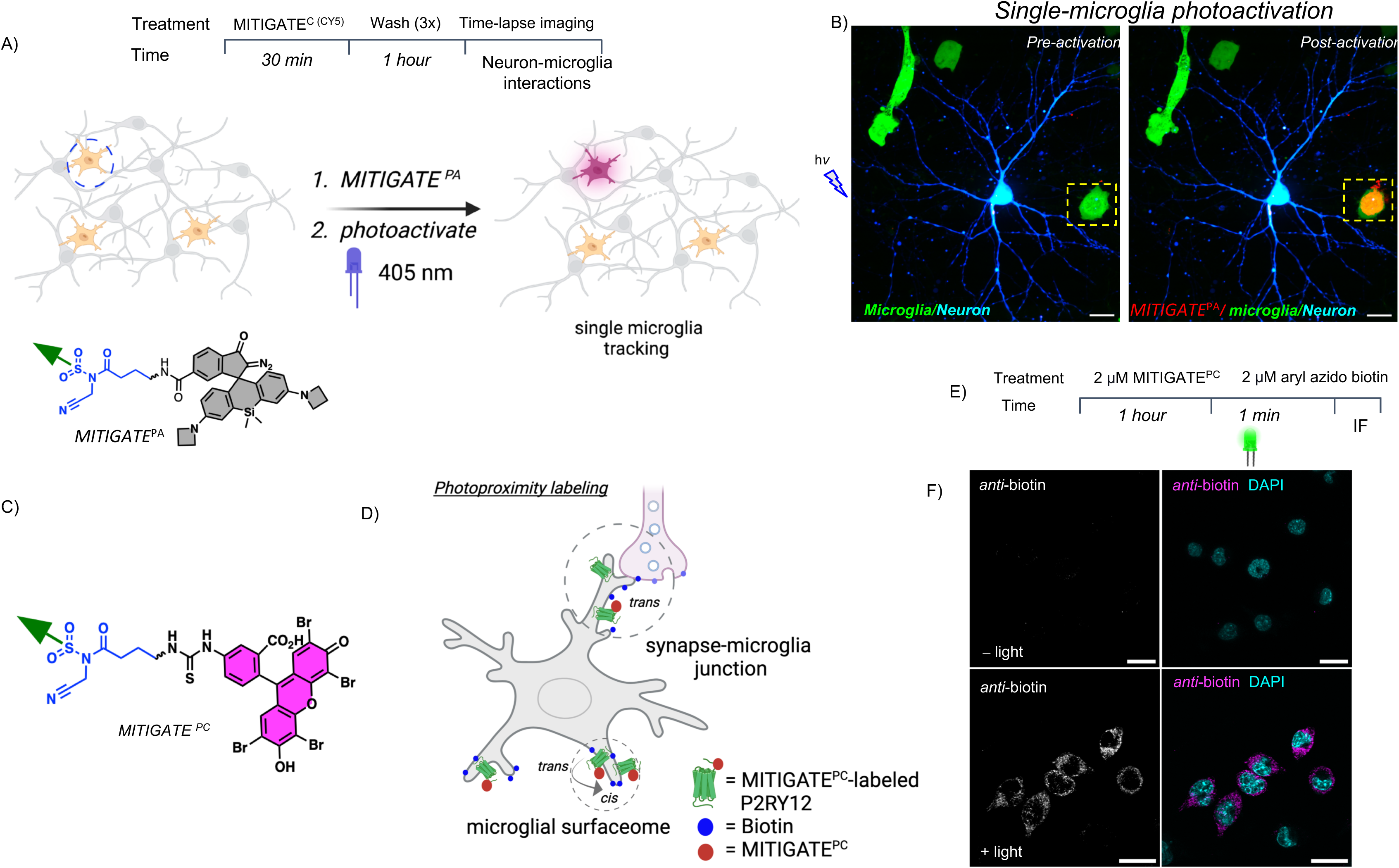
MITIGATE^PA^ and MITIGATE^PC^ probes enable single-microglia tracking and photoproximity labelling of microglial surface proteins. A) A schematic showing the treatment of neuron-BV2 microglia co-culture with **MITIGATE**^PA^. Photoactivation of single-microglia (dotted circle) using blue light (405 nm) produces red fluorescence (675 nm). The chemical structure of **MITIGATE**^PA^ is also shown. B) A confocal image showing hP2RY12-GFP-expressing BV2-microglia (green) before photoactivation in co-culture with BFP expressing neurons (blue). The preselected ROI for photoactivation is indicated inside a dotted square. Upon photoactivation, red fluorescence is generated within the ROI, which is colocalized with the microglia (green). Scale bar: 20 µm C) A schematic depicting the chemical structure of **MITIGATE**^PC^ probe. D) A schematic illustrating the proximity labelling of microglial surface proteins (*cis*-labelling) and proteins on the neurons (*trans*-labeling) with biotin using **MITIGATE**^PC^ -based photoproximity approach. P2RY12 receptor (green) labelled with **MITIGATE**^PC^ probe (red circle) initiates biotin (blue circle) labeling of nearby proteins upon photoactivation with a green LED (532 nm) light E) Schematic showing the experimental design F) Photoproximity labelling of the microglial surfaceome and microglial-contact sites on the neurons revealed by the **MITIGATE**^PC^ probe in microglia-neuron co-cultures. Biotinylated proteins on the microglia and neuronal surfaces are detected using an anti-biotin antibody (magenta). Scale bar: 10 µm. All the results are reproduced in three independent replicates.

Several lines of evidence suggest the existence of subtypes of microglia across brain regions^42,46^. Microglial plasticity, originating from an intrinsic transcriptional state, can lead to unique functional states^42^. The transcriptional and functional differences observed in microglia are strongly dependent on the CNS microenvironments^47,48^. For example, cerebellar microglia in mice are known to display higher phagocytic clearance abilities when compared to striatal microglia in a healthy brain^49^. Microglial subtypes such as keratin sulfate proteoglycan (KSPG)-microglia, Hox8b-microglia, TREM2-microglia etc., are known to enrich only in certain brain regions, and their numbers vary in health versus disease contexts^42^. For instance, TREM2-microglia is enriched in cingulate and entorhinal cortex, and TREM2 influences microglial proliferation, survival, and phagocytosis^50^. The biological significance of these subtypes of microglia in a healthy brain is not well understood. This is because the the identity of proteins governing such microglial subtypes, as well as their interactions with the microenvironment and other cell types in the brain (e.g. neurons), is difficult to study using proteomic tools such as BioID^51^ and Turbo-ID^52^. These tools rely on genetic expression in cells and therefore require delivery through viruses which would prime microglia to an activated proinflammatory state, thus hindering their study in a homeostatic condition. To address this issue, we envisaged the use of a **MITIGATE**^C^ probe to map microglial surface proteins (surfaceome).

A recent study has mapped the EGFR-interactome in cancer cells using proximity-induced photocatalyst approach^53^. Inspired by this study, we set out to map the P2RY12-interactome of microglial cells. To achieve this, we have first synthesized a **MITIGATE**^C^ probe containing a microglial P2RY12-targeting ligand, a reactive centre, and a photocatalyst (Eosin-Y) (Figure 6C, SI Scheme 4), which we call **MITIGATE**^PC^ (Figure 6C). Incubation of 2 µM of **MITIGATE**^PC^ with BV2-microglia has led to the covalent attachment of the photocatalyst specifically to the P2RY12 receptor present on the plasma membrane of microglia (Figure 6D). Following ligand-induced Eosin-Y catalyst immobilization on the microglial P2RY12 receptor, excess **MITIGATE**^PC^ probe was washed away and removed from the medium (Figure 6E).

Subsequently, the cells were treated with 100 µM of aryl tetrafluoroazido biotin substrate and irradiated using a custom-built green LED system (532 nm, 3 mW, 1-2 min). The cells were then washed, fixed and stained for biotinylated proteins using an *anti*-biotin antibody (Figure 6E). Through **MITIGATE**^PC^-induced proximity labelling, we observed extensive biotinylation of microglial proteins (Figure 6F) in the presence of green light. Biotinylation was not detected when the cells were not irradiated (Figure 6F). This proof-of-concept proximity labelling experiment revealed the potential of **MITIGATE**^PC^ in functionalizing microglia for surface protein labelling. Since **MITIGATE**^PC^ labelling of microglial P2RY12 receptor is non-perturbative, it would help to reveal the homeostatic surfaceome of microglia. This is particularly important in studying subtypes of microglia present in brain-specific regions (e.g., KSPG *vs* TREM2 *vs* Hox8b-microglia).

We anticipate that utilizing **MITIGATE**^PC^ alongside proteomic approaches in an *in vivo* setting will reveal microglial subtypes in a context dependent manner (homeostatic *vs* reactive state). This information is difficult to obtain using genetic techniques in microglia. We currently plan to expand the **MITIGATE^PC^**-based photoproximity labelling strategy to iPS-derived neuro-immune models and in *in vivo* systems to reveal molecular players involved in microglial surfaceome and microglia-neuron interactions in microglial subtypes.

## Discussion

Microglial cells are refractory to genetic manipulations due to the rapid phenotypic polarization exhibited by these cells when sensing external stimuli. Here, we demonstrate a platform technology for the imaging and manipulation of microglia based on the ligand-directed labelling of the P2RY12 receptor. We show that the **MITIGATE^C^** probes introduced in this study enable targeted covalent labelling of P2RY12 receptors with various fluorophores and functional chemical handles, thus offering a modular and powerful platform for imaging and manipulation of homeostatic microglia.

The non-covalent **MITIGATE^NC^** is also specific to P2RY12, but it detaches from the cognate receptors within a couple of hours. While working with sensitive cells like microglia, minimal probe dosage is recommended to keep the homeostatic state of the cells preserved. In our experiments, we have achieved labelling of microglia using probe concentrations as low as 1.5 nM in cell culture. We could also capture minor levels of the endogenous P2RY12 receptors expressed by BV2-microglia using **MITIGATE^C^**. Previous studies have described a positive correlation between the resting state of microglia and P2RY12 expression levels^28^. Microglia freshly isolated from the brain lose expression of P2RY12 within ten hours and the expression of the receptor can be regained by culturing in serum-deprived medium^28^. P2RY12 levels revealed by **MITIGATE^C^**imaging can be used as a proxy for microglial activation state under a given paradigm and it offers a reliable method for mapping the transition of resting state microglia to an activated phenotype.

Using **MITIGATE^C^**, we monitored the distribution of P2RY12 receptors on the plasma membrane of BV2 cells and in primary microglia. Particularly, the enrichment of P2RY12 receptors at the filopodia of the cells were evident during imaging. Such enriched zones of P2RY12 at the tips of the filopodia could serve as specialized domains for communication with nearby cells (e.g., astrocytes and neurons) through ATP signaling.

It is crucial to verify the non-perturbative nature of **MITIGATE^C^** -labelling of P2RY12 receptor on microglia because purinergic signaling operating through the ATP-P2RY12 axis is vital for several signaling events that occur during homeostasis and brain injury. P2RY12 knockout in microglia is known to result in diminished branch extensions, polarization, and migration capability (28,29). We have verified the targetability of **MITIGATE^C^** to the P2RY12 receptor using western blot and functional calcium imaging studies. We detected a band corresponding to the P2RY12-GFP fusion protein on the western blot using immunoprecipitation. Additionally, we validated the presence of A647-labelled, solubilized P2RY12 receptors in lysates and quantified it using spectrofluorimetric analysis. Using Rhod-4-AM calcium imaging dye, we observed comparable amplitudes and durations of calcium signals when cells were first labelled with the **MITIGATE^C^** probe and subsequently treated with ATP. This experiment indicated the regeneration of an empty ADP-binding pocket within the P2RY12 upon completion of **MITIGATE^C^** labelling, and confirmed the detachment of the antagonist from the pocket. This is crucial for studying dynamic microglial interactions that operate within microglia and between microglia and other brain cells (e.g., neurons) in brain development.

Next, we demonstrated the potential of **MITIGATE^C^**in imaging microglial cells in developing zebrafish larvae. Microglia in zebrafish are known to respond to ATP produced during injury and they exhibit targeted migration toward the injury site, demonstrating an evolutionarily conserved P2RY12 pathway in vertebrates^54^. Injection of **MITIGATE**^C(CY5)^ probe into the optic tectum of 4 dpf old wild-type zebrafish enabled the visualization of microglia. We also demonstrated the utility of **MITIGATE**^C(CY5)^ in revealing E. coli-microglia interactions and phagocytic engulfment. Future studies will focus on using **MITIGATE**^C(CY5)^ to capture microglial dynamics and polarization in injury models at single-cell resolution.

Next, we have demonstrated the utility of **MITIGATE^C^**in capturing morphologically diverse microglial cells in the mouse brain. Lateral ventricle injection of the probe revealed ramified morphology with numerous processes of microglia in wild-type mice. Morphologically distinct microglial cells were evident throughout the brain. Microglia at the site of injection showed activated morphology with fewer processes due to needle injury-induced inflammation. The image quality of **MITIGATE^C^**-labelled microglia was comparable to a commercial IBA-1 antibody used in the field for imaging microglia and macrophages. P2RY12 expression is specific to microglia, showing no off-target effects from peripheral macrophages; however, IBA-1 also stain for peripheral macrophages. A thorough study of brain-wide **MITIGATE^C^** labelling to reveal all the subtypes of microglia is ongoing in our laboratory. In the future, we anticipate developing a series of **MITIGATE^C^** probes containing photostable fluorophores such as Janelia Fluors to make them multi-photon compatible for deep tissue imaging applications. We also plan to test **MITIGATE^C^**in freshly isolated primary human brain tissue samples to understand microglial dynamics and interactions within. This is particularly important, as it can only be revealed by a fast method like **MITIGATE** owing to the short lifespan of the primary tissue in culture.

In addition to the microglial imaging angle, we have also explored the functional applications of **MITIGATE^C^** probes by synthesizing two new **MITIGATE**-series probes: **MITIGATE^PA^**(PA: photoactivatable) and **MITIGATE^PC^** (PC: photocatalyst). Using **MITIGATE^PA^**, we photoactivated single microglia in a microglia-neuron coculture system. This enabled tracking of single microglia among a collection of cells. This is important for the sparse labelling of a distinct microglial population in a region-specific manner and to follow their interactions and dynamics in the brain. The second probe, **MITIGATE^PC^**, enabled surfaceome labelling of microglia in the presence of a biotin-azide intermediate using photoproximity labelling. An unperturbed proteomic map of homeostatic microglia is difficult to achieve because the expression of genetic tools such as biotin ligase (BioID) requires viral transduction approaches in microglia, which are perturbative. **MITIGATE**^PC^ would be an ideal tool for mapping the interactome of microglia with other glial cells and neurons in the brain.

In conclusion, we have presented a novel chemical tool for imaging microglia in their homeostatic state. Unlike viral-mediated methods, this tool will be particularly useful for investigating healthy microglia and their phenotypic polarization in real-time across species. Future studies are underway to explore the possibility of the **MITIGATE** tools for investigating neuron-microglia communications in the human iPS-induced neuro-immune organoid models.

## Acknowledgements

We thank Prof. Yamuna Krishnan (University of Chicago) for her critical inputs to the manuscript. We thank Drs. Prasad Tammineni (University of Hyderabad), Deepak Nair (IISc Bangalore), Prasada Rao (NIAB, Hyderabad), Tamal Das (TIFRH), Manish Jaiswal (TIFRH) and Adish Dani (TIFRH) for their kind support. We thank Kanakam Kishore (animal facility of TIFRH) for the help with animals. We thank Akhil Mohanan from the Central Imaging Facility (TIFR Hyderabad), NIAB histology facility, and the Core Imaging Facility at the University of Hyderabad for providing access to the Imaris 3D-rendering software.

## Funding

We are thankful for the funding from the India Alliance Intermediate Fellowship of the DBT-Wellcome Trust Foundation (Award number: IA/I/21/2/505936) and for the generous support from the Department of Atomic Energy (DAE, Project Identification No. RTI 4007), India, provided to the Tata Institute of Fundamental Research (TIFR), Hyderabad.

## Author contributions

ATV and SVC conceived the project and designed all the experiments and wrote the manuscript. SVC synthesized and characterized the MITIGATE probes, and validated them in cell-culture. PC, RG and SK, established primary microglia cultures. PC cultured primary neurons and designed neuron-microglia coculture experiments. PC and SR created stable cell lines of P2RY12 constructs. SR performed probe labelling and FACS experiments in HEK29T cells. SC and RG conducted calcium imaging experiments and data analyses. PL synthesized photoproximity and photoactivation reagents. PL and PC performed photoproximity assays. PC performed single-microglia tracking experiments. SK and GR conducted probe injection experiments in the mice. SK performed cryosectioning, IHC experiments and confocal imaging of brain tissues. KJ cloned the mP2RY12-GFP construct, performed protein structure predictions and conducted the P2RY12 pull-down experiments. RM performed microinjection experiments in zebrafish and SVC imaged the zebrafish. SD provided the aryl azido biotin reagent. KM and KC contributed valuable inputs in the project.

## Declaration of interests

The authors declare no competing interest

## Data and materials availability

All the data are provided in the main text and in the supporting files.

## Notes

### Competing Interest Statement

The authors have declared no competing interest.

