## Supplemental File for "GPCR-targeted imaging and manipulation of homeostatic microglia in living systems"

INDEX

**Supplementary figure details**

Fig S1: Detailed chemical structure of different **MITIGATE**$^{C}$ Probes
Fig S2: **MITIGATE**$^{C}$ labels P2RY12 receptors at both room temperature and at 4 degrees
Fig S3: **MITIGATE**$^{C}$ and **MITIGATE**$^{NC}$ competition experiments with P2RY12 antagonist (competitor)
Fig S4: Concentration titration with **MITIGATE**$^{C}$ (0-100 nM)
Fig S5: P2RY12 receptor labelling with **MITIGATE**$^{C}$ checked with fluorescence measurement
Fig S6: Immunostaining of primary mouse microglia and neurons
Fig S7: Labelling of mP2RY12-CHO with **MITIGATE**$^{C}$ probe
Fig S8: Structural analysis of P2RY12 receptors between different species and labelling of zP2RY12-CHO with **MITIGATE**$^{C}$ probe
Fig S9: **MITIGATE**$^{C}$ targeting microglia in different regions of the adult mouse brain
Fig S10: Protein sequences for the different constructs used for the study
Fig S11: Alignment of hP2RY12, mP2RY12 and zP2RY12 protein sequences

**Methods**

Animal model subject details
Antibodies and biological reagents used
Plasmid construction and cell expression
Cell culture details: cell lines and primary cell culture details
Chemical labelling of cells
Competition assay to validate covalent labelling by fluorescence imaging
Calcium sensing analysis of CHO cells expressing hP2RY12 receptors
Flow cytometry experiments
Immunoprecipitation
Structural alignment of P2RY12 receptors from different species
Mouse brain injection protocol and experimental details: injection and brain slice preparation
Zebrafish injection protocol and experimental details
Immunostaining of cells and brain slices
Image acquisition and analysis
Software

Statistical analysis
Sholl analysis
Extended video legends
Synthetic scheme, procedure and characterization
References

**Extended videos**

SI Video 1: Calcium spikes in hP2RY12-GFP CHO cells labelled with **MITIGATE**$^{C}$
SI Video 2: Calcium spikes in hP2RY12-GFP CHO cells
SI Video 3: 3D surface rendering of **MITIGATE**$^{C}$ labelled primary microglia
SI Video 4: Whole brain microglia labelling in zebrafish with **MITIGATE**$^{C(Cy5)}$
SI Video 5: E-coli uptake by microglia in larval zebrafish brain
SI Video 6: IBA-1 colocalization with **MITIGATE**$^{C(A647)}$ labelled cell
SI Video 7: GFAP anti-colocalization with **MITIGATE**$^{C(A647)}$ labelled cell
SI Video 8: **MITIGATE**$^{C(A647)}$ labelled microglia 3D rendering (cerebellum)
SI Video 9: Photoactivation of single microglia using **MITIGATE**$^{\mathbf{PA}}$ in BV2 cells
SI Video 10: Neuron-BV2 microglia co-culture
SI Video 11: Photoactivation and single-microglia tracking in coculture

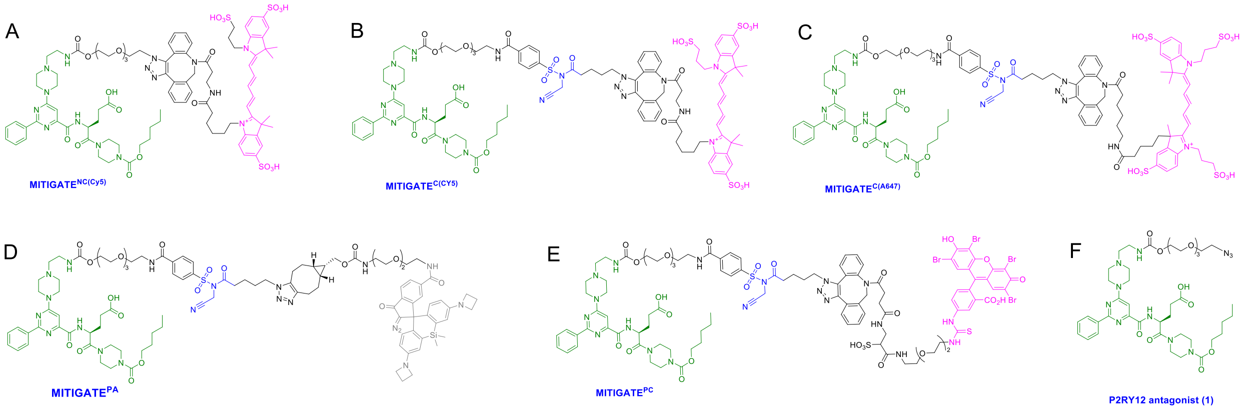


**Fig S1: Detailed chemical structure of different MITIGATE Probes:** Chemical structures of A) **MITIGATE**$^{NC(Cy5)}$, B) **MITIGATE**$^{C(Cy5)}$, C) **MITIGATE**$^{C(A647)}$, D) **MITIGATE**$^{\mathbf{PA}}$, E) **MITIGATE**$^{\mathbf{PC}}$ and F) P2RY12 antagonist. Green module: P2RY12 ligand, blue module: reactive centre and magenta module: fluorophores/photoactivatable chemical groups.

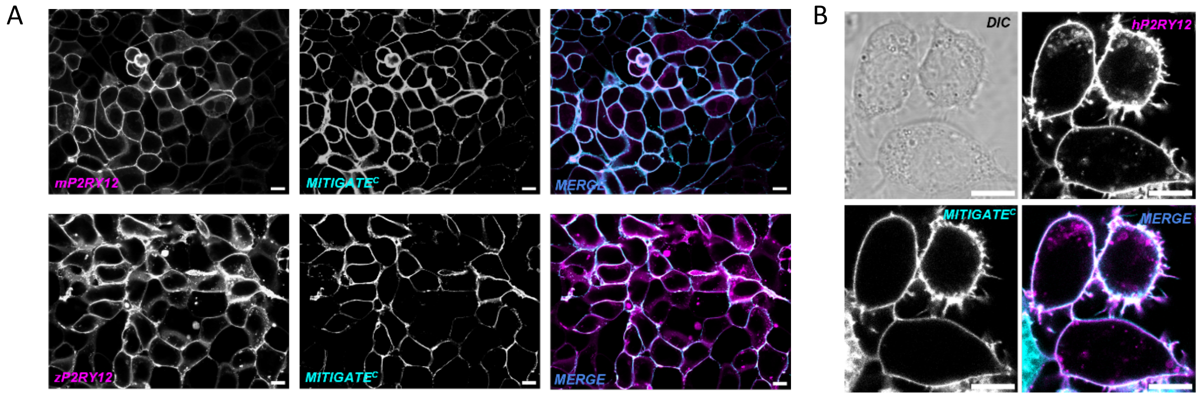


**Fig S2: MITIGATE**$^{\mathbf{C}}$ **labels P2RY12 receptors at both room temperature and at 4 degrees Celsius:** CLSM images showing colocalization of A) mP2RY12-GFP and B) zP2RY12-Staygold (magenta, $\lambda_{ex}$ = 488 nm) with 250 nM **MITIGATE**$^{\mathbf{C}}$ (cyan, $\lambda_{ex}$ = 647 nm) in HEK293T cells at 37 °C B) colocalization of hP2RY12-GFP (magenta, $\lambda_{ex}$ = 488 nm) with **MITIGATE**$^{\mathbf{C}}$ (cyan, $\lambda_{ex}$ = 647 nm) in CHO cells at 4 °C. Scale bar: 10 µm.

**Supplementary note for the Fig. S2: MITIGATE**$^{\mathbf{C}}$ labels P2RY12 receptors at both room temperature and at 4 degrees Celsius. Labelling of cells at 4 degree Celsius requires 2 hours of incubation to achieve a good signal and it reduces the receptor endocytosis.

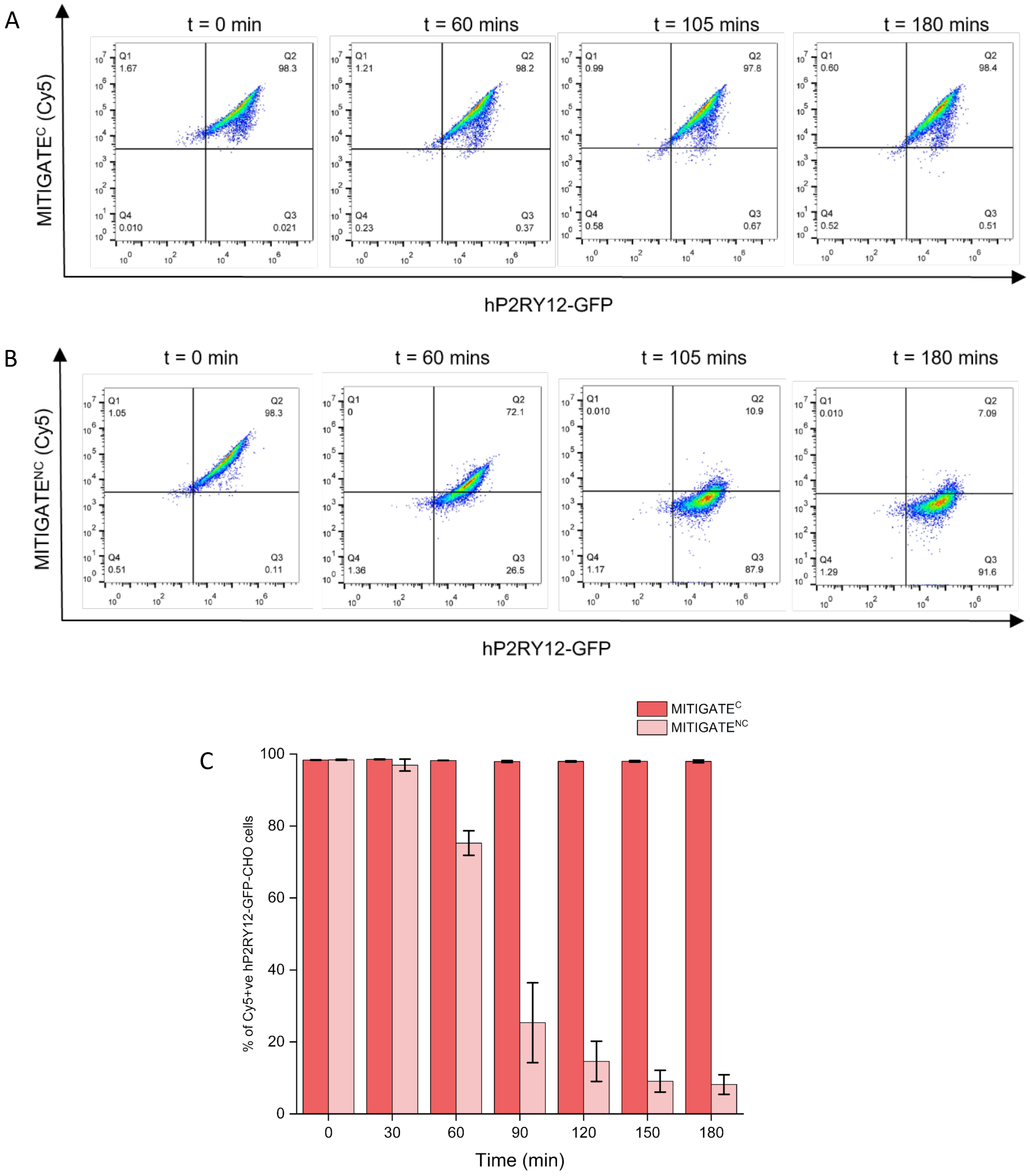


**Fig S3: MITIGATE$^{C}$ and MITIGATE$^{NC}$ competition experiments with P2RY12 antagonist (competitor):** Flow cytometry readouts showing A) irreversible binding of **MITIGATE$^{C}$** and B) reversible binding of **MITIGATE$^{NC}$** to hP2RY12-GFP expressed in CHO cells over a duration of 3 hours C) histogram plot showing relative binding of **MITIGATE$^{C}$** vs **MITIGATE$^{NC}$** to hP2RY12-GFP recorded every 30 min for 3 hours.

**Supplementary note for Fig. S3: MITIGATE$^{C}$ and MITIGATE$^{NC}$** competition experiments with P2RY12 antagonist were carried out at 4-degrees, to reduce receptor endocytosis. Non-covalent P2RY12-bound, **MITIGATE$^{NC}$** probes that are endocytosed are difficult to get competed out. The 5% residual signal observed even after 180 minutes of competitor addition is due to endocytosis. This is also verified in the confocal imaging experiments (see Fig. 1).

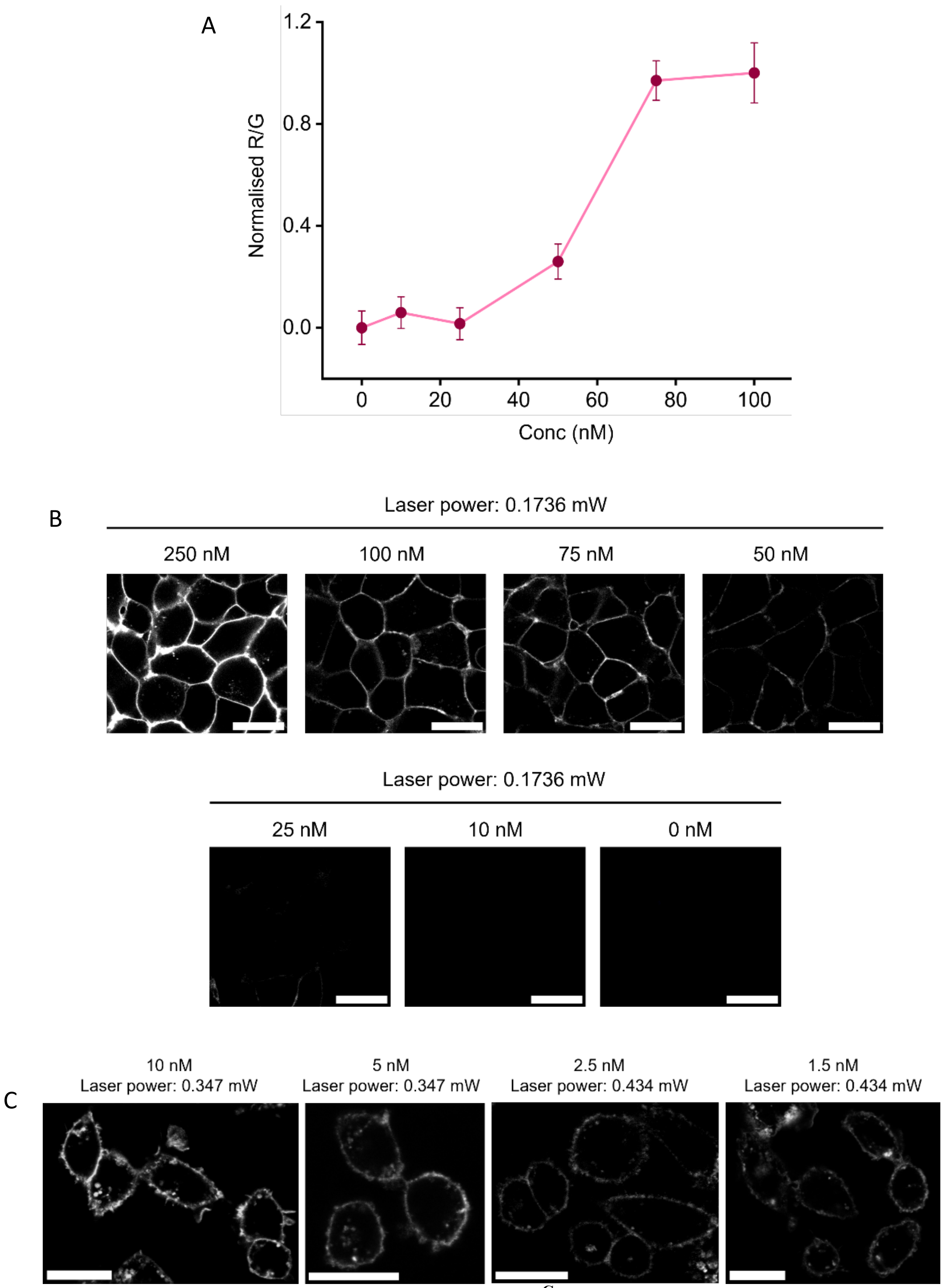


**Fig S4: Concentration titration with MITIGATE$^C$:** A) Normalized fluorescence ratio R/G (**MITIGATE$^C$** (R, red, $\lambda_{ex}$ = 647 nm, laser power = 0.1736 mW) to mP2RY12-GFP (G, green, $\lambda_{ex}$ = 647 nm, laser power = 0.02264 mW) over a concentration range of 0-100 nM of **MITIGATE$^{C(CY5)}$** in DMEM media B) CLSM images showing membrane labelling of HEK293T cells expressing mP2RY12-GFP with **MITIGATE$^C$** (grey, $\lambda_{ex}$ = 647 nm, laser power = 0.1736 mW) over a range of concentrations (0-100 nM in DMEM media) C) CLSM images showing membrane labelling of CHO cells expressing hP2RY12-GFP with **MITIGATE$^C$** (red, $\lambda_{ex}$ = 647 nm) at sub-nanomolar concentrations (10 nM, 5 nM, 2.5 nM and 1.5 nM in HBSS with $Ca^{2+}$ and $Mg^{2+}$) as a function of laser power (5-10 nM; laser power = 0.347 mW, 1.5-2.5 nM; laser power = 0.434 mW). Scale bar: 20 µm.

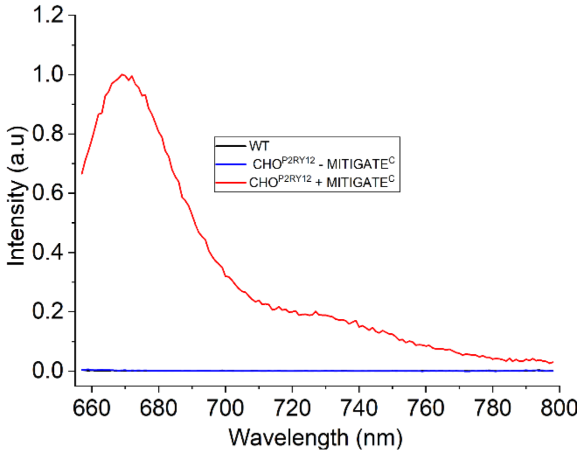


**Fig S5: Spectrofluorimetric measurement of P2RY12 receptor labelled with MITIGATE$^{C(A647)}$:** Fluorescent spectra of solubilized receptors after treatment with **MITIGATE$^{C}$** (AF647). Emission spectrum corresponding to the A647 fluorophore-P2RY12 receptor conjugate is observed at $\lambda_{ex}$ = 647 nm.

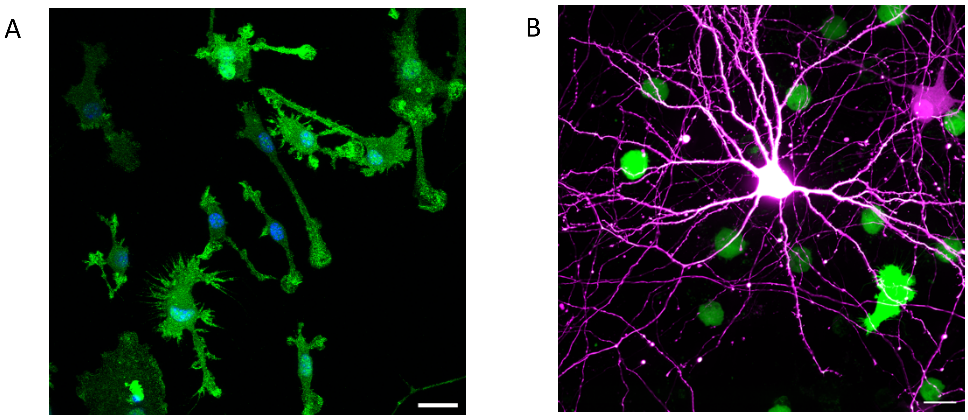


**Fig S6: Immunostaining of primary mouse microglia and neurons:** CLSM images of A) High purity of the primary microglial culture demonstrated by immunostaining with the microglia marker IBA-1 antibody; IBA-1 (green, $\lambda_{ex}$ = 488 nm) and DAPI (blue, $\lambda_{ex}$ = 405 nm) Scale bar: 20 µm. B) Representative images of neuron-microglia co-culture. tdTomoato transfected DIV10 neurons (magenta, $\lambda_{ex}$ = 554 nm) cocultured with BV2 cells expressing mP2RY12-GFP (green, $\lambda_{ex}$ = 488 nm) at a ratio 1:10 (microglia:neuron) Scale bar: 20 µm.

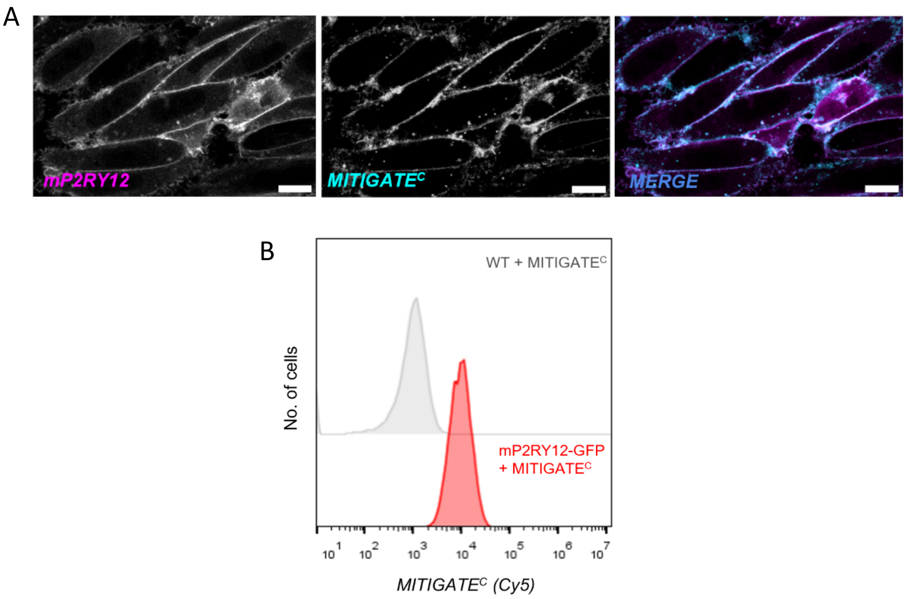


**Fig S7: Labelling of mP2RY12-CHO with MITIGATE$^{C}$ probe:** A) CLSM images showing colocalization of mP2RY12-GFP (magenta, $\lambda_{ex}$ = 488 nm) with **MITIGATE$^{C}$** (cyan, $\lambda_{ex}$ = 647 nm) in CHO cells B) Flow cytometry readouts for **MITIGATE$^{C(Cy5)}$** ($\lambda_{ex}$ = 640 nm) signal from CHO cells expressing mP2RY12-GFP (bottom panel, red) and WT-CHO (top panel, grey) respectively.Scale bar: 10 µm.

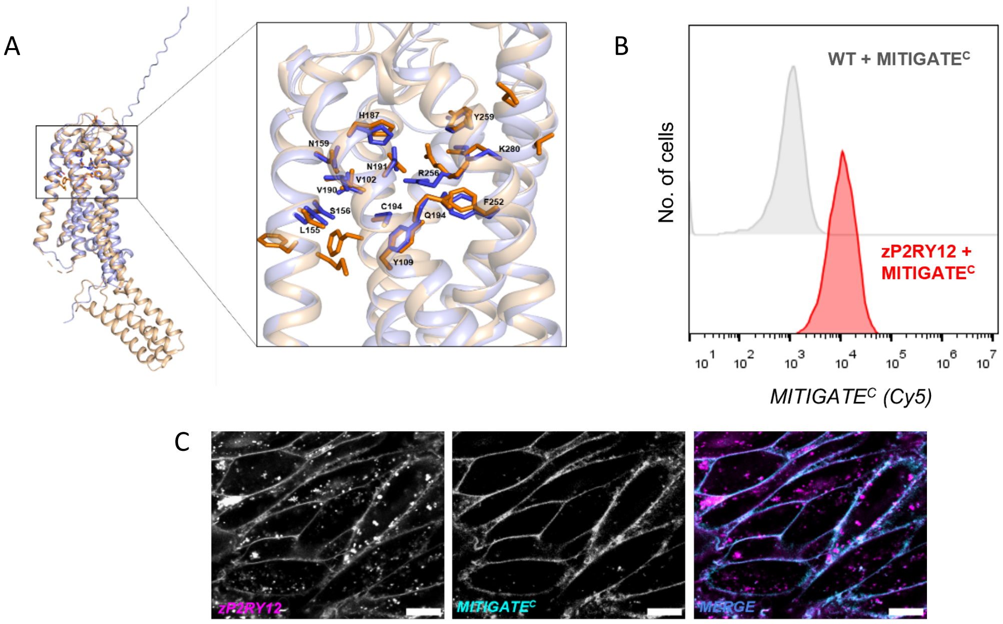


**Fig S8: Structural analysis of P2RY12 receptors between different species and labelling of zP2RY12-CHO with MITIGATE$^{C}$ probe:** A) Structural alignment of hP2RY12 receptor; PDB ID: 4PXZ (wheat, cartoon representation) with zP2RY12 receptor; UniProt ID: E7F2G6 (light blue, cartoon representation). Conserved residues interacting with antagonist are highlighted in stick conformation B) CLSM images of colocalization of zP2RY12-Staygold (magenta, $\lambda_{ex}$ = 488 nm) with **MITIGATE$^{C}$** (cyan, $\lambda_{ex}$ = 647 nm) in CHO cells. C) Flow cytometry readouts for **MITIGATE$^{C(Cy5)}$** signal from CHO cells expressing zP2RY12-Staygold (bottom panel, red) and WT-CHO (top panel, grey) respectively. Scale bar: 10 µm.

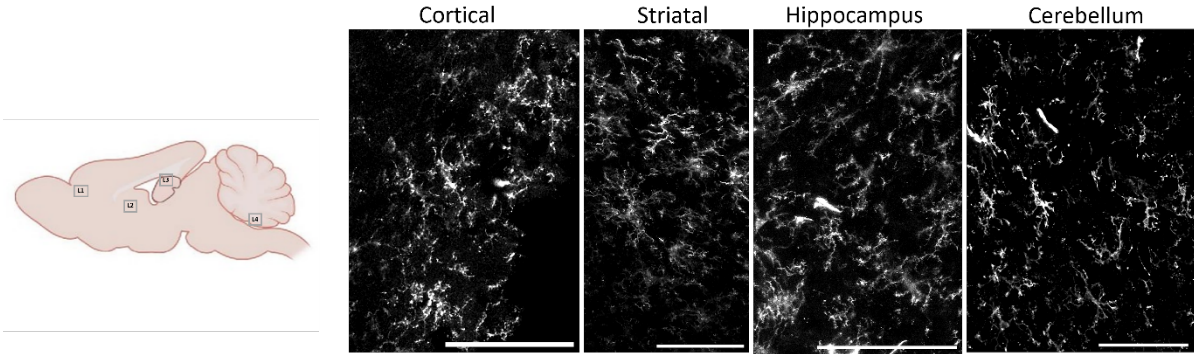


**Fig S9: MITIGATE$^{C}$ targeting microglia in different regions of the adult mouse brain:** CLSM images showing **MITIGATE$^{C}$** (grey, $\boldsymbol{\lambda}_{ex}$ = 640 nm) targeting microglia in different regions of brain in the sagittal sections. L1: Cortical region, L2: Striatal region, L3: Hippocampus regions and L4: cerebellum region. Scale bar: 100 µm.

**Supplementary note:** 5 µL of 50 µM **MITIGATE$^{C(A647)}$** was injected in each side of the brain in the intracerebroventricular space of the adult mouse.

### Methods

#### Animal model subject details

*Animal ethics procedures for mice*

All mouse experiments were conducted in accordance with the Committee for the Purpose of Control and Supervision of Experiments on Animals (CPCSEA) and were approved by the Institutional Animal Ethics Committee [IAEC]. All mice were kept *ad libitum* with food and water, under a 12 hours light-dark cycle. Healthy C57BL/6J mice with no abnormal phenotype were used for experiments and sacrificed during the light cycle. All experiments with zebrafish were done in a CPCSEA-approved zebrafish facility at Dr. Reddy’s Institute of Life Sciences in Hyderabad, India. All procedures and protocols were reviewed and approved by the Institutional Animal Ethics Committee. The “Guidelines for Experimentation on Fishes, 2021” published by CPCSEA was used as a reference.

*Zebrafish husbandry and maintenance*

The zebrafish used for the experiments were of Indian wild-caught fish origin, obtained from a commercial breeder. The fish were maintained in a semi-automated housing system (ZebTEC rack with active blue technology, Tecniplast, Italy). This system circulates filtered and aerated water to the breeding tanks and maintains stable pH of 7.0, temperature of 28°C, and a conductivity of 500 µS. The fish were reared at 14:10 light-dark cycles and were fed three times every day with either dry or live feed (pellets or live brine shrimp hatched in-house) with live feed provided at least once daily. Healthy fishes with normal behavioural and physiological parameters were used for conducting experiments. All injections were done in the light cycle.

*Zebrafish embryo collection*

For experiments, 3-6 months old male and female adult fish were set up for breeding and embryos were collected by natural spawning and raised at 28°C. Experiments were planned in the desired larval stage embryos (4 dpf).

**Antibodies and biological reagents**

The following antibodies were used in the study:
Anti-IBA-1 Rabbit mAb (#17198 Cell Signalling Technologies, IHC- 1:500/1:1000), Anti-β3-Tubulin Rabbit mAb (#5568 CST, IHC- 1:500), Anti-Alexa Fluor 647 Mouse mAb (#M647-65A-400 Immunology Consultants Laboratory, IP - 1:5000), Anti-P2Y12 Rabbit polyclonal (#APR-020 Alomone labs, IP- 1:2000), Anti-Biotin mouse mAb (#Z021 Invitrogen, IHC- 1:1000), chicken anti-GFAP polyclonal (# AB4674, Abcam, IHC- 1:1000)
Anti-rabbit Alexa Fluor 488 (#4412 CST, IF- 1:1000), Anti-rabbit Alexa Fluor 546 (#A-11010 Invitrogen, IHC- 1:1000), Anti-mouse Alexa Fluor 488 (#4408 CST, IHC-1:1000), Anti-rabbit HRP conjugated antibody ( #AB97051 AbCam, WB- 1:2000), Anti-mouse HRP conjugated antibody ( #97023 AbCam,WB- 1:2000), Anti-mouse Alexa Fluor 647 (#4410, CST, 1:1000), Anti-Chicken Alexa Fluor 488 (#AB150169, AbCam, IHC- 1:1000)

**Plasmid construction and cell expression**

*Expression of P2RY12 receptors in different cell lines*

Human P2RY12-GFP (HG17591-ACGLN) cDNA in a lentiviral backbone was sourced from Sino Biologicals. Mouse P2RY12-GFP ORF (MG58491-ACG) was sourced from Sino Biologicals and cloned in a lentiviral backbone (Addgene #113020) using Mlu1 (R3198-NEB) and Not1 (R3189-NEB) enzymes and sequence verified post cloning. Zebrafish P2RY12-StayGold ORF was gene synthesised through Twist Bioscience in a lentiviral backbone.

*Lentivirus production and transduction protocol*

Stable lines expressing human, mouse and zebrafish P2RY12 were generated via lentivirus transduction. The lentivirus particles were prepared by co-transfecting 2 µg of respective P2RY12 plasmids along with 2 µg psPAX2 (#12260, Addgene) and 1µg pMD2.G (#12259, Addgene) plasmid in HEK293T cells. Media supernatant was collected 72 hr post transfection and viral particles were concentrated using Lenti-X concentrator (#631232 Takara). HEK293T, CHO and BV2 cells were incubated with the lentiviral particles in presence of 8 µg/ml of polybrene (#H9268 Sigma) for 24 hours followed by media change and imaged at 72 hours post transduction. Cells were sorted using the fluorescent reporter (GFP or Staygold).

**Cell culture**

All cultures were maintained at 37 °C in a 5% CO2 incubator (Eppendorf cellXpert). CHO cells were cultured in Minimum essential medium eagle (MEM, Earle's salts, 2 mM L-Glutamine, 1 mM sodium pyruvate, NEAA and 1.5 g/L sodium bicarbonate) supplemented with 10 % fetal bovine serum (FBS) and 100U/ml Penicillin and 0.1mg/ml Streptomycin. BV2 cells were cultured in Dulbecco's modified eagle medium (DMEM, high glucose (4.5 g glucose per litre), 4mM L-Glutamine and sodium bicarbonate medium) supplemented with 5% heat-inactivated fetal bovine serum (HI-FBS), 1% N-2-hydroxyethylpiperazine-N-2-ethane sulfonic acid (HEPES) buffer and 100U/ml Penicillin and 0.1mg/mL Streptomycin. HEK293T cells were cultured in Dulbecco's modified eagle medium (DMEM, high glucose (4.5 g glucose per litre), L-Glutamine and sodium bicarbonate) supplemented with 10 % fetal bovine serum (FBS) and 100U/mL Penicillin and 0.1 mg/mL Streptomycin.

*Primary microglia isolation*

Primary mouse microglial cells were prepared as described in the reported literature[1]. Briefly, 2 C57BL/6J P2-P4 pups were euthanized by decapitation and the brain was harvested in dissection media to clean meninges. Cortex and hippocampus were collected and minced in 2 mL of mincing media (HBSS with $Ca^{2+}$

and $Mg^{2+}$ and 1% HEPES) using a scalpel. The minced tissue was transferred to a fresh falcon, followed by addition of 2 mL plating media (DMEM + 6% horse serum + 4% HI-FBS + 2 mM Glutamax + 1% Pen-Strep + 1 mM Sodium Pyruvate + 0.6% glucose). The tissue was triturated in 2 mL fresh plating media using a 1mL pipette, followed by addition of fresh 2 mL plating media and centrifugation at 210g for 5 minutes. Fresh plating media was added to the cell pellet and the cell suspension was plated in pre-equilibrated media on a T75 flask. Complete media was changed 48 hours post plating and the cells were grown for 17-21 days post plating with half media changes given every 3 days. After 17-21 Days In Vitro (DIV), circular microglia progenitors, growing on top of an astrocytic bed, were harvested by shaking the flask at 120 rpm for 20 minutes. The supernatant was collected and cells were pelleted by centrifugation at 210g for 7 minutes followed by plating the required number of cells for experiments. After 12 hours of plating, the media was changed to fresh plating media. Primary microglial cells were maintained in culture for 6 days post harvesting. All the experiments were done 3-4 days post plating/harvesting.

*Primary neuron culture*

Primary neuron culture protocol was adapted and modified from reported literature[2]. Briefly, E18.5 aged pups were obtained from pregnant C57BL/6J mice. Following euthanization of the pups by decapitation, the brain tissue was harvested and placed in dissection media (HBSS + 2 mM glucose). After removal of meninges, clean cortex and hippocampus were obtained and washed three times in 2 mL of dissection media each. A fresh digestion cocktail was made with prewarmed 50 mg/ml papain (LS003119, Worthington) solution in dissection media. In 4 mL of this solution, 200 µL of 10 mg/mL DNase (DN25, Sigma) was dissolved, followed by addition of this solution to the tissue. The solution was incubated with the tissue for 20 minutes at 37°C with shaking every 4 minutes. After 20 minutes, the solution was removed and the settled tissue was washed once with the dissection buffer followed by two washes with plating media. The tissue was then triturated using 1 mL tips by gently pipetting up and down till no clumps were left. Sequential trituration with 200 µL and 10 µL tips was performed, followed by pelleting the cells at 150g for 4 minutes. The pellet was resuspended in 1 mL fresh plating media (Neurobasal™ Plus Medium, 10% heat inactivated FBS, 1% Glutamax) and 50,000 cells were plated on imaging dishes. Plating media was replaced by maintenance media (Neurobasal, 1x Pen-Strep, 1% Glutamax, 2x B-27) after 4 hours of plating and the cells were kept in culture until DIV21. Primary neurons were transfected with 1.8 $\mu$g of pAAV-SFFV-tdTomato plasmid at DIV 4. The transfection mixture was made by mixing 1.8 $\mu$g of DNA with 3.3 $\mu$L of Lipofectamine™ 2000 (Cat. no 11668019) in 150 $\mu$L of Neurobasal media and incubated for 30 mins. The transfection cocktail was added to the neurons containing 150 $\mu$L of conditioned media and incubated for 1.5 hours. After incubation, full media was replaced with half conditioned media and half fresh media.

*Primary neuron-BV2 co-culture*

Primary hippocampal neurons were isolated from E18.5 pups as described above. BV2 microglia expressing mP2RY12-GFP were layered on DIV7 neurons at a ratio of 1:10 (microglia:neuron) and co-cultured for 6 hours before probe labelling.

**Sample preparation and storage**

**1** or **2** was clicked with desired fluorophores and functional chemical handles using copper-free click chemistry to obtain **MITIGATE$^{NC}$** or **MITIGATE$^{C}$** respectively. Main stocks of samples prepared in dry DMSO (mol-bio grade) were stored at -80 °C refrigerator. Working aliquots were stored in a -20 °C.

**Chemical labelling of the cells**

Cells were treated with 250 nM of **MITIGATE$^{C}$** or **MITIGATE$^{NC}$** in HBSS (with $Ca^{2+}$ and $Mg^{2+}$) for 30 min-2 hours at 37 °C in a 5 % $CO_2$ incubator. In case of CLSM assays to reduce receptor endocytosis and FACS analysis for competition, incubation was done at 4 °C for 2 hours. Cells were then washed three times with HBSS to remove excess reagent before experiments.

**Chemical labelling of endogenous P2RY12 in primary mice microglia culture**

Cultured primary microglia was treated with 250 nM **MITIGATE$^{C}$** in HBSS (with $Ca^{2+}$ and $Mg^{2+}$) for 2h at 37 °C in a 5% $CO_2$ incubator. Probe labelled microglial cells were then washed three times with HBSS buffer and imaged using a confocal microscope.

**Co-culture experiments**

For labelling BV2 microglia in neuron-microglia co-cultures, 250 nM of **MITIGATE$^{C}$** in HBSS (with $Ca^{2+}$ and $Mg^{2+}$) was added to the co-cultured cells and incubated at 37 °C in a 5 % $CO_2$ incubator for 30 minutes. Cells were then washed three times with HBSS to remove excess reagent before experiments. Cells were imaged on the Nikon spinning disk confocal system.

For photo-activating single BV2 microglia in neuron-microglia co-cultures, DIV8 neurons were co-cultured with mP2RY12-GFP expressing BV2 microglia at 1:10 (microglia:neuron) ratio for 10 hours. 250 nM of **MITIGATE$^{PA}$** in HBSS (with $Ca^{2+}$ and $Mg^{2+}$) was added to the co-culture and incubated for 30 minutes. Probe labelled cells were washed three times with HBSS and imaged using Nikon spinning disk confocal system. Photoactivation was done with the 405 nm laser using the FRAP module.

For proximity labelling studies in microglia-neuron co-cultures, DIV6 neurons were co-cultured with mP2RY12-GFP expressing BV2 microglia at 1:10 (microglia:neuron) ratio for 10 hours. Cells were washed four times with HBSS (with $Ca^{2+}$ and $Mg^{2+}$). 2 µM of **MITIGATE$^{PC}$** in HBSS (with $Ca^{2+}$ and $Mg^{2+}$) was added to the co-culture and incubated for 1 hour. Probe labelled cells were washed three times with HBSS. 100 µM of azido-biotin intermediate was added to the cells and the dish was irradiated with a custom-built green LED light (532 nm, 3 mW) for 1.5 minutes. Cells were washed three times with HBSS and fixed using 4% PFA. Cells were then immunostained with anti-biotin antibody and imaged using an Olympus FV3000 confocal system.

**Competition assay to validate covalent binding**

Cells chemically labelled with **MITIGATE$^{C}$** or **MITIGATE$^{NC}$** were imaged using CLSM (FV3000, Olympus) followed by which 1 µM competitor (P2RY12 antagonist) was added to the respective wells. After incubating for 30 minutes, imaging was done without washing.

**$Ca^{2+}$ sensing analysis of CHO cells expressing hP2RY12 receptor**

CHO cells expressing hP2RY12-GFP were treated with 100 nM **MITIGATE$^{C}$** in MEM for 1 hour at 37 °C in a 5% $CO_2$ incubator. Probe labelled cells were washed three times with MEM and then incubated for 1 hour in MEM at 37 °C. As a control, untreated cells were incubated in MEM for 2 hrs at 37ºC in 5% $CO_2$. 7 µM Rhod-4™AM (#21121 AAT Bioquest) containing 0.02% Pluronic F-127 in MEM was then incubated with the cells for 30 minutes at 37 °C in a 5% $CO_2$ incubator. Cells were washed three times with HBSS (with $Ca^{2+}$ and $Mg^{2+}$) and imaged in a widefield microscope (IX83, Olympus) equipped with a Tokai chamber. 300 µM ATP was then added to the dish and the Rhod-4™AM ($\lambda_{em}$ = 590 nm) signal was acquired at an interval of 2 s for a duration of 3 minutes.

**Flow cytometry analysis**

Chemically labelled cells were washed three times with HBSS. Cells were then incubated with 0.25% Trypsin-EDTA for 1 minute and neutralised with media. The cell suspension was centrifuged at 300 rcf for 5 minutes at 4 ºC. Supernatant was discarded and the cells were resuspended in HBSS buffer with $Ca^{2+}$ and taken for flow cytometry in ice-bath. Flow analysis was done using the flow cytometer (Beckman Coulter CytoFLEX S). A total of 10,000 events were recorded per treatment in each experiment.

For the competition assay to validate the covalent binding of the probe, P2RY12 antagonist in excess concentration (30 µM) was added to the respective MCTs and the tubes were gently vortexed at room temperature for 180 minutes. Data was recorded every 30 minutes using the flow cytometer.

**Immunoprecipitation**

WT CHO and hP2RY12-CHO cells seeded in 60 mm dishes were treated with 10 µM **MITIGATE**$^{C(A647)}$ for 1 hour at 37$^{o}$C. Untreated hP2RY12-CHO cells served as negative control. Cells were washed thrice with 1X PBS after 1 hour and then incubated with 500 µL of lysis buffer (150 mM NaCl, 10 mM Tris-HCl pH 7.4, 1 mM EDTA, 0.2 mM PMSF, 1% Triton-X-100, 0.5% NP-40, 1X protease inhibitor cocktail) with continuous shaking at 4 $^{o}$C for 30 mins followed by scraping the cells using a cell scraper. The cells were sonicated 4 times for 5 seconds each. The cell- lysate was centrifuged at maximum speed for 5 mins at 4 $^{o}$C. 200 µL of the supernatant was incubated with 25 µL of Protein-A magnetic beads for 1 hour at 4 $^{o}$C. The beads were then removed using a magnetic stand. The supernatant was incubated with 3 µg of hP2Y12 receptor antibody (APR-020, Alomone labs) overnight at 4 $^{o}$C. 25 µL of Protein-A magnetic beads was added and incubated at 4 $^{o}$C for 6 hours. The beads were pelleted using a magnetic stand, washed three times with 1X PBS. 30 µL of Laemmli sample buffer was added to the beads and heated at 70 $^{o}$C for 5 minutes followed by pelleting the beads. 10 µL of the sample was then run in 10% SDS-PAGE, transferred to PVDF membrane, blocked using 5% BSA for 1 hour at room temperature and incubated overnight with primary anti- hP2Y12 antibody (1:2000) and primary anti-AF647 antibody (1:1000). Following overnight incubation with primary antibodies, the blot was washed thrice with 1X TBST and HRP conjugated secondary antibodies were added and incubated for 1 hour at room temperature. The blots were washed thrice with 1X TBST and imaged using a Cytiva Amersham ImageQuant 800 gel doc with CCD imager.

**Structural alignment of P2RY12 receptors from different species**

The crystal structure of human P2Y12 receptor[3], PDB ID: 4PXZ (wheat, cartoon representation) was aligned with mouse P2Y12 receptor (UniProt ID: Q9CPV9) structure predicted using the ColabFold[4] v1.5.5 web server (wheat, cartoon representation) on PyMol. The receptor residues interacting with the antagonist[5] that are conserved in human and mouse are shown in stick conformation (human: orange; mouse: blue) and labelled.

The crystal structure of human P2Y12 receptor, PDB ID: 4PXZ (wheat, cartoon representation) was aligned with zebrafish P2Y12 receptor (UniProt ID: E7F2G6) structure predicted using the ColabFold v1.5.5 web server (lightblue, cartoon representation) on PyMol. The receptor residues interacting with the antagonist that are conserved in human and zebrafish are shown in stick conformation (human: orange; zebrafish: blue) and labelled.

**Injection of probe into intracerebroventricular space of adult mice**

All the experiments were conducted on 14–16 weeks old C57BL/6J male mice (weight around 26-32 g) as per the protocol adapted and modified from literature[6]. The mice were anesthetized in an induction chamber using 5% isoflurane at an air flow rate of 75 mL/minute for about 1-2 minutes. After inspecting for toe reflexes, the mice were placed on a surgical table with a nose cone anaesthesia with 3% isoflurane at flow rate of 30 mL/minute. Post disinfection using povidone iodine on the site of injection, 3-5 µL of 50 µM of **MITIGATE**$^{C}$ or DMSO control was directly injected into the intracerebroventricular space (either left side or both sides of the brain) using 31 G Hamilton syringes and the mice were returned back to their cages.

**Brain slice preparation and immunohistochemistry of brain slices**

Post 18 hours of injections, mice were transcardially perfused with 0.1 M PBS (pH 7.4), followed by 4% paraformaldehyde perfusion. The brain was harvested and placed directly in 4% paraformaldehyde overnight. After three washes with 0.1 M PBS, the brain was cryoprotected in 30% sucrose solution for 2 days at 4 $^{o}$C. The brain was then embedded in Optimum Cutting Temperature (OCT) media and flash frozen using dry ice/ethanol mixture. The brain was further sliced into 50-100 µm sections using Cryostar NX50 Cryostat. The sections were air dried at room temperature and fixed using 4% PFA at room temperature for 10 minutes and washed three times with 0.1M PBS. The slices were then incubated in a blocking solution (5% NGS in 0.1M PBS with 0.3% Triton X-100) for 1 hour at room temperature. Sections were further incubated with primary antibodies diluted in blocking solution and incubated at 4 $^{o}$C overnight. The slices were then washed three times with 0.1 M PBS containing 0.3% Triton X-100 for 5 minutes each and then incubated with respective secondary antibodies solution containing DAPI (1 µg/mL in 0.1M PBS), prepared in the blocking solution, for 2 hours. The slices were washed three times with 0.1 M PBS containing 0.3%

Triton-X 100 before being mounted on glass slides. The sections were imaged using Leica stellaris 5 upright microscope.

#### Brain microinjections in zebrafish larvae

Microinjections were conducted under a stereomicroscope in order to monitor and validate each successful injection. The larvae were anaesthetised and mounted in a low-melting agarose (1.5 %) bed with capillary well and dorsally oriented for accessing optic tectum. Needle was placed directly above the brain optic tectum. The skin was gently punctured with the needle followed by pressing the PicoPump foot pedal once to eject 3 nL (injection pressure = 297 hPa, injection time = 0.2 s, compensation pressure = 7 hPa) of the desired reagent. Injections were done only on one side.
For the chemical labelling experiments, 3 nL of **MITIGATE**$^{C(A647)}$ (10 µM) in HBSS (with $Ca^{2+}$ and $Mg^{2+}$) was injected into the optic tectum and the larvae were incubated in E3 buffer (5 mM NaCl, 0.17 mM KCl, 0.33 mM $CaCl_2$, 0.33 mM $MgSO_4$ in water) for 3 hours.
For the phagocytosis experiments, post injection of **MITIGATE**$^{C(A647)}$, larvae were incubated in the E3 buffer for 3 hours. The larvae were then anaesthetised using tricaine (0.003% in E3 buffer) and 3 nL pHrodo-*E. Coli* (100 particles per nL, P35361, Invitrogen) was injected into the optic tectum. Larvae were incubated in the E3 buffer for 30 min.
For imaging, larvae were anaesthetised in tricaine (0.003% in E3 buffer) and embedded in a 1.3 % agarose bed (prepared in 0.003% tricaine in E3 buffer) and were imaged using Leica stellaris 5 upright microscope.

#### Immunostaining

Cells were treated with 4 % paraformaldehyde (PFA) for 10 minutes. PFA was aspirated and the fixed cells were washed three times with 0.1 M PBS for 5 minutes each. Cells were then incubated in a blocking buffer (5 % BSA in 0.1 M PBS) in 1X PBS for 40 minutes. Primary antibody solution, prepared in PBS, was then added to the cells and incubated overnight at 4 $^{o}$C. Primary antibody solution was aspirated and the cells were washed three times with 1X PBS for 5 minutes each. Secondary antibody was then incubated with the cells in dark for 2 hours at 37 $^{o}$C. The cells were then washed thrice with 1X PBS for 5 minutes each followed by incubation with DAPI (1 µg/mL in 0.1M PBS) in the dark for 10 minutes at 37 $^{o}$C. DAPI staining solution was aspirated and the cells were washed three times with 1X PBS for 5 minutes. Imaging was done in 1X PBS using a confocal laser scanning microscope (FV3000, Olympus) equipped with a 60x, 1.42 NA, oil objective.

#### Image acquisition and analysis

Widefield images for calcium sensing studies were captured using Olympus IX83 inverted microscope equipped with 60x, 1.42 NA, oil immersion objective. Cy5, Rhod-4-AM and GFP were excited using Cy5 channel (excitation filter: 595-645nm, emission filter: 665-715 nm, dichroic mirror: 655 nm), TRITC channel (excitation filter: 540- 550 nm, emission filter: 575- 625 nm, dichroic mirror: 570 nm) and FITC channel (excitation filter: 470-495 nm, emission filter: 510-550 nm, dichroic mirror: 505 nm) respectively. Fluorescence emission was recorded using digital CMOS camera C11440-42U40. Live cell imaging was performed in the Tokai incubation chamber with regulated humidity and temperature. Time of calcium spikes was compared by normalization of fluorescence intensity of Rhod-4$^{TM}$AM in a cell at each time point to the maximum intensity. Amplitude of calcium spikes was analysed by the ratio of change of fluorescence intensity of Rhod-4-AM at a time to the initial fluorescence intensity in each cell.

Confocal images from CHO, HEK293T, BV2 cells and primary microglial cultures were captured using Olympus confocal laser scanning microscope Fluoview FV3000 equipped with 60x, 1.42 NA, oil immersion objective. DAPI, GFP and Cy5 were excited sequentially using 405 nm, 488 nm and 640 nm wavelengths respectively using optically pumped semiconductor lasers. Fluorescence emission was recorded using a hybrid photomultiplier tube detector (HyD). Live cell imaging was performed in the Tokai incubation chamber with regulated humidity and temperature.

Zebrafish brain and mice brain slices were imaged using Leica stellaris 5 upright microscope. Zebrafish brain images were captured with 40x, 0.8 NA, water immersion objective. Brain slice images were captured with 63x, 1.4 NA, oil immersion objective. DAPI, AF488, pHrodo, TRITC and Cy5/A647 were excited sequentially using 405 nm, 488 nm, 514 nm, 561 nm and 638 nm wavelengths respectively using optically pumped semiconductor lasers. Fluorescence emission was recorded using a hybrid photomultiplier tube detector (HyD).

Confocal images for co-cultures of primary neurons and BV2 cells were captured using Nikon eclipse Ti2-E inverted microscope with Yokogawa CSU-W1 SoRA super resolution spinning disk confocal microscope with 60x, 1.42 NA, oil immersion objective. DAPI, GFP and Cy5 were excited using 405 nm 488 nm laser, 640 nm wavelengths respectively using diode pumped semiconductor lasers. Photoactivation studies were carried out with laser scanning ablation set-up. Fluorescence emission was recorded using a 16-bit CMOS image sensor. Live imaging was performed in the OKOlab live cell chamber with regulated temperature, humidity and stable $CO_2$.

**Sholl analysis**

Sholl analysis was performed on microglia using the SNT module of neuroanatomy plug-in in imageJ. Briefly, images acquired from **MITIGATE**$^{C(A647)}$ treated brain slices were processed and skeletonised using a reported protocol[7]. Skeletonised images were processed using the sholl analysis 4.2.1 option in the SNT module of neuroanatomy plug-in, keeping the start radius 2.5 µm, step size 1.5 µm and end radius 60 µm. Number of intersections derived from images of distal and proximal microglia from the site of injury were plotted as a function of distance from soma. For each group, the average number of intersections for 10 cells were calculated.

**Software**

All DNA and protein sequences were visualised and analysed on SnapGene viewer. The schematics were created using online editing software, Biorender. All quantifications and statistical tests were performed using OriginPro (version 2022). All images were visualised and analysed using ImageJ win32 software (version 1.54m). Flow cytometry data was analysed and plotted using FlowJo. All the chemical structures were created and mass calculations were done using ChemDraw (version 21.0.0.).

Imaris image processing: 3D-images of microglia were generated using Imaris 10.0 software (Bitplane). Briefly, z-stacks (step size = 1 micron, 20-30 steps) of images are imported to Imaris. Surface volume feature was used to 3D-render the images. Animation was created and recorded using the 3D-animation tool in Imaris. Single-view of the rendered images was captured using the animation snapshot tool. Absolute intensity was used for optimized visualization of green (IBA-1 or GFAP) and red (MITIGATE (CY5) or A647) channels. Threshold values were adjusted to aid the best visualization of surface features.

**Statistical analysis**

A two-sample t-test (Welch's t-test) was used to compare calcium spike amplitudes between two sets of data and to calculate p-values. For Sholl analysis, one-tailed Student's t-test was performed to calculate p values for each distance point plotted. Mean values and standard deviations are shown in all figures.

1. Human P2RY12-GFP

MQAVDNLTSAPGNTSLCTRDYKITQVLFPLLYTVLFFVGLITNGLAMRIFFQIRSKSNFIIFLKNTV
ISDLLMILTFPFKILSDAKLGTGPLRTFVCQVTSVIFYFTMYISISFLGLITIDRYQKTTRPFKTSNPK
NLLGAKILSVVIWAFMFLLSLPNMILTNRQPRDKNVKKCSFLKSEFGLVWHEIVNYICQVIFWINF
LIVIVCYTLITKELYRSYVRTRGVGKVPRKKVNVKVFIIIAVFFICFVPFHFARIPYTLSQTRDVFDC
TAENTLFYVKESTLWLTSLNACLDPFIYFFLCKSFRNSLISMLKCPNSATSLSQDNRKKEQDGGDP
NEETPMGGGGSVSKGEELFTGVVPILVELDGDVNGHKFSVSGEGEGDATYGKLTLKFICTTGKLP
VPWPTLVTTLTYGVQCFSRYPDHMKKHDFFKSAMPEGYVQERTIFFKDDGNYKTRAEVKFEGD

TLVNRIELKGIDFKEDGNILGHKLEYNYNSHNVYIMADKQKNGIKANFKVRHNIEDGSVQLADH
YQQNTPIGDGPVLLPDNHYLSTQSALSKDPNEKRDHMVLLEFVTAAGITLGMDELYK

hP2RY12; GS linker; GFP

2. Mouse P2RY12-GFP

MDVPGVNTTSANTTFSPGTSTLCVRDYKITQVLFPLLYTVLFFAGLITNSLAMRIFFQIRSKSNFIIF
LKNTVISDLLMILTFPFKILSDAKLGAGPLRTLVCQVTSVTFYFTMYISISFLGLITIDRYLKTTRPF
KTSSPSNLLGAKILSVVIWAFMFLISLPNMILTNRRPKDKDVTKCSFLKSEFGLVWHEIVNYICQVI
FWINFLIVIVCYSLITKELYRSYVRTRGSAKVPKKKVNVKVFIIIAVFFICFVPFHFARIPYTLSQTR
AVFDCSAENTLFYVKESTLWLTSLNACLDPFIYFFLCKSFRNSLTSMLRCSNSTSTSGTNKKKGQE
GGEPSEETPMGGGGSVSKGEELFTGVVPILVELDGDVNGHKFSVSGEGEGDATYGKLTLKFICTT
GKLPVPWPTLVTTLTYGVQCFSRYPDHMKKHDFFKSAMPEGYVQERTIFFKDDGNYKTRAEVK
FEGDTLVNRIELKGIDFKEDGNILGHKLEYNYNSHNVYIMADKQKNGIKANFKVRHNIEDGSVQ
LADHYQQNTPIGDGPVLLPDNHYLSTQSALSKDPNEKRDHMVLLEFVTAAGITLGMDELYK

mP2RY12; GS linker; GFP

3. Zebrafish P2RY12-GFP

MEQTTQLSFSNSSVSNSSSCSRDGALKTIVFPVLYSILLILGLSLNALAAWVFLRIPSKSHFIIYLKNI
VVADIIMTLTFPFKILSDANVASVGIRIFVCRVSSVLFYLTMYISILFFGLISIDRCRKTMWPFVGTN
PKRLLHRKLLSGVIWTSLLALSLPNVILTSRPNIGERFKCSDLKTELGLQWHEVVNHICQVIFWGN
LIIVIICYTLISRELYKSYARTRSPRGTQSKKKHIQVNVFLVLAVFFICFVPFHFARVPYTISQTRALL
FSCPQKLFFFKLKESTLWLSSLNSVLDPLIYFFLCKSFRSSLFNVMRLAPGRCRILRELGTDSAGDQ
QGNALTGGGGSMASTPFKFQLKGTINGKSFTVEGEGEGNSHEGSHKGKYVCTSGKLPMSWAAL
GTSFGYGMKYYTKYPSGLKNWFHEVMPEGFTYDRHIQYKGDGSIHAKHQHFMKNGTYHNIVEF
TGQDFKENSPVLTGDMNVSLPNEVQHIPRDDGVECPVTLLYPLLSDKSKCVEAHQNTICKPLHN
QPAPDVPYHWIRKQYTQSKDDTEERDHICQSETLEAHL

zP2RY12; GS linker; StayGold

**Fig S10: Protein sequences of different constructs:** Protein sequences for 1. Human P2RY12-GFP, 2. Mouse P2RY12-GFP and 3. Zebrafish P2RY12-Staygold

**Alignment of hP2RY12, mP2RY12 and zP2RY12 protein sequences**

```
hP2RY12         ------MQAVDNLTSAPGNTSLCTRDYKITQVLFPLLYTVLFFVGLITNGLAMRIFFQIR
mP2RY12         MDVPGVNTTSANTTFSPGTSTLCVRDYKITQVLFPLLYTVLFFAGLITNSLAMRIFFQIR
zP2RY12         ---MEQTTQLSFSNSSVSNSSSCSRDGALKTIVFPVLYSILLILGLSLNALAAWVFLRIP
                             . : ..:: * **  :. ::**:**::*:: **  *.**  :*::*

hP2RY12         SKSNFIIFLKNTVISDLLMILTFPFKILSDAKLGTGPLRTFVCQVTSVIFYFTMYISISF
mP2RY12         SKSNFIIFLKNTVISDLLMILTFPFKILSDAKLGAGPLRTLVCQVTSVTFYFTMYISISF
zP2RY12         SKSHFIIYLKNIVVADIIMTLTFPFKILSDANVASVGIRIFVCRVSSVLFYLTMYISILF
                ***:***:*** *::*::* ***********::.:  :* :**:*:** **:****** *

hP2RY12         LGLITIDRYQKTTRPFKTSNPKNLLGAKILSVVIWAFMFLLSLPNMILTNRQPRDKNVKK
mP2RY12         LGLITIDRYLKTTRPFKTSSPSNLLGAKILSVVIWAFMFLISLPNMILTNRRPKDKDVTK
zP2RY12         FGLISIDRCRKTMWPFVGTNPKRLLHRKLLSGVIWTSLLALSLPNVILTSR-PNIGERFK
                :***:***  **  **  :.*..**  *:** ***: :: :****:***.* *.  :  *

hP2RY12         CSFLKSEFGLVWHEIVNYICQVIFWINFLIVIVCYTLITKELYRSYVRTRG--VGKVPRK
mP2RY12         CSFLKSEFGLVWHEIVNYICQVIFWINFLIVIVCYSLITKELYRSYVRTRG--SAKVPKK
zP2RY12         CSDLKTELGLQWHEVVNHICQVIFWGNLIIVIICYTLISRELYKSYARTRSPRGTQSKKK
                ** **:*:** ***:**:******* *::***:**:**::***:**.***.    :  :*
```

```
hP2RY12         KVNVKVFIIIAVFFICFVPFHFARIPYTLSQTR-DVFDCTAENTLFYVKESTLWLTSLNA
mP2RY12         KVNVKVFIIIAVFFICFVPFHFARIPYTLSQTR-AVFDCSAENTLFYVKESTLWLTSLNA
zP2RY12         HIQVNVFLVLAVFFICFVPFHFARVPYTISQTRALLFSCPQKLFFFKLKESTLWLSSLNS
                :::*:**:::**************:***:****  :*.*. :  :* :*******:***:

hP2RY12         CLDPFIYFFLCKSFRNSLISMLKCPNSATSLSQDNRKKEQDGGDPNEETPM
mP2RY12         CLDPFIYFFLCKSFRNSLTSMLRCSN-STSTSGTNKKKGQEGGEPSEETPM
zP2RY12         VLDPLIYFFLCKSFRSSLFNVMRLAPGRCRILRELGTDSAGDQQGNALT--
                 ***:**********.** .::: .           ..   . : .  *
```

**Fig S11: Alignment of hP2RY12, mP2RY12 and zP2RY12 protein sequences:** Multiple sequence alignment of P2Y12 receptors from *Homo sapiens* (hP2RY12), *Mus musculus* (mP2RY12) and *Danio rerio* (zP2RY12). The CLUSTALW Multiple Sequence Alignment Program (version 1.83, Feb 2003). The antagonist binding site is highlighted in red dashed boxes.

**Supplementary video legends**

**SI Video 1. Calcium spikes in hP2RY12-GFP CHO cells labelled with MITIGATE$^{C}$:** Time lapse images showing calcium spikes in **MITIGATE**$^{C(Cy5)}$ labelled hP2RY12-CHO cells treated with 300 µM ATP. Calcium spikes were observed as an increase in fluorescence intensity of Rhod-4$^{TM}$AM dye (red, $\lambda_{ex}$= 561 nm), measured at every 2-seconds in Olympus IX83 widefield microscope.

**SI Video 2. Calcium spikes in hP2RY12-GFP CHO cells:** Time lapse images showing calcium spikes in hP2RY12-CHO cells treated with 300 µM ATP. Calcium spikes were observed as an increase in fluorescent intensity of Rhod-4$^{TM}$AM dye (red, $\lambda_{ex}$= 561 nm), measured at every 2-seconds in Olympus IX83 widefield microscope.

**SI Video 3. 3D surface rendering of MITIGATE$^{C}$ labeled primary microglia:** Primary microglia labeled with **MITIGATE**$^{C(Cy5)}$ (magenta, $\lambda_{ex}$= 647 nm) were imaged in Olympus FV3000 confocal system. The z- projected images were surface 3D rendered with the 3D viewer plugin in ImageJ. The video was exported at 20 frames per second.

**SI Video 4. Whole brain microglia labeling in zebrafish:** Leica Stellaris 3D rendered animation of microglia in zebrafish (4dpf, whole brain) highlighted with **MITIGATE**$^{C(Cy5)}$ (green, $\lambda_{ex}$= 640 nm).

**SI Video 5. E. coli uptake by microglia in larval zebrafish brain:** Leica Stellaris 3D rendered animation of pHrodo E. coli (red, $\lambda_{ex}$= 514 nm) uptake by **MITIGATE**$^{C(Cy5)}$ (green, $\lambda_{ex}$= 640 nm) labelled microglial cells in zebrafish brain (4dpf).

**SI Video 6. IBA-1 colocalization with MITIGATE$^{C(A647)}$ labelled cells in adult mice brain slices:** Imaris 3D rendering of **MITIGATE**$^{C(A647)}$ treated brain slices co-stained with IBA-1 antibody showing colocalization of IBA-1 (green, $\lambda_{ex}$= 546 nm) signals in **MITIGATE**$^{C(A647)}$ (red, $\lambda_{ex}$= 640 nm) labeled cells.

**SI Video 7. GFAP anti-colocalization with MITIGATE labelled cell in adult mice brain slices:** Imaris 3D rendering of **MITIGATE**$^{C(A647)}$ treated brain slices co-stained with GFAP antibody showing anti-colocalization of GFAP (green, $\lambda_{ex}$= 488 nm) signals in **MITIGATE**$^{C(A647)}$ (red, $\lambda_{ex}$= 640 nm) labeled cells.

**SI Video 8. 3D rendering of MITIGATE$^{C(A647)}$ labelled microglia in cerebellum of adult mouse brain:** Imaris 3D rendering of microglia labeled with **MITIGATE**$^{C(A647)}$ (red, $\lambda_{ex}$= 640 nm) in the cerebellum of adult mice brain.

**SI Video 9. Photoactivation of single microglia using MITIGATE$^{PA}$ in BV2 cells:** Time Lapse video of localized photolysis of **MITIGATE**$^{PA}$ with a 405 nm laser enabling visualization of single microglia (red, $\lambda_{em}$= 660 nm) in the mP2RY12-GFP (green, $\lambda_{ex}$= 488 nm) expressing BV2 cells.

Photolysis was performed using the FRAP module in the Nikon eclipse Ti2-E inverted microscope and images were captured at frame rate of 30 secs per frame.

**SI Video 10. Neuron-BV2 microglia co-culture:** Time Lapse imaging of mP2RY12-GFP (green, $\lambda_{ex}$= 488 nm) expressing BV2 cells co-cultured with tdTomato transfected DIV7 (red, $\lambda_{em}$= 554 nm) primary neurons in 1:10 ratio. Images were captured at frame rate of 2 mins per frame.

**SI Video 11. Photoactivation and single-microglia tracking in coculture:** Time Lapse video of localized photolysis of **MITIGATE**$^{PA}$ with a 405 nm laser enabling visualization of single microglia (red, $\lambda_{em}$= 660 nm) in the microglia-neuron co-culture system where mP2RY12-GFP (green, $\lambda_{ex}$= 488 nm) expressing BV2 cells co-cultured with tdTomato transfected DIV8 primary neurons (red, $\lambda_{em}$= 554 nm) in 1:10 ratio. Photolysis was performed using the FRAP module in the Nikon eclipse Ti2-E inverted microscope and images were captured at frame rate of 2 mins per frame.

### Materials and methods for organic synthesis

Chemicals used for the syntheses were purchased from Sigma Aldrich and TCI, and were used as received. TLC analyses were performed on aluminium support silica gel 60 matrix with binder organic polymer fluorescent indicator, and column chromatography was performed on 200–400 mesh silica gel. $^{1}H$ NMR, $^{13}C$ NMR spectra were recorded on a Bruker Avance 300 MHz DPX spectrometer using 1,1,1,1-tetramethylsilane (TMS) as the internal standard. Molecular masses were recorded by LC-MS using an Agilent 1290 infinity II/6530 Q-TOF LC/MS. Vacuubrand chemistry-hybrid RC 6 pump with 5.9 $m^3/h$ pumping speed was used for vacuum drying all the compounds. All the solvents were dried by heating under reflux with $CaCl_2$ and were stored over activated 3 Å molecular sieves. Analytical separations of sensitive compounds were performed on glass support silica gel 60 matrix preparative TLC plates of 1000 µm layer thickness, 20-40 µm particle size and 60 Å medium pore diameter. Reactive compounds with reactive acylsulfonamide groups were handled by maintaining dry conditions under nitrogen atmosphere. All reactive intermediates were vacuum dried and stored at -20 °C.

### Synthesis and Characterization

**1** and **2** were synthesized using multi-step organic synthesis and were characterized using $^{1}H$ NMR spectroscopy and mass spectrometry (**Schemes 1 & 2**). **1a, 1d** and **2a** were synthesised as per the reported procedures.[8–10]

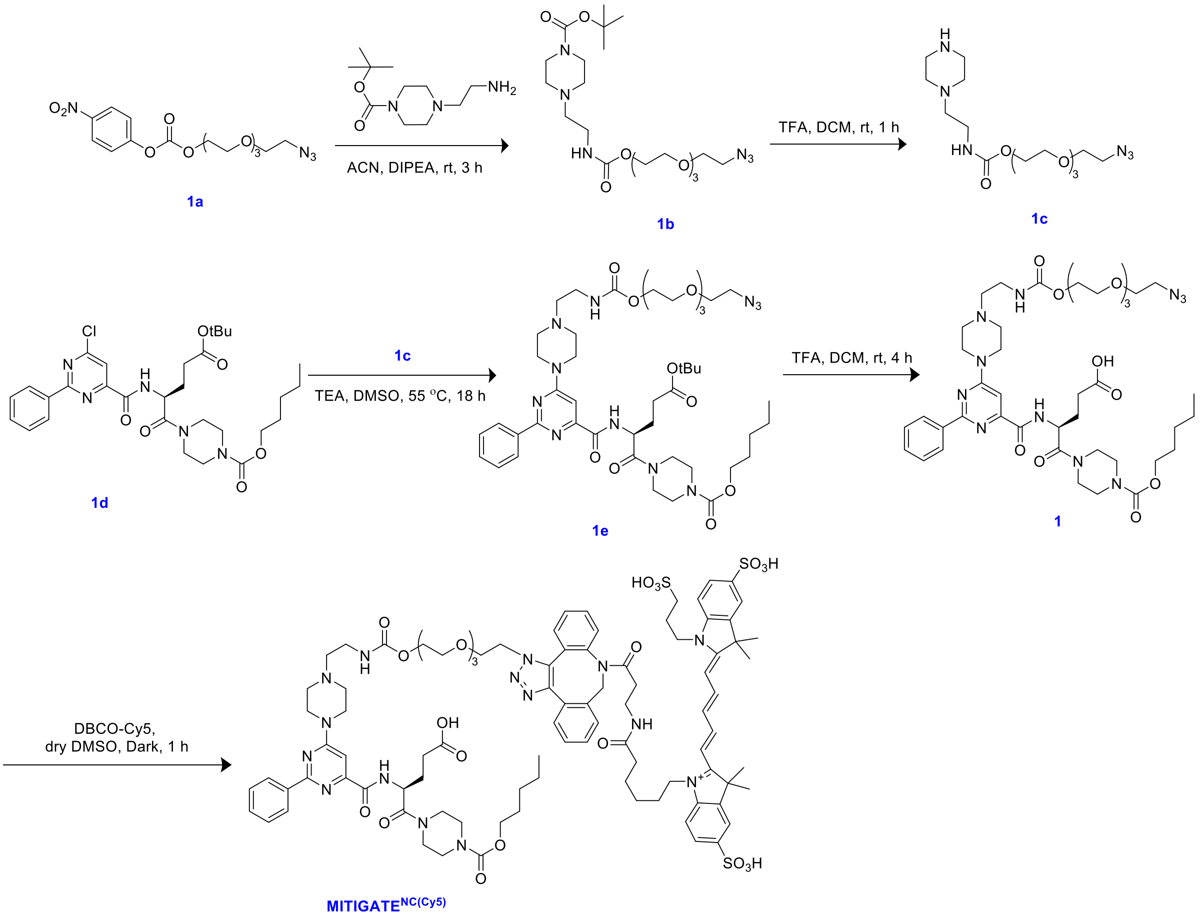


**Scheme 1:** Scheme for the synthesis of **MITIGATE**$^{NC(Cy5)}$

**1b:** A flame dried 100 mL round bottom flask equipped with a magnetic bead under nitrogen atmosphere was loaded with **1a** (0.25 g, 0.676 mmoles) and was dissolved in 2 mL dry ACN. DIPEA (0.13 mL, 1.015 mmoles) was added to the solution and stirred for 2 minutes followed by addition of tert-Butyl 4-(2-aminoethyl) piperazine-1-carboxylate (0.202 g, 0.879 mmoles). The reaction mixture was stirred at room temperature for 3 hours. The reaction progress was monitored using TLC. Upon completion of the reaction, solvent was removed under reduced pressure. DCM was added to the crude mixture and the organic phase was washed with brine and dried over $Na_2SO_4$. The crude mixture was purified using silica gel column chromatography in 10% MeOH in DCM to afford desired product as colourless oil (Yield = 95 %). $R_f$ = 0.3 (8 % MeOH in DCM)); $^1$H NMR (300 MHz, $CDCl_3$) δ (ppm): 5.22 (s, 1H), 4.16 (m, 2H), 3.6 (m, 11H), 3.33 (m, 6H), 3.21 (m, 2H), 2.4 (m, 2H), 2.31 (m, 4H), 1.59 (m, 3H), 1.38 (s, 9H), 1.18 (m, 3H).

**1c:** A 50 mL round bottom flask equipped with a magnetic bead was loaded with **1b** (0.12 g, 0.25 mmoles) and was dissolved in 5 mL DCM. 0.5 mL of TFA was added to the solution and the mixture was stirred for 1 hour. The reaction progress was monitored using TLC. Upon completion of the reaction, the reaction mixture was neutralized using 1N NaOH solution. DCM was added to the crude mixture. The organic phase was washed with brine, dried over $Na_2SO_4$ and evaporated to dryness to afford the desired product as colourless oil in quantitative yield. **1c** acquired was used for the next step without further purification.

**1e:** A 10 mL round bottom flask equipped with a magnetic bead was loaded with **1d** (200 mg, 0.152 mmoles) was dissolved in 2 mL DMSO. TEA (0.027 mL, 0.197 mmoles) was added to the solution, followed by **1e** (76 mg, 0.1580 mmoles). The reaction mixture was stirred for 18 hours. The reaction progress was monitored using TLC. Upon completion of the reaction, solvent was removed under reduced pressure. DCM was added to the crude mixture and the organic phase was washed with brine, dried over $Na_2SO_4$ and evaporated to dryness to afford the desired product as colourless oil (Yield = 98 %). $R_f$ = 0.55 (3% MeOH in DCM); Calculated mass: 939.52 g/mol, experimental mass: 940.53 g/mol [M+H$^+$]. $^1$H NMR (300 MHz, $CDCl_3$) δ (ppm): 8.90 (d, J = 8.1 Hz, 1H), 8.38 (s, 2H), 7.41 (d, J = 5.6 Hz, 3H), 5.23 (d, J = 3.5 Hz, 1H), 4.19 (d, J = 5.8 Hz, 3H), 4.12 – 3.74 (m, 6H), 3.61 (d, J = 4.2 Hz, 16H), 3.47 (s, 5H), 3.32 (d, J = 5.4 Hz, 3H), 1.45 – 1.29 (m, 15H), 0.81 (m, 12H). $^{13}$C NMR (300 MHz, $CDCl_3$) δ (ppm): 171.11, 168.85, 163.02, 162.15, 161.95, 155.38, 154.67, 154.40, 136.39, 129.65, 127.34, 127.21, 123.74, 97.67, 69.64, 69.50, 69.03, 68.66, 64.99, 62.98, 56.04, 51.51, 49.65, 47.24, 44.79, 44.35, 42.80, 42.39, 41.04, 36.35, 33.27, 30.89, 30.01, 29.73, 29.28, 28.66, 28.33, 27.63, 27.31, 27.08, 21.66, 21.33, 13.10, 12.96, 7.59

**1:** A 10 mL round bottom flask equipped with a magnetic bead was loaded with **1e** (8 mg, 8.5 mmoles) and was dissolved in 2 mL DCM. 0.4 mL of TFA was added to the solution and the mixture was stirred for 4 hours. The reaction progress was monitored using TLC. Upon completion of the reaction, the reaction mixture was neutralized using 1N NaOH solution. DCM was added to the crude mixture. The organic phase was washed with brine, dried over $Na_2SO_4$ and evaporated to dryness to afford the desired product as colourless waxy solid (Yield = 90%). **1** was used without further purification. Calculated mass: 883.46 g/mol, experimental mass: 884.46 g/mol [M+H]$^+$.

**MITIGATE**$^{NC(Cy5)}$: Sulfo-DBCO-Cy5 (2.3 mg, 2.26 µmoles) in 1 µL dry DMSO (molecular biology grade) was mixed with **1** (2 mg, 2.26 µmoles) in 1 µL dry DMSO in the dark for 1 hour. Product formation was verified using mass spectrometry. Calculated mass: 1892.77 g/mol, experimental mass: 1892.77 g/mol (deconvoluted mass) [M]$^+$

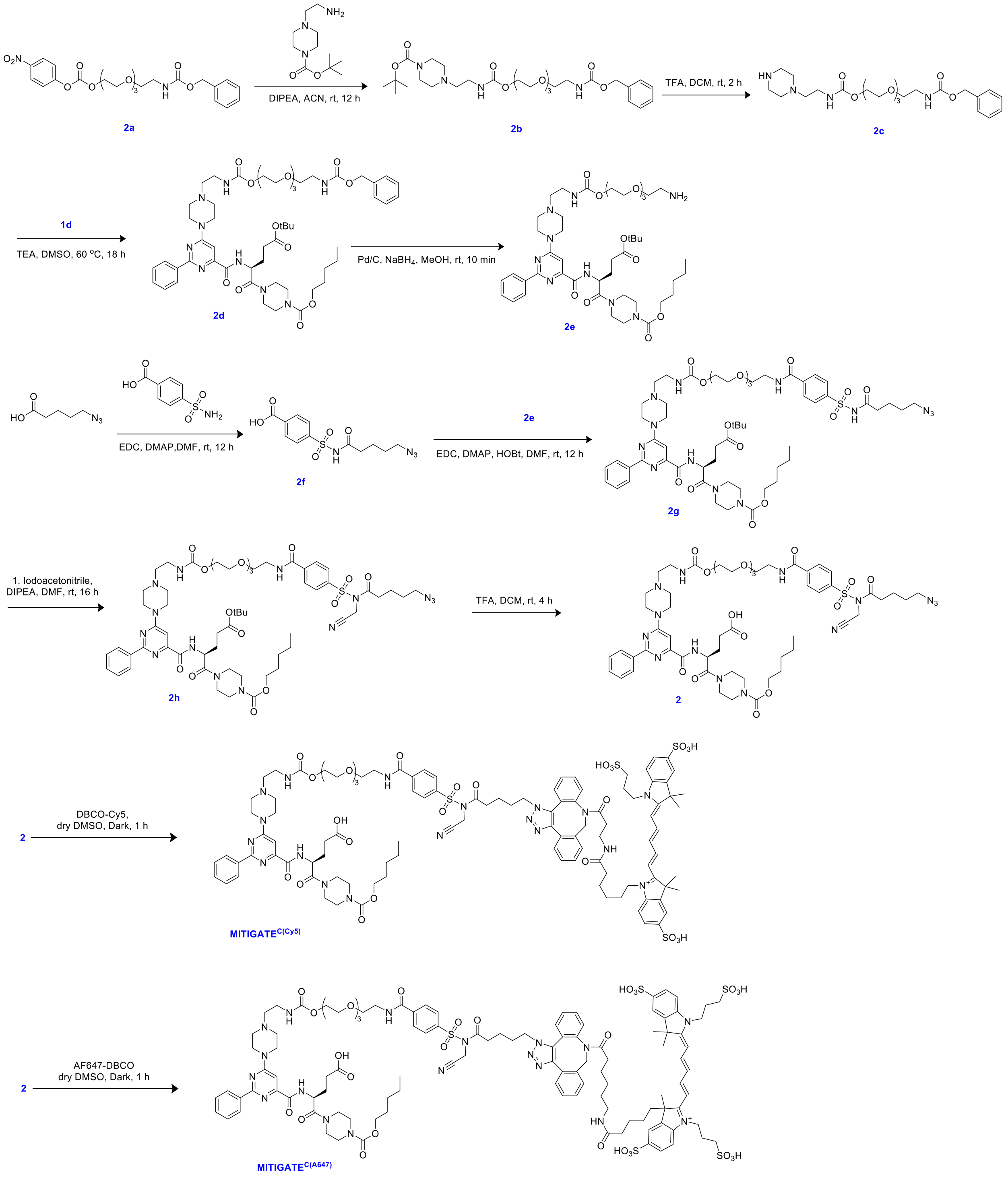

**Scheme 2:** Scheme for the synthesis of **MITIGATE**$^{C(Cy5)}$ and **MITIGATE**$^{C(AF647)}$

**2b**: A flame dried 50 mL round bottom flask equipped with a magnetic bead under nitrogen atmosphere was loaded with **2a** (1.906 g, 3.87mmoles) and was dissolved in 6 mL dry ACN. DIPEA (1.01 mL, 5.81 mmoles) was added to the solution and stirred for 2 minutes followed by addition of tert-Butyl 4-(2-aminoethyl) piperazine-1-carboxylate (1.07 g, 4.65 mmoles). The reaction mixture was stirred at room temperature for 3 hours. The reaction progress was monitored using TLC. Upon completion of the reaction, solvent was removed under reduced pressure. DCM was added to the crude mixture and the organic phase was washed with brine and dried over $Na_2SO_4$. The crude mixture was purified using silica gel column chromatography in 10% MeOH in DCM to afford desired product as colourless oil (Yield = 95 %). $R_f$ = 0.55 (10% MeOH in DCM); $^1H$ NMR (300 MHz, $CDCl_3$) δ (PPM): 7.42 – 7.31 (m, 5H), 5.44 (s, 1H), 5.31

(s, 1H), 5.11 (s, 2H), 4.22 (t, J = 4.6 Hz, 2H), 3.64 (d, J = 5.7 Hz, 10H), 3.58 (t, J = 5.2 Hz, 2H), 3.41 (q, J = 6.4 Hz, 6H), 3.29 (q, J = 5.8 Hz, 2H), 2.47 (t, J = 6.0 Hz, 2H), 2.39 (t, J = 5.0 Hz, 4H), 1.47 (s, 9H). $^{13}$C NMR (300 MHz, $CDCl_3$) δ (PPM): 156.51, 156.35, 154.70, 136.63, 128.50, 128.09, 79.70, 70.56, 70.47, 70.27, 70.07, 69.69, 66.65, 63.88, 57.03, 53.44, 52.69, 40.90, 37.44, 29.70, 28.42

**2c:** A 50 mL round bottom flask equipped with a magnetic bead was loaded with **2c** (2 g, 3.44 mmoles) and was dissolved in 5 mL DCM. 0.5 mL of TFA was added to the solution and the mixture was stirred for 1 hour. The reaction progress was monitored using TLC. Upon completion of the reaction, the reaction mixture was neutralized using 1N NaOH solution. DCM was added to the crude mixture. The organic phase was washed with brine, dried over $Na_2SO_4$ and evaporated to dryness to afford the desired product as colourless oil in quantitative yield. **2c** acquired was used for the next step without further purification.

**2d:** A 10 mL round bottom flask equipped with a magnetic bead was loaded with **1d** (200 mg, 0.152 mmoles) was dissolved in 2 mL DMSO. TEA (0.027 mL, 0.197 mmoles) was added to the solution, followed by **2c** (76 mg, 0.1580 mmoles). The reaction mixture was stirred at 60 °C for 18 hours. The reaction progress was monitored using TLC. Upon completion of the reaction, solvent was removed under reduced pressure. DCM was added to the crude mixture. The organic phase was washed with brine, dried over $Na_2SO_4$ and evaporated to dryness to afford the desired product as colourless oil (Yield = 98 %). $R_f$ = 0.55 (3% MeOH in DCM). Calculated mass: 1047.56 g/mol, experimental mass: 1047.58 g/mol $[M]^+$. $^1$H NMR (300 MHz, $CDCl_3$) δ (ppm) 8.99 (d, *J* = 8.5 Hz, 1H), 8.47 (s, 2H), 7.55 – 7.45 (m, 3H), 7.44 – 7.31 (m, 5H), 5.46 (s, 1H), 5.38 (s, 1H), 5.32 (s, 1H), 5.21 (m, 1H), 5.12 (s, 2H), 4.24 (s, 2H), 4.12 (t, *J* = 6.7 Hz, 2H), 3.85 (s, 3H), 3.72 (m, 3H), 3.65 (m, 11H), 3.58 (m, 4H), 3.41 (q, *J* = 5.3 Hz, 2H), 3.34 (d, *J* = 5.8 Hz, 2H), 2.56 (m, 5H), 2.40 (m, 2H), 2.07 – 1.87 (m, 3H), 1.66 (m, 2H), 1.47 (s, 9H), 1.42 – 1.31 (m, 4H), 0.99 – 0.88 (m, 3H). $^{13}$C NMR (300 MHz, $CDCl_3$) δ (ppm): 172.14, 169.87, 164.07, 163.16, 162.99, 156.52, 156.40, 155.64, 155.43, 137.46, 136.63, 130.65, 128.51, 128.36, 128.24, 128.10, 98.68, 80.76, 70.57, 70.49, 70.26, 70.09, 69.70, 66.65, 66.02, 63.04, 57.04, 52.54, 48.27, 45.39, 43.95, 42.07, 40.92, 37.50, 30.77, 29.70, 28.66, 28.35, 28.11, 22.36, 13.99

**2e:** A flame dried 10 mL round bottom flask equipped with a magnetic bead under nitrogen atmosphere was loaded with **2d** (120 mg, 0.114 mmoles). 1.5 mL MeOH was added to the RB followed by palladium on activated charcoal (Pd-C) (12 mg, 10 w% of 120 mg). The nitrogen balloon was replaced with an empty balloon, followed by addition of $NaBH_4$ (5 mg, 0.114 mmoles). The RB was tightly sealed and the reaction mixture was allowed to stir for 15 min. The reaction progress was monitored using TLC. Upon 60 % completion of the reaction, reaction mixture was filtered through celite and filtrate was evaporated to dryness. DCM was added to the crude mixture. The organic phase was washed with brine, dried over $Na_2SO_4$ and evaporated to dryness to afford the desired product as off-white oil. The compound was used for the next step without further purification.

**2f:** A flame dried 10 mL round bottom flask equipped with a magnetic bead under nitrogen atmosphere was loaded with azidovaleric acid (67 mg, 0.46 mmoles) and was dissolved in 0.5 mL dry DMF. DIPEA (0.04 mL, 0.234 mmoles) was added to the solution followed by EDC.HCl (36 mg, 0.234 mmoles). The resultant mixture was stirred for 1 hour at room temperature. The reaction mixture was added to another round bottom flask containing 4-sulfamoylbenzoic acid (36 mg, 0.234 mmoles) and DMAP (7 mg, 0.0234 mmoles) dissolved in 0.5 mL DMF. The reaction mixture was stirred at room temperature for 12 hours. The reaction progress was monitored using TLC. Upon completion of the reaction, solvent was removed under reduced pressure. DCM was added to the crude mixture. The organic phase was washed with 10% HCl, dried over $Na_2SO_4$ and was evaporated to dryness to afford the desired product as white solid (Yield = 80%). $R_f$ = 0.55 (10% MeOH in DCM with a drop of AcOH). Calculated mass: 326.07 g/mol, experimental mass: 348.08 g/mol $[M+Na]^+$. $^1$H NMR (300 MHz, MeOH-$d_4$) δ (ppm): 8.19 (d, J = 8.2 Hz, 2H), 8.09 (d, J = 8.1 Hz, 2H), 3.24 (t, J = 6.5 Hz, 2H), 2.29 (t, J = 7.0 Hz, 2H), 1.65 – 1.55 (m, 2H), 1.50 (m, 2H). $^{13}$C NMR (300MHz, MeOD) δ (ppm): 172.67, 143.10, 129.61, 127.69, 50.59, 35.11, 27.66, 21.44

**2g:** A flame dried 10 mL round bottom flask equipped with a magnetic bead under nitrogen atmosphere was loaded with **2f** (45 mg, 0.138 mmoles) dissolved in 0.5 mL dry DMF. DIPEA (0.07 mL, 0.414 mmoles) was added to the solution followed by EDC.HCl (32 mg, 0.207 mmoles) and HOBt (28 mg, 0.207 mmoles). The resultant mixture was stirred at room temperature for 30 minutes followed by addition of the **2e** (126 mg, 0.138 mmoles). The reaction mixture was stirred at room temperature for 12 hours. The reaction progress was monitored using TLC. Upon completion of the reaction, solvent was removed under reduced pressure. The crude mixture was purified using preparative TLC in 6% MeOH in DCM to afford the afford the desired product as off-white waxy solid (yield = 20 %). $R_f$ = 0.3 (6% MeOH in DCM); Calculated mass: 1221.59 g/mol, experimental mass: 1222.61 g/mol $[M+H]^+$. $^1$H NMR (300 MHz, $CDCl_3$) δ (ppm): 9.01 (m, 1H), 8.47 (s, 2H), 8.09 (d, *J* = 8.1 Hz, 1H), 7.99 (d, *J* = 8.2 Hz, 1H), 7.53 – 7.43 (m, 4H), 7.01 (s, 1H), 5.46 (s, 1H), 5.22 (s, 1H), 5.06 (m, 1H), 5.00 – 4.90 (m, 1H), 4.33 – 4.17 (m, 3H), 4.11 (m, 4H), 3.87 (s, 5H), 3.81 – 3.60 (m, 25H), 3.57 (m, 6H), 3.39 (s, 3H), 3.24 (t, *J* = 6.5 Hz, 2H), 2.97 – 2.81 (m, 4H), 2.73 – 2.52 (m, 9H), 2.46 – 2.30 (m, 5H), 2.19 (s, 2H), 2.11 – 1.94 (m, 4H), 1.73 – 1.53 (m, 9H), 1.37 (s, 9H), 0.94 – 0.88 (m, 8H). $^{13}$C NMR (300 MHz, $CDCl_3$) δ (ppm): 171.14, 168.91, 163.13, 161.97, 154.42, 140.73, 136.30, 134.85, 129.70, 127.34, 127.21, 126.80, 123,73, 113.04, 97.72, 79.83, 69.50, 69.37, 69.10, 68.54, 65.02, 51.44, 49.95, 47.37, 44.39, 41.08, 35.48, 34.59, 33.27, 32.79, 30.90, 30.00, 29.73, 29.28, 28.66, 28.48, 28.33, 28.24, 28.13, 27.93, 27.62, 27.62, 27.22, 27.08, 24.89, 21.66, 21.32, 20.47, 13.10, 12.95

**2h:** A flame dried 10 mL round bottom flask equipped with a magnetic bead under nitrogen atmosphere was loaded with **2g** (7.5 mg, 6 mmoles) and was dissolved in 0.5 mL dry DMF. DIPEA (5.4 mL, 31 mmoles) was added to the solution followed by iodoacetonitrile (3 mL, 31 mmoles). The reaction mixture was allowed to stir for 16 hours at room temperature. The reaction progress was monitored using TLC. Upon completion of the reaction, solvent was removed under reduced pressure. The crude mixture was purified using preparative TLC in 5% MeOH in DCM to afford the desired product as off-white waxy solid (yield = 91 %). $R_f$ = 0.4 (5% MeOH in DCM); Calculated mass: 1260.60 g/mol, experimental mass: 1261.60 g/mol $[M+H]^+$. $^1$H NMR (300 MHz, $CDCl_3$) δ (ppm): 9.00 (m, 1H), 8.48 (m, 2H), 8.14 (d, *J* = 8.4 Hz, 2H), 8.07 (d, *J* = 8.5 Hz, 2H), 7.53 – 7.47 (m, 3H), 7.01 (s, 2H), 5.09 (s, 1H), 4.98 (m, 1H), 4.22 (s, 2H), 4.11 (m, 4H), 3.87 (s, 4H), 3.80 – 3.41 (m, 28H), 3.35 (s, 2H), 3.27 (t, *J* = 6.4 Hz, 2H), 2.94 – 2.84 (m, 2H), 2.73 (t, *J* = 7.1 Hz, 3H), 2.67 – 2.49 (m, 8H), 2.36 (t, *J* = 7.4 Hz, 5H), 2.10 – 2.00 (m, 3H), 1.63 (m, 9H), 1.47 (s, 9H), 0.90 (m, 22H). $^{13}$C NMR (300 MHz, $CDCl_3$) δ (ppm): 176.55, 171.11, 170.31, 168.85, 162.14, 154.41, 146.04, 138.25, 134.84, 130.13, 129.66, 127.82, 127.34, 127.21, 126.78, 123.74, 123.43, 122.95, 118.07, 113.04, 97.68, 79.68, 69.38, 69.17, 68.64, 65.00, 63.62, 51.49, 49.97, 47.27, 44.34, 42.87, 39.18, 35.50, 34.46, 33.84, 33.48, 33.27, 32.79, 32.10, 31.75, 30.90, 30.59, 30.41, 30.00, 29.73, 29.27, 29.16, 29.01, 28.67, 28.49, 28.42, 28.33, 28.24, 28.13, 28.07, 27.92, 27.62, 27.08, 26.89, 26.07, 25.71, 24.88, 23.72, 21.67, 21.33, 20.61, 13.09, 12.96.

**2:** A 10 mL round bottom flask equipped with a magnetic bead was loaded with **2h** (3.5 mg, 2.7 mmoles) and was dissolved in 2 mL DCM. 0.4 mL of TFA was added to the solution and the mixture was stirred for 4 hours. The reaction progress was monitored using TLC. Upon completion of the reaction, the reaction mixture was neutralized using 1N NaOH solution. DCM was added to the crude mixture and the organic phase was washed with brine and dried over $Na_2SO_4$ and evaporated to dryness to afford the desired product as colourless waxy solid (Yield = 90%). **2** was used for the next step without further purification. Calculated mass: 1204.53 g/mol, experimental mass: 1205.54 g/mol $[M+H]^+$. $^1$H NMR (300 MHz, $CDCl_3$) δ (ppm): 9.08 (m, 1H), 8.43 (m, 2H), 8.15 (m, 2H), 8.04 (d, *J* = 7.9 Hz, 2H), 7.52 (m, 5H), 5.09 (s, 4H), 4.80 (s, 4H), 4.20 (s, 5H), 4.16 – 4.00 (m, 11H), 3.54-3.71 (m, 43H), 3.26 (t, *J* = 6.5 Hz, 5H), 2.89 (dd, *J* = 9.2, 6.7 Hz, 5H), 2.62 (t, *J* = 8.0 Hz, 6H), 2.52 (s, 3H), 2.37 (t, *J* = 7.5 Hz, 6H), 2.06 (m, 7H), 1.72 – 1.58 (m, 22H), 0.90 (s, 25H). $^{13}$C NMR (300 MHz, $CDCl_3$) δ (ppm): 177.01, 172.37, 151.10, 146.06, 138.26, 134.84, 130.12, 127.95, 127.55, 127.25, 126.72, 123.74, 123.44, 122.95, 118.08, 113.04, 69.42, 69.30, 68.98, 62.62, 49.97, 36.08, 35.49, 34.47, 33.84, 33.49, 33.27, 32.80, 32.21, 31.71, 30.90, 30.40, 30.00, 29.28, 29.16, 28.67, 28.49, 28.34, 28.23, 28.13, 28.05, 27.93, 27.62, 27.03, 26.86, 26.06, 25.68, 24.89, 24.47, 23.70, 21.67, 21.31, 20.59, 13.09, 12.95

**MITIGATE$^{C(Cy5)}$:** Sulfo-DBCO-Cy5 (0.133 µmoles) in 1 µL dry DMSO (molecular biology grade) was mixed with **2** (0.133 µmoles) in 1 µL dry DMSO in the dark for 1 hour. Product formation was verified using mass spectrometry. Calculated m/z: 2213.85 g/mol, experimental mass: 2213.83 g/mol (deconvoluted for resolved isotope) [M] +.

**MITIGATE$^{C(AF647)}$:** AF647-DBCO (0.133 µmoles) in 1 µL dry DMSO (molecular biology grade) was mixed with **2** (0.133 µmoles) in 1 µL dry DMSO in the dark for 1 hour. Product formation was verified using mass spectrometry. Calculated m/z: 2349.87 g/mol, experimental mass: 2349.85 g/mol (deconvoluted for resolved isotope) $[M]^+$.

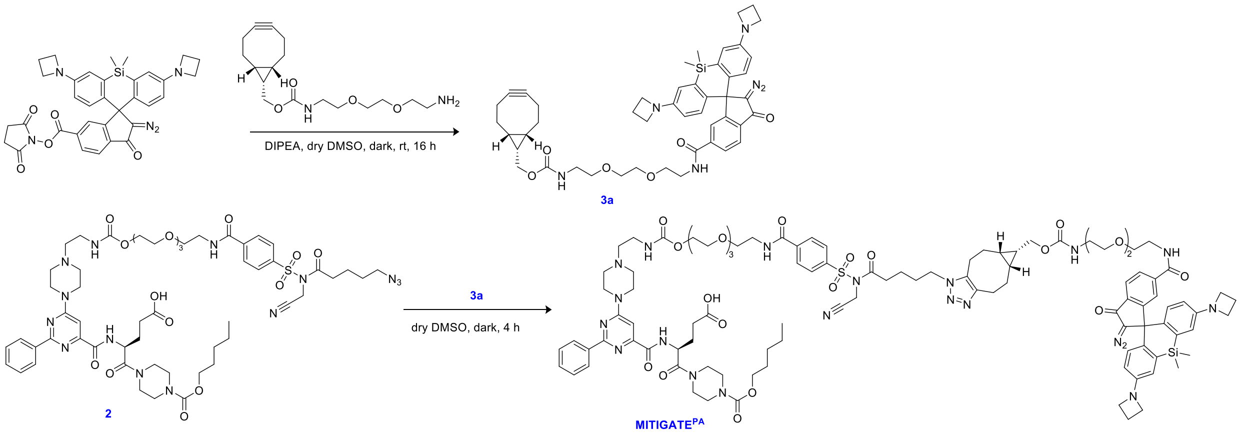


**Scheme 3:** Scheme for the synthesis of **3a**

**3a:** An amber glass vial equipped with a magnetic bead was loaded with Photoactivatable (PA) Janelia Fluor 646 NHS ester (0.3 mg, 0.00049 mmol) and ((1R,8S,9s)-bicyclo[6.1.0]non-4-yn-9-yl)methyl (2-(2-(2-aminoethoxy)ethoxy)ethyl)carbamate (0.159 mg, 0.00049 mmol) were dissolved in 100 µL dry DMSO. DIPEA (0.26 µL, 0.00147 mmol) was added to the reaction mixture and was allowed to stir overnight in the dark. After completion of the reaction, the mixture was freeze dried to afford the desired product as lyophilised powder. Product formation was verified using mass spectrometry. Calculated mass: 826.39 g/mol, experimental mass: 826.39 g/mol $[M]^+$.

**MITIGATE$^{PA}$: 3a** (0.7 mg, 0.8 µmoles) in 1 µL dry DMSO (molecular biology grade) was mixed with **2** (0.1 mg, 0.8 µmoles) in 1 µL dry DMSO in the dark for 4 hours.

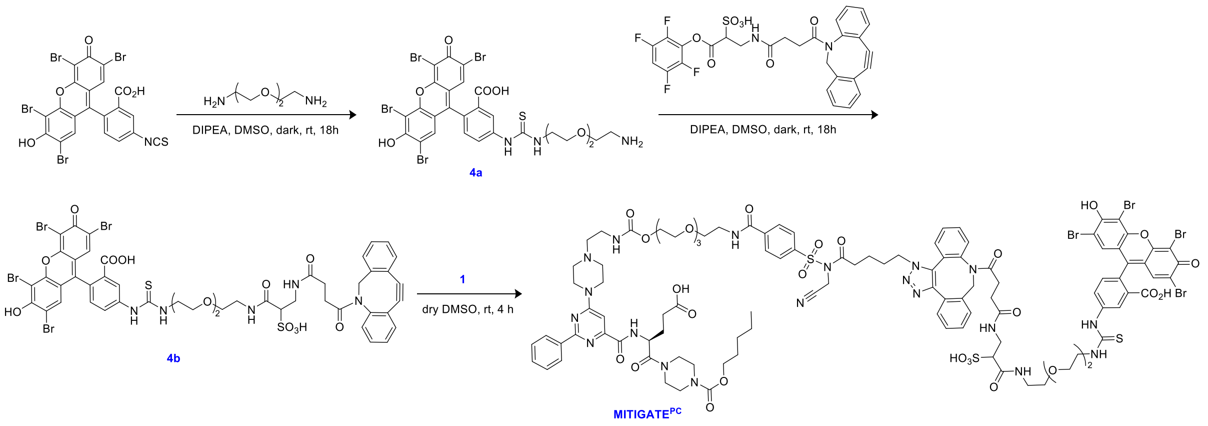


**Scheme 4:** Scheme for the synthesis of **MITIGATE$^{PC}$**

**4a:** An amber glass vial equipped with a magnetic bead was loaded with Eosin-5-isothiocyanate (5 mg, 0.0071 mmol) and 2, 2'-(Ethylenedioxy) bis(ethyldiamine) (26.3 mg, 0.1775 mmol) were dissolved in 100 µL dry DMSO. DIPEA (3.7 µL, 0.0213 mmol) was added to the reaction mixture and was allowed to stir for overnight in the dark. After completion of the reaction, reaction mixture was freeze dried to afford of desired product as orange lyophilised powder. Product formation was verified using mass spectrometry and was used for the next step without further purification. Calculated m/z: 852.79 g/mol, experimental m/z: 852.79 g/mol $[M]^+$.

**4b:** An amber glass vial equipped with a magnetic bead was loaded with **4a** (1.4 mg, 0.0016 mmol) and Sulfo-DBCO-TFP Ester (2 mg, 0.0032 mmol) were dissolved in 60 µL dry DMSO. DIPEA (1.6 µL, 0.0096 mmol) was added to the reaction mixture and was allowed to stir for overnight in the dark. After completion of the reaction, desired product was precipitated in ACN and was washed twice with DCM. The solid obtained was then was freeze dried to afford of desired product as orange lyophilised powder. Product formation was verified using mass spectrometry and was used for the next step without further purification. Calculated m/z: 1290.88 g/mol, experimental m/z: 1291.91 g/mol $[M+H]^+$

**MITIGATE$^{PC}$: 4b** (0.64 mg, 0.5 μmoles) in 1 μL dry DMSO (molecular biology grade) was mixed with **2** (0.6 mg, 0.5 μmoles) in 1 μL dry DMSO in the dark for 4 hours. Product formation was verified using mass spectrometry. Calculated m/z: 2498.43 g/mol (80%), experimental m/z: 2498.43 g/mol $[M]^+$

**$^{1}$H-NMR, $^{13}$C-NMR, mass spectra and HPLC chromatograms**

**Note:** Mass spectrum is provided for the asterisk marked peak in the chromatogram of each intermediate.

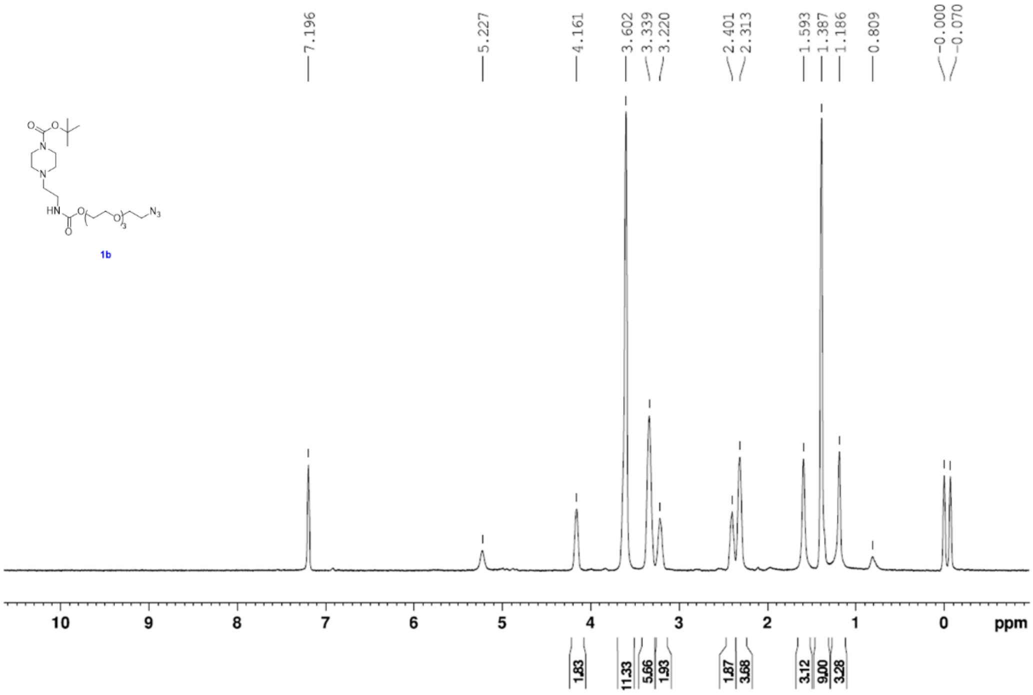


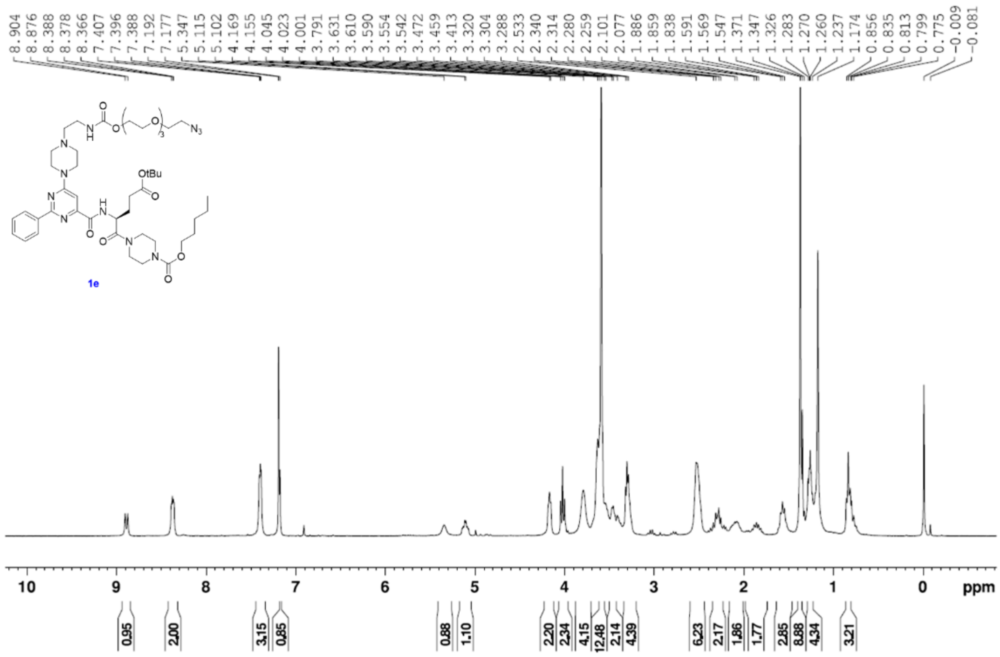

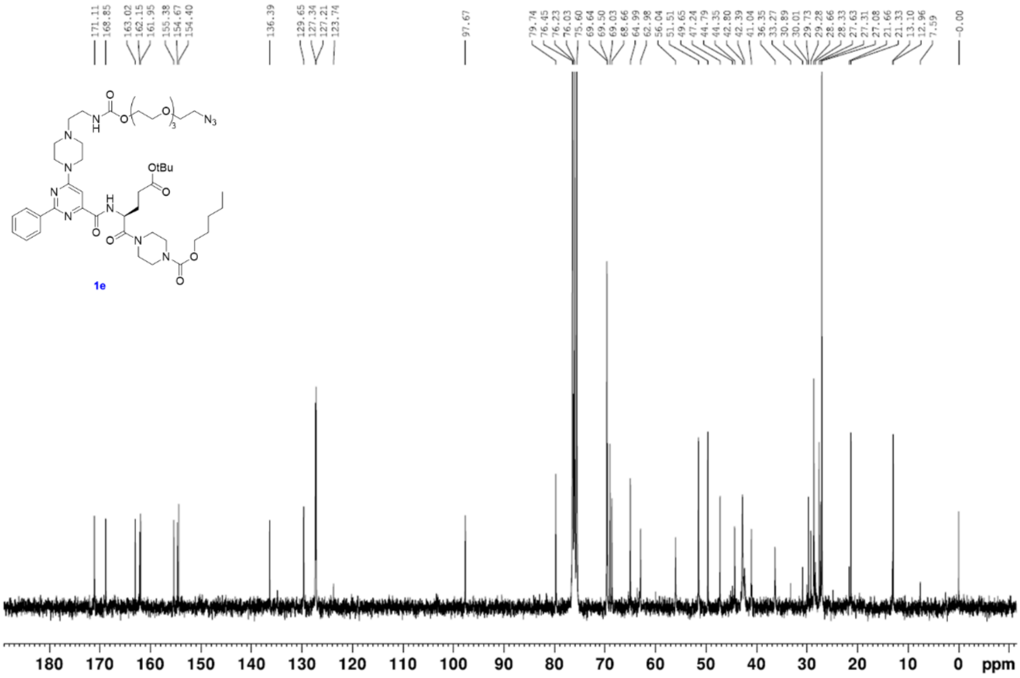
171.11
168.85
163.02
162.15
161.95
155.38
154.67
154.40
136.39
129.65
127.34
127.21
123.74
97.67
79.74
76.45
76.23
76.03
75.60
69.64
69.50
69.03
68.66
64.99
62.98
56.04
51.51
49.65
47.24
44.79
44.35
42.80
42.39
41.04
36.35
33.27
30.89
30.01
29.73
29.28
28.66
28.33
27.63
27.31
27.08
21.66
21.33
13.10
12.96
7.59
-0.00
OtBu
$N_3$
1e
180 170 160 150 140 130 120 110 100 90 80 70 60 50 40 30 20 10 0 ppm

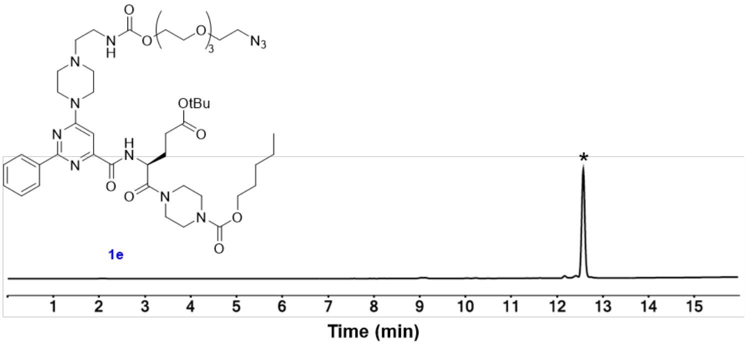
OtBu
$N_3$
1e
*
1 2 3 4 5 6 7 8 9 10 11 12 13 14 15
Time (min)

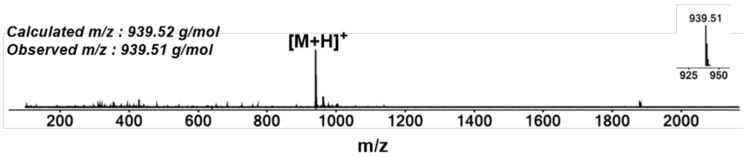
Calculated m/z : 939.52 g/mol
Observed m/z : 939.51 g/mol
[M+H]+
939.51
925 950
200 400 600 800 1000 1200 1400 1600 1800 2000
m/z

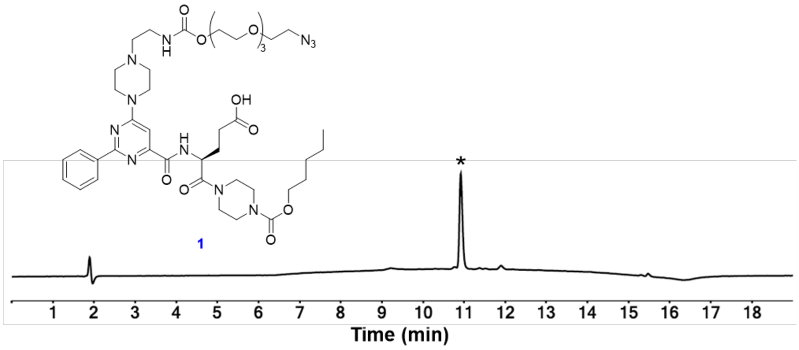
OH
$N_3$
1
*
1 2 3 4 5 6 7 8 9 10 11 12 13 14 15 16 17 18
Time (min)

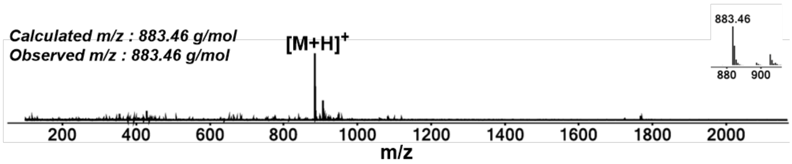
Calculated m/z : 883.46 g/mol
Observed m/z : 883.46 g/mol
[M+H]+
883.46
880 900
200 400 600 800 1000 1200 1400 1600 1800 2000
m/z

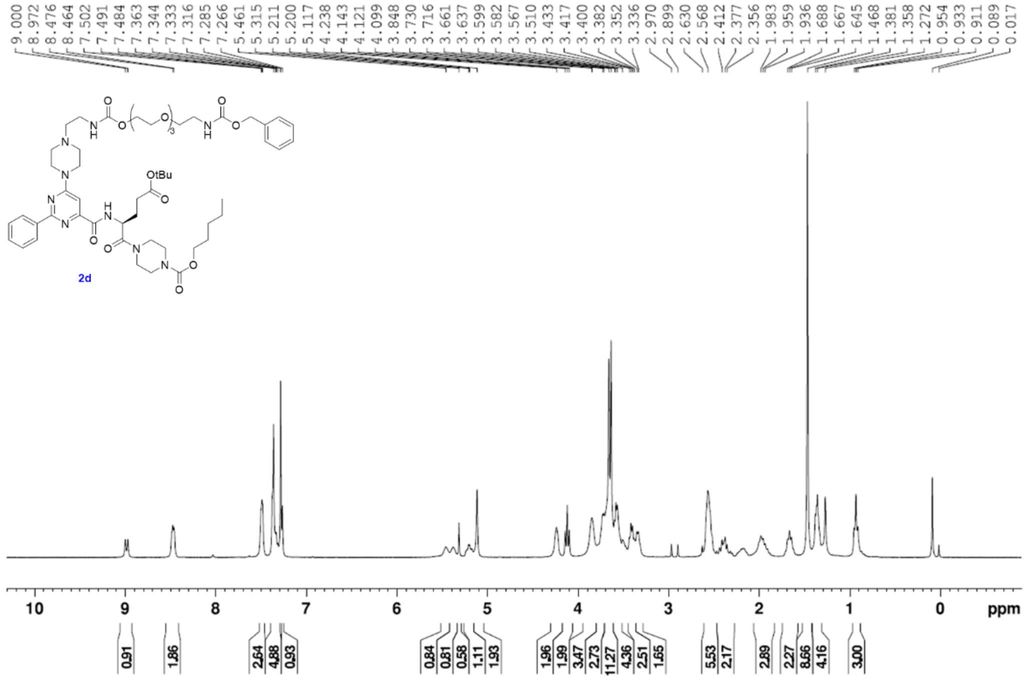
9.000 8.972 8.464 8.476 7.502 7.491 7.484 7.363 7.344 7.333 7.316 7.285 7.266 5.461 5.315 5.211 5.200 5.117 4.238 4.143 4.121 4.099 3.848 3.730 3.716 3.661 3.637 3.599 3.582 3.567 3.510 3.433 3.417 3.400 3.382 3.352 3.336 2.970 2.899 2.630 2.568 2.412 2.377 2.356 1.983 1.959 1.936 1.688 1.667 1.645 1.468 1.381 1.358 1.272 0.954 0.933 0.911 0.089 0.017
OtBu
2d
10 9 8 7 6 5 4 3 2 1 0 ppm
0.91 1.86 2.64 4.88 0.93 0.84 0.81 0.58 1.11 1.93 1.96 1.99 3.47 2.73 11.27 4.36 2.51 1.65 5.53 2.17 2.89 2.27 8.66 4.16 3.00

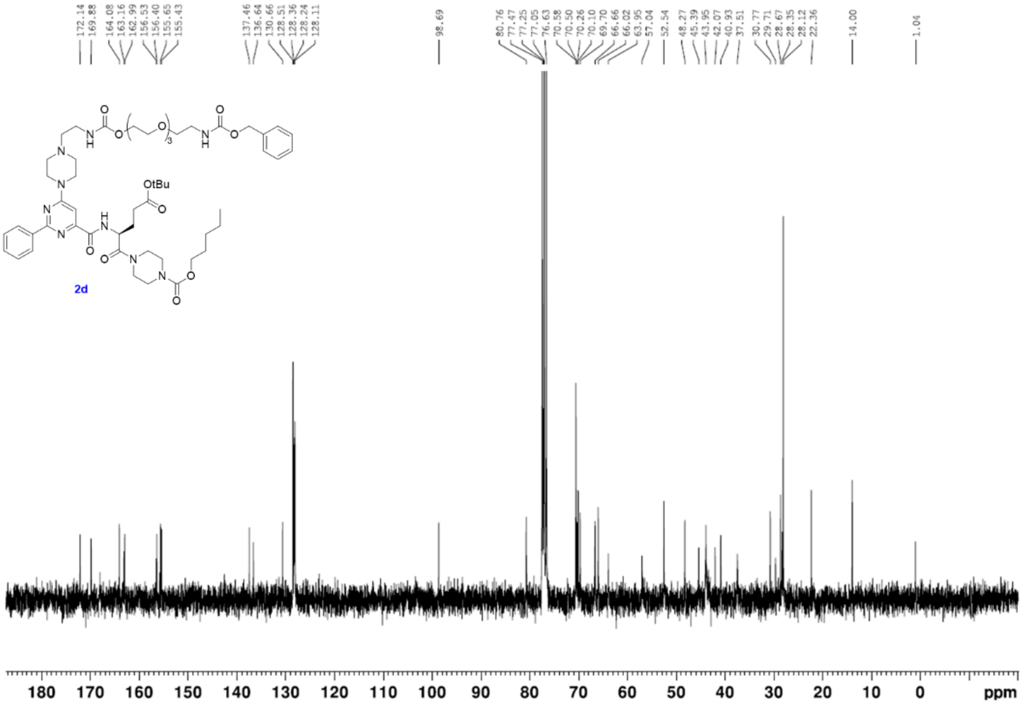
172.14 169.88 164.08 163.16 162.99 156.53 156.40 155.65 155.43 137.46 136.64 130.66 128.51 128.36 128.24 128.11 98.69 80.76 77.47 77.25 77.05 76.63 70.58 70.50 70.26 70.10 69.70 66.66 66.02 63.95 57.04 52.54 48.27 45.39 43.95 42.07 40.93 37.51 30.77 29.71 28.67 28.35 28.12 22.36 14.00 1.04
OtBu
2d
180 170 160 150 140 130 120 110 100 90 80 70 60 50 40 30 20 10 0 ppm

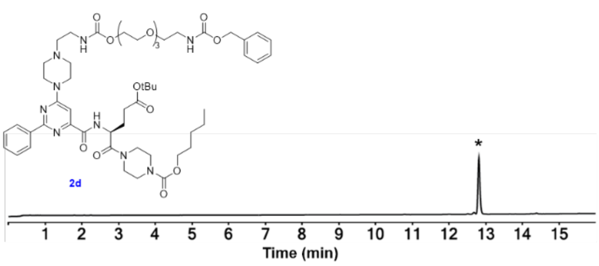
OtBu
2d
*
1 2 3 4 5 6 7 8 9 10 11 12 13 14 15
Time (min)

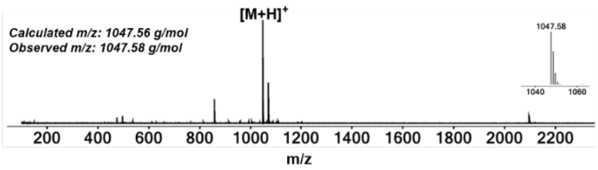
[M+H]+
Calculated m/z: 1047.56 g/mol
Observed m/z: 1047.58 g/mol
1047.58
1040 1060
200 400 600 800 1000 1200 1400 1600 1800 2000 2200
m/z

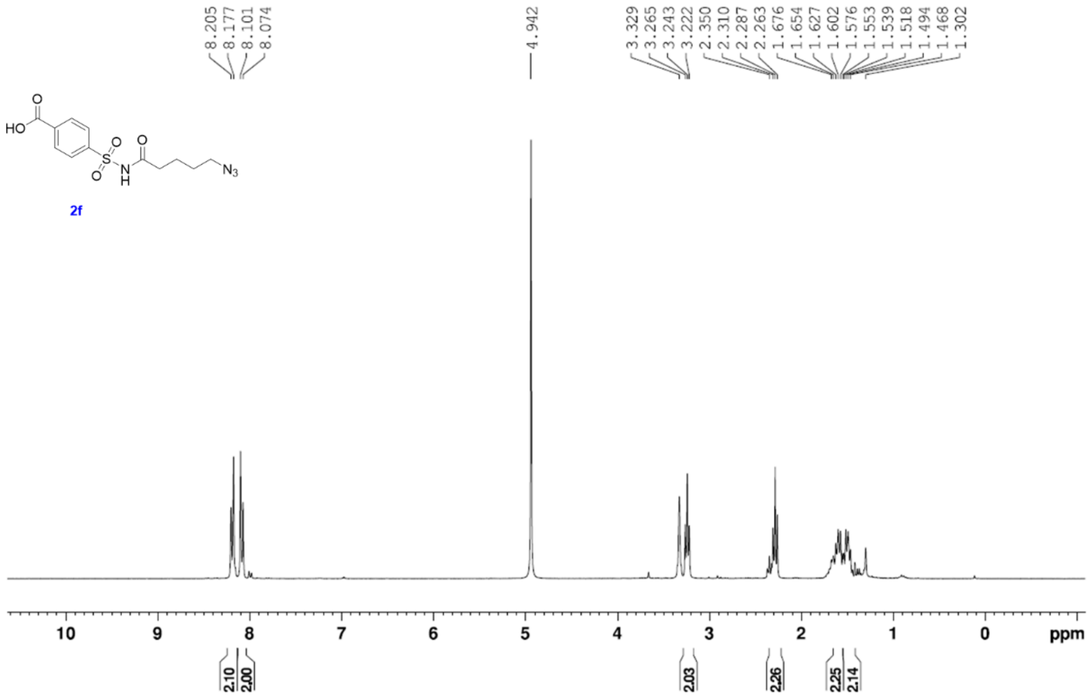
8.205
8.177
8.101
8.074
4.942
3.329
3.265
3.243
3.222
2.350
2.310
2.287
2.263
1.676
1.654
1.627
1.602
1.576
1.553
1.539
1.518
1.494
1.468
1.302
HO
O
S
O
O
N
H
O
N3
2f
10 9 8 7 6 5 4 3 2 1 0 ppm
2.10
2.00
2.03
2.26
2.25
2.14

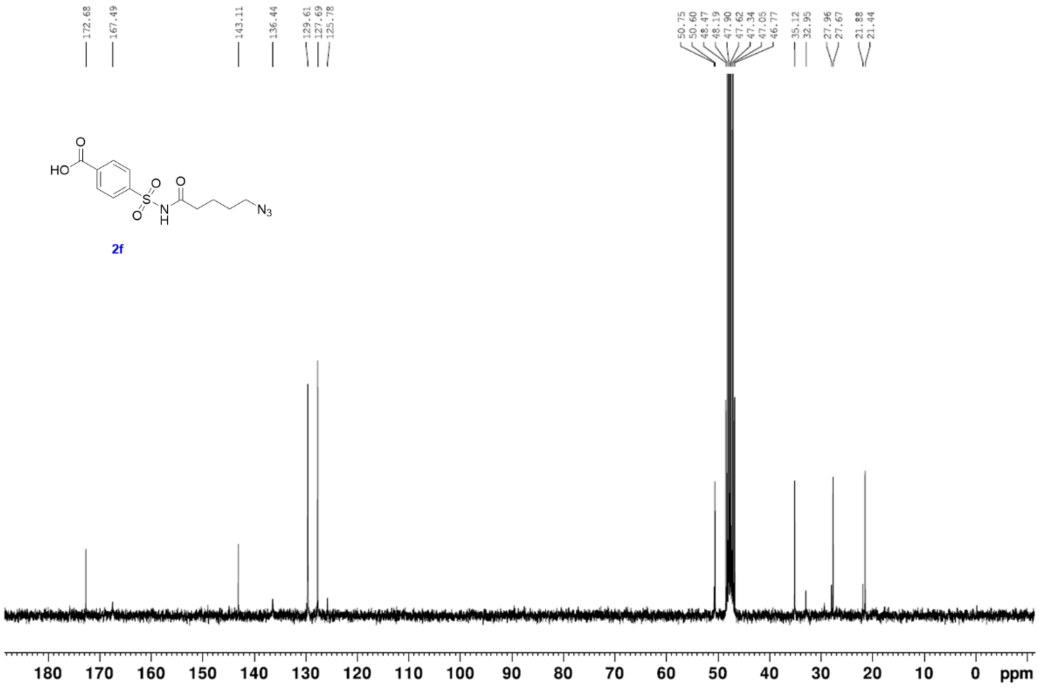
172.68
167.49
143.11
136.44
129.61
127.69
125.78
50.75
50.60
48.47
48.19
47.90
47.62
47.34
47.05
46.77
35.12
32.95
27.96
27.67
21.88
21.44
HO
O
S
O
O
N
H
O
N3
2f
180 170 160 150 140 130 120 110 100 90 80 70 60 50 40 30 20 10 0 ppm

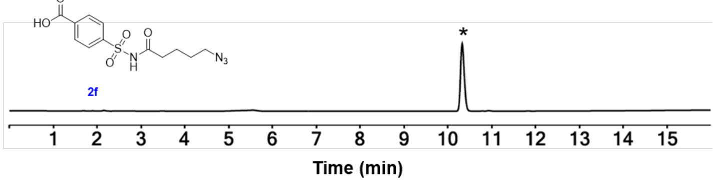
HO
O
S
O
O
N
H
O
N3
2f
*
1 2 3 4 5 6 7 8 9 10 11 12 13 14 15
Time (min)

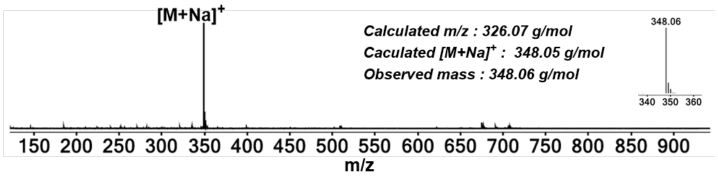
[M+Na]+
Calculated m/z : 326.07 g/mol
Caculated [M+Na]+ : 348.05 g/mol
Observed mass : 348.06 g/mol
348.06
340 350 360
150 200 250 300 350 400 450 500 550 600 650 700 750 800 850 900
m/z

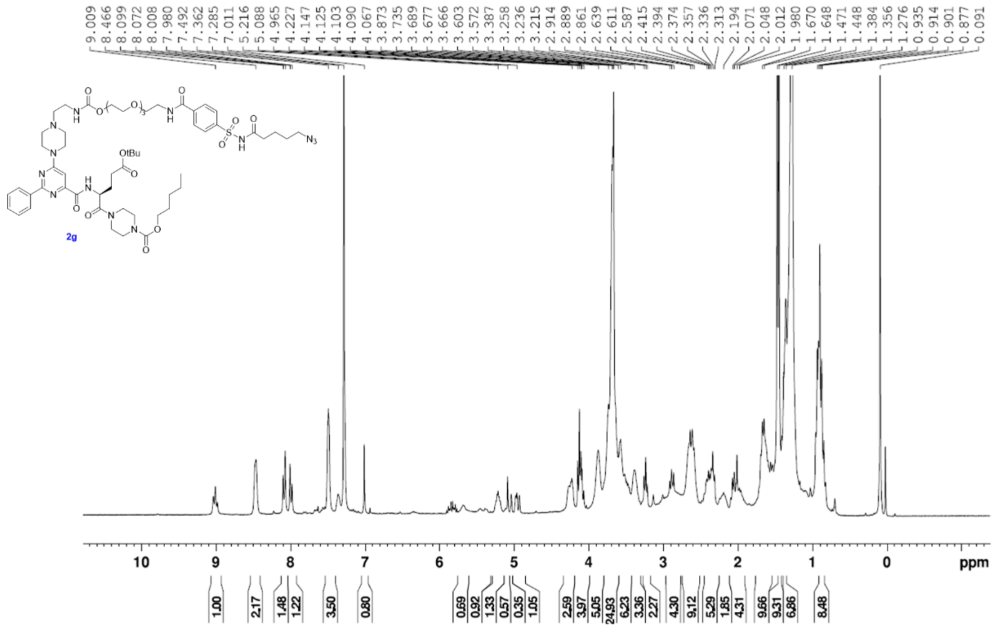
9.009 8.466 8.099 8.072 8.008 7.980 7.492 7.362 7.285 7.011 5.216 5.088 4.965 4.227 4.147 4.125 4.103 4.090 4.067 3.873 3.735 3.689 3.677 3.666 3.603 3.572 3.387 3.258 3.236 3.215 2.914 2.889 2.861 2.639 2.611 2.587 2.415 2.394 2.374 2.357 2.336 2.313 2.194 2.071 2.048 2.012 1.980 1.670 1.648 1.471 1.448 1.384 1.356 1.276 0.935 0.914 0.901 0.877 0.091
OtBu
N3
2g
10 9 8 7 6 5 4 3 2 1 0 ppm
1.00 2.17 1.48 1.22 3.50 0.80 0.69 0.92 1.33 0.57 0.35 1.05 2.59 3.97 5.05 24.93 6.23 3.36 2.27 4.30 9.12 5.29 1.85 4.31 9.66 9.31 6.86 8.48

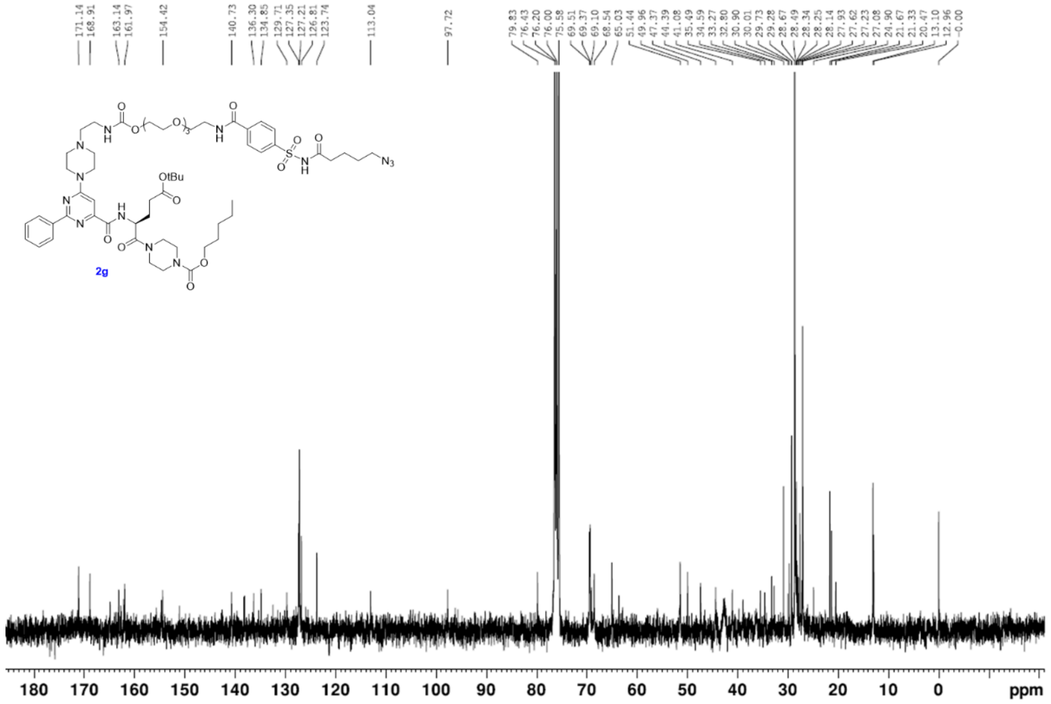
171.14 168.91 163.14 161.97 154.42 140.73 136.30 134.85 129.71 127.35 127.21 126.81 123.74 113.04 97.72 79.83 76.43 76.20 76.00 75.58 69.51 69.37 69.10 68.54 65.03 51.44 49.96 47.37 44.39 41.08 35.49 34.59 33.27 32.80 30.90 30.01 29.73 29.28 28.67 28.49 28.34 28.25 28.14 27.93 27.62 27.23 27.08 24.90 21.67 21.33 20.47 13.10 12.96 -0.00
OtBu
N3
2g
180 170 160 150 140 130 120 110 100 90 80 70 60 50 40 30 20 10 0 ppm

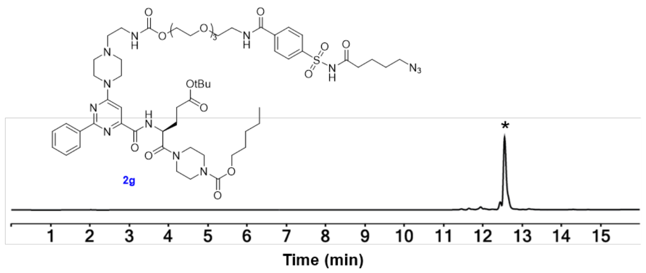
OtBu
N3
2g
*
1 2 3 4 5 6 7 8 9 10 11 12 13 14 15
Time (min)

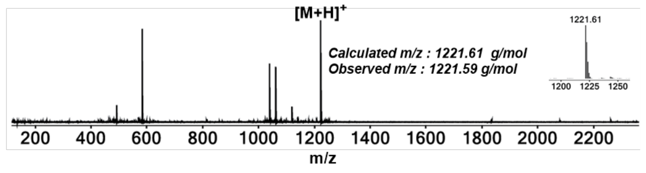
[M+H]+
Calculated m/z : 1221.61 g/mol
Observed m/z : 1221.59 g/mol
1221.61
1200 1225 1250
200 400 600 800 1000 1200 1400 1600 1800 2000 2200
m/z

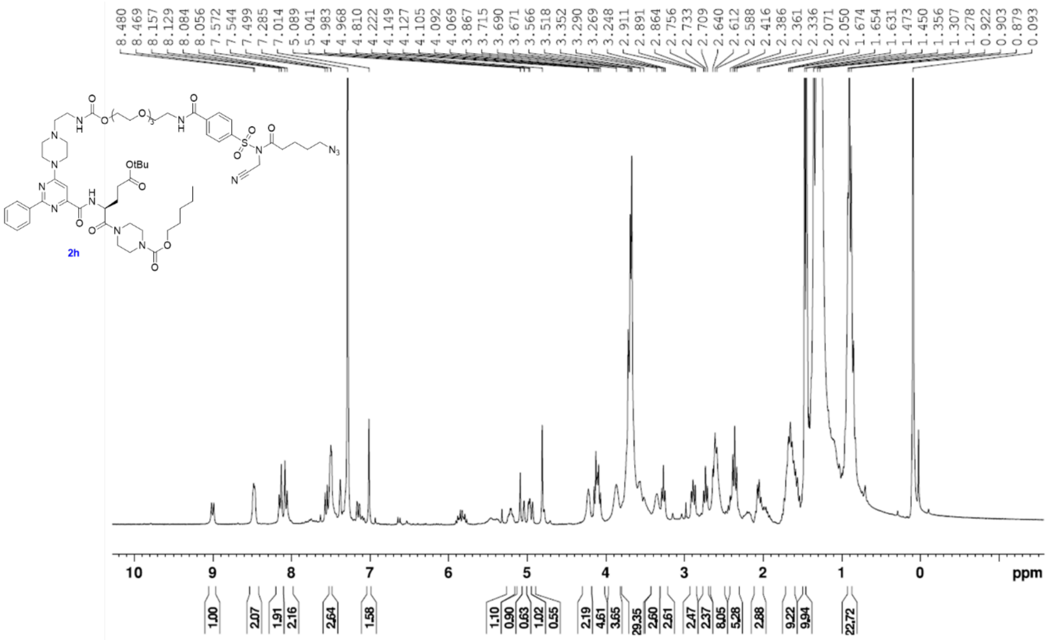
8.480 8.469 8.157 8.129 8.084 8.056 7.572 7.544 7.499 7.285 7.014 5.089 5.041 4.983 4.968 4.810 4.222 4.149 4.127 4.105 4.092 4.069 3.867 3.715 3.690 3.671 3.566 3.518 3.352 3.290 3.269 3.248 2.911 2.891 2.864 2.756 2.733 2.709 2.640 2.612 2.588 2.416 2.386 2.361 2.336 2.071 2.050 1.674 1.654 1.631 1.473 1.450 1.356 1.307 1.278 0.922 0.903 0.879 0.093
2h
10 9 8 7 6 5 4 3 2 1 0 ppm
1.00 2.07 1.91 2.16 2.64 1.58 1.10 0.90 0.63 1.02 0.55 2.19 4.61 3.65 29.35 2.60 2.61 2.47 2.37 8.05 5.28 2.88 9.22 9.94 22.72

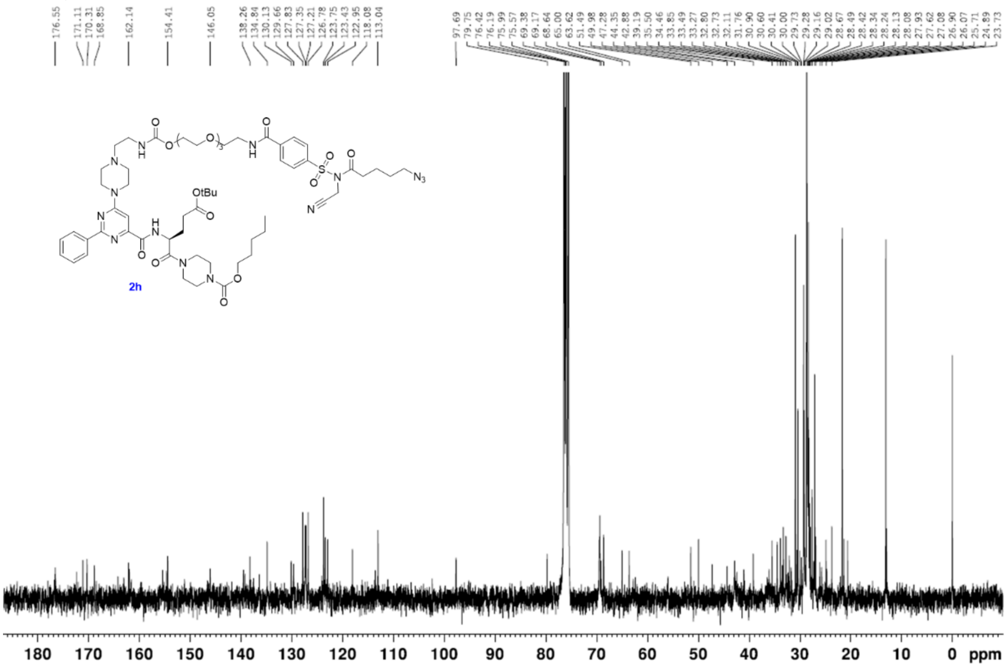
176.55 171.11 170.31 168.85 162.14 154.41 146.05 138.26 134.84 130.13 129.66 127.83 127.35 127.21 126.78 123.75 123.43 122.95 118.08 113.04 97.69 79.75 76.42 76.19 75.99 75.57 69.38 69.17 68.64 65.00 63.62 51.49 49.98 47.28 44.35 42.88 39.19 35.50 34.46 33.85 33.49 33.27 32.80 32.73 32.11 31.76 30.90 30.60 30.41 30.00 29.73 29.28 29.16 29.02 28.67 28.49 28.42 28.34 28.24 28.13 28.08 27.93 27.62 27.08 26.90 26.07 25.71 24.89 23.73
2h
180 170 160 150 140 130 120 110 100 90 80 70 60 50 40 30 20 10 0 ppm

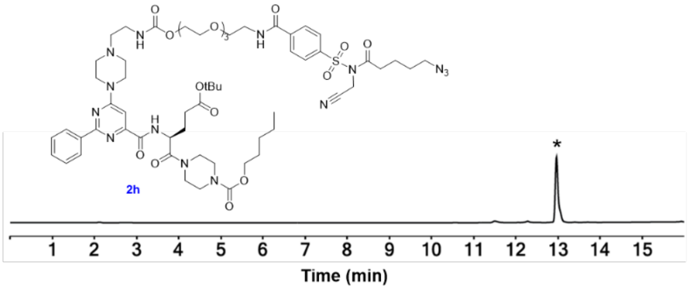
2h
*
1 2 3 4 5 6 7 8 9 10 11 12 13 14 15
Time (min)

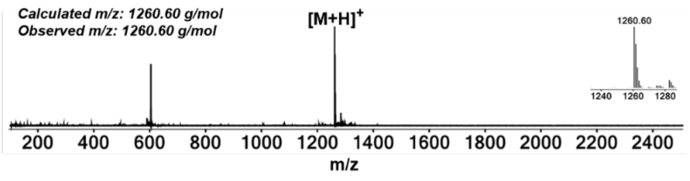
Calculated m/z: 1260.60 g/mol
Observed m/z: 1260.60 g/mol
[M+H]+
1260.60
1240 1260 1280
200 400 600 800 1000 1200 1400 1600 1800 2000 2200 2400
m/z

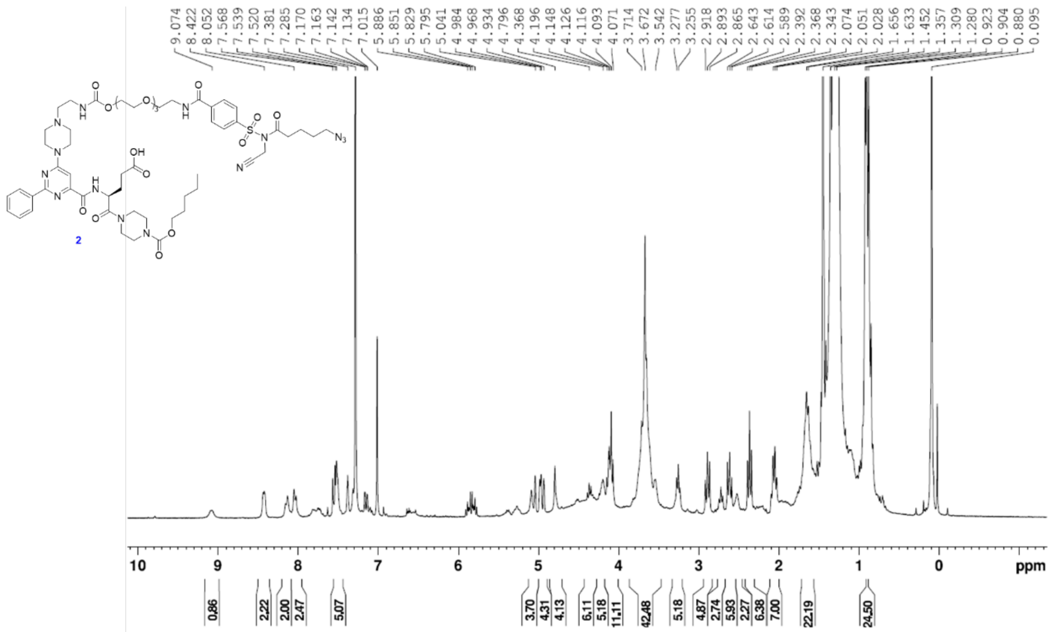

2
ppm


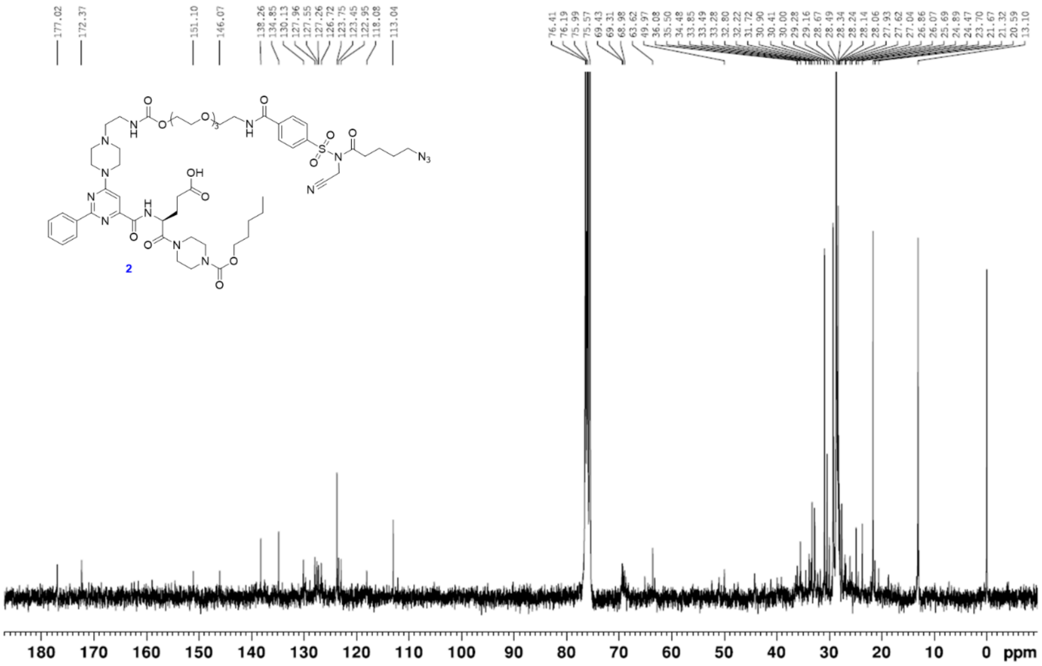

2
ppm


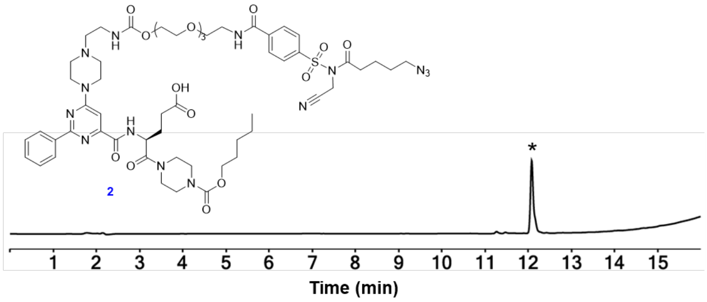

2
*
Time (min)


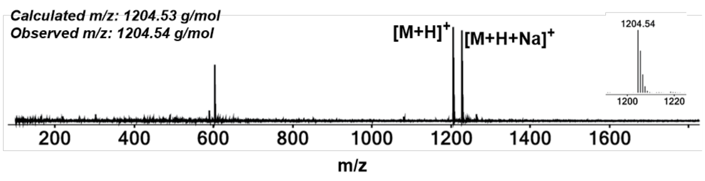

Calculated m/z: 1204.53 g/mol
Observed m/z: 1204.54 g/mol
[M+H]+
[M+H+Na]+
1204.54
m/z

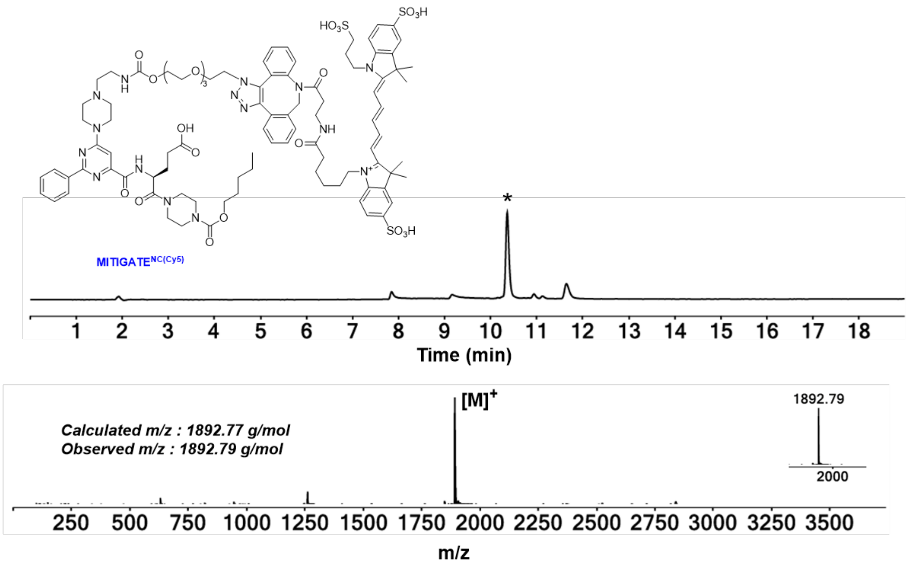
MITIGATE^NC(Cy5)
Time (min)
Calculated m/z : 1892.77 g/mol
Observed m/z : 1892.79 g/mol
[M]+
1892.79
2000
m/z

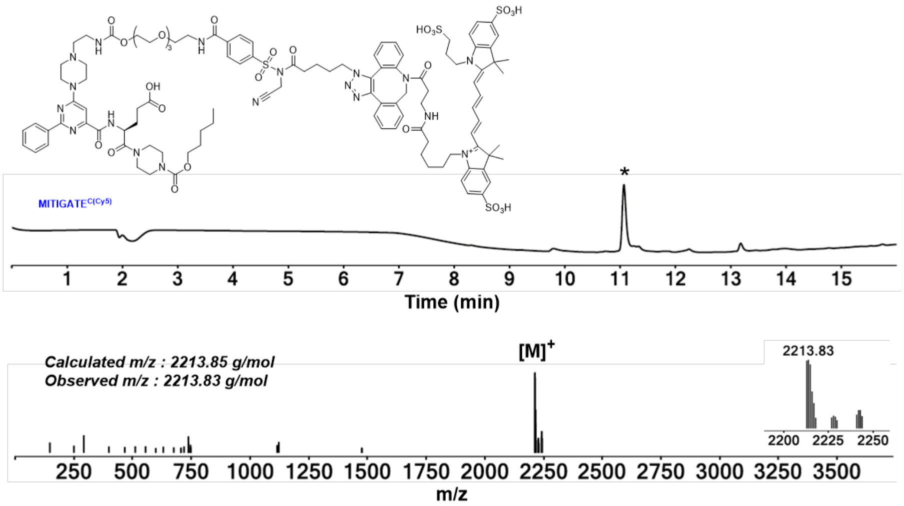
MITIGATE^C(Cy5)
Time (min)
Calculated m/z : 2213.85 g/mol
Observed m/z : 2213.83 g/mol
[M]+
2213.83
m/z

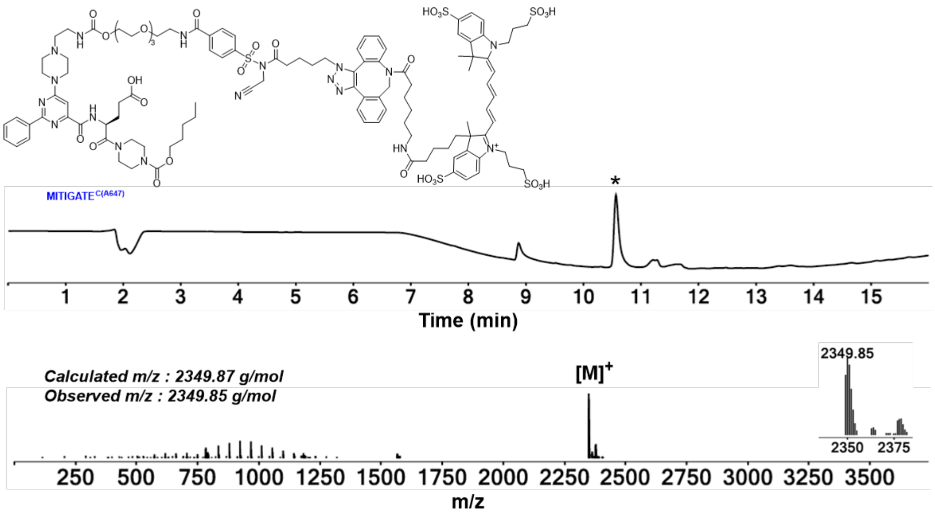
MITIGATE^C(A647)
Time (min)
Calculated m/z : 2349.87 g/mol
Observed m/z : 2349.85 g/mol
[M]+
2349.85
m/z

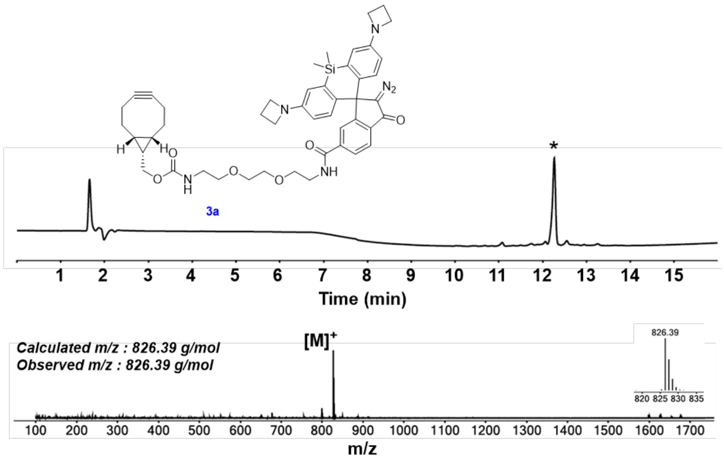
3a
Time (min)
Calculated m/z : 826.39 g/mol
Observed m/z : 826.39 g/mol
[M]+
826.39
m/z

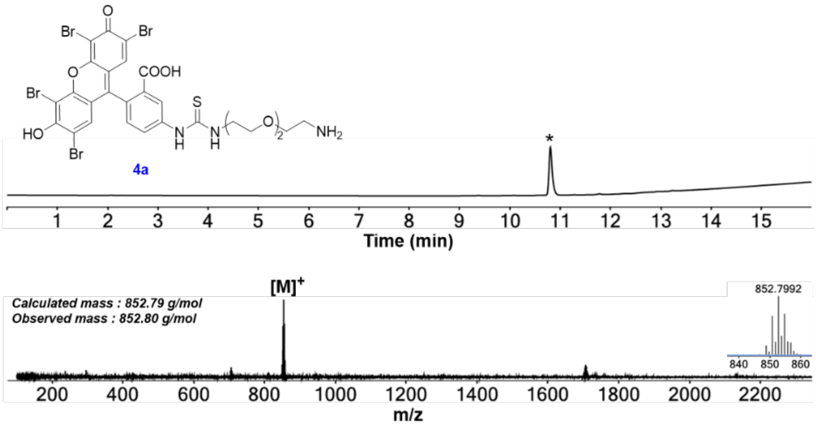
4a
Time (min)
Calculated mass : 852.79 g/mol
Observed mass : 852.80 g/mol
[M]+
852.7992
m/z

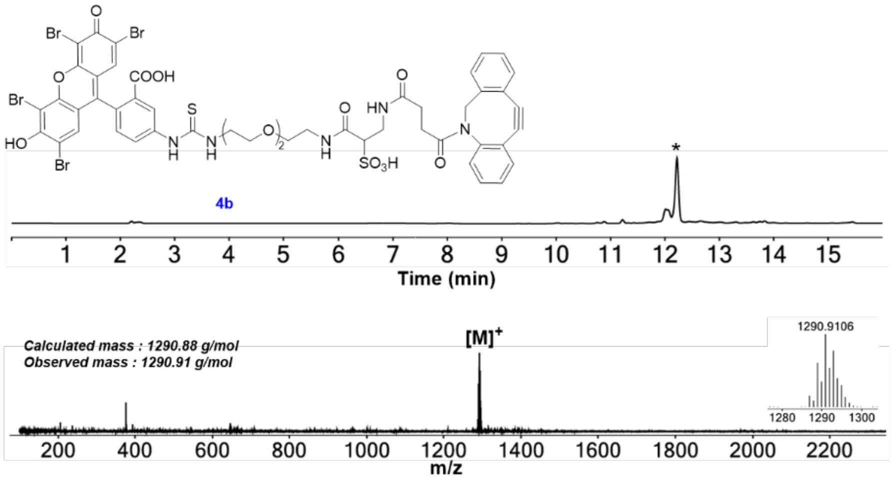
4b
Time (min)
Calculated mass : 1290.88 g/mol
Observed mass : 1290.91 g/mol
[M]+
1290.9106
m/z

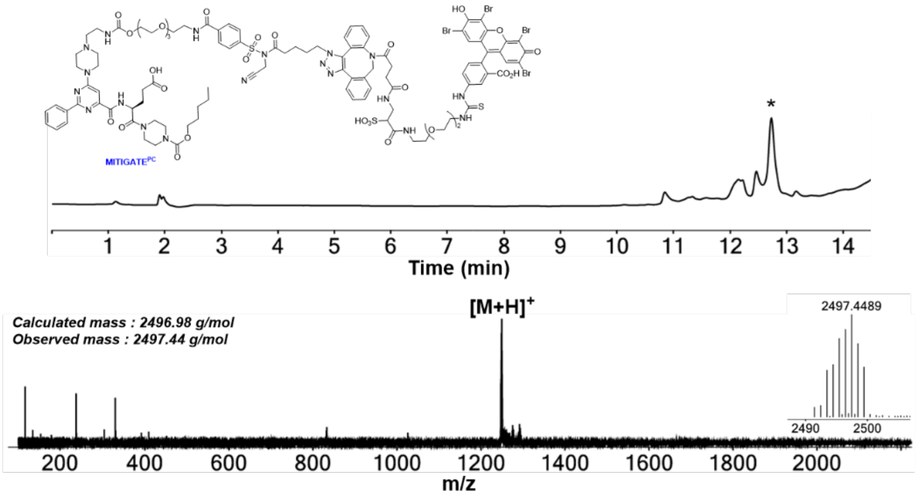
